# Detection of fusion transcripts and their genomic breakpoints from RNA sequencing data

**DOI:** 10.1101/2021.05.17.441778

**Authors:** Youri Hoogstrate, Malgorzata A. Komor, René Böttcher, Job van Riet, Harmen J. G. van de Werken, Stef van Lieshout, Ralf Hoffmann, Evert van den Broek, Anne S. Bolijn, Natasja Dits, Daoud Sie, David van der Meer, Floor Pepers, Chris H. Bangma, Geert J. L. H. van Leenders, Marcel Smid, Pim French, John W.M. Martens, Wilbert van Workum, Peter J. van der Spek, Bart Janssen, Eric Caldenhoven, Christian Rausch, Mark de Jong, Andrew P. Stubbs, Gerrit A. Meijer, Remond J.A. Fijneman, Guido Jenster

## Abstract

Spliced fusion-transcripts are typically identified by RNA-seq without elucidating the causal genomic breakpoints. However, non poly(A)-enriched RNA-seq contains large proportions of intronic reads spanning also genomic breakpoints. Using 1.274 RNA-seq samples, we investigated what additional information is embedded in non poly(A)-enriched RNA-seq data. Here, we present our novel, graph-based, Dr. Disco algorithm that makes use of both intronic and exonic RNA-seq reads to identify not only fusion transcripts but also genomic breakpoints in gene but also in intergenic regions. Dr. Disco identified TMPRSS2-ERG fusions with genomic breakpoints and other transcribed rearrangements from multiple RNA-sequencing cohorts. In breast cancer and glioma samples Dr. Disco identified rearrangement hotspots near CCND1 and MDM2 and could directly associate this with increased expression. A comparison with matched DNA-sequencing revealed that most genomic breakpoints are not, or minimally, transcribed while also revealing highly expressed translocations missed by DNA-seq. By using the full potential of non poly(A)-enriched RNA-seq data, Dr. Disco can reliably identify expressed genomic breakpoints and their transcriptional effects.

## Introduction

Genomic rearrangements are frequently observed in cancer and these can drive disease initiation and progression through disruption of tumour suppressor genes and activation of oncogenes ^1–3^. Marked examples include TMPRSS2-ERG fusions in prostate adenocarcinoma ^4^ and *BCR*-*ABL* in chronic myelogenous leukaemia ^5^. DNA breakpoints and their aberrant ligations are identified by whole genome sequencing (WGS) but their potential role as driver mutations is mostly unresolved as-of-yet. The majority of DNA breakpoints involve intergenic and intronic regions and thus not part of messenger RNA (mRNA) and protein coding sequences ^6^ and genomic breakpoints of fusion genes are mostly located intronic ^7^. To reveal their downstream effects, RNA-sequencing (RNA-seq) is crucial to investigate changes at the transcriptional level and identify actual (in-frame) fusion transcripts. Conversely, for fusion-transcripts, identification of the exact genomic breakpoint(s) can be essential to explain changes in gene expression and to define the origins of alternative promoter usage and altered splicing or polyadenylation events. Combined genomic and expression data allows to further study functional consequences of genomic rearrangements and signifies whether the event is merely a passenger or a putative driver mutation ^7,8^. However, for many transcriptome studies, the exact genomic breakpoints of expressed rearrangements have not been resolved as matched WGS, Sanger sequencing, or similar analyses were not performed. Therefore, we set out to determine whether exact genomic breakpoints could be identified from RNA-seq data.

Next to targeted gene approaches, there are two main approaches in preparing RNA-seq libraries ^9^. First, the more traditional method includes the positive selection of poly-adenylated messenger RNA (mRNA; poly(A)^+^) to specifically target mRNA and eliminate abundant ribosomal RNA (rRNA). Alternatively, one may extract total RNA and use random hexamer primers to initiate cDNA synthesis while removing abundant unwanted RNAs by various additional methods. This approach is referred to as rRNA-minus and is commonly applied when (partially) degraded RNA from formalin-fixed paraffin-embedded (FFPE) samples are sequenced.

rRNA-minus RNA-seq is thus capable of identifying non-poly(A) transcripts such as circRNAs, specific types of small and long non-coding RNAs and actively-transcribed precursor mRNAs (pre-mRNAs) ^10^. Although the exact numbers depend on the used protocol, tissue type, lariats ^11^ and intron lengths, typically 30-40% of rRNA-minus RNA-seq reads map to intronic features compared with 5-10% in poly(A)^+^ RNA-seq ^12^. Therefore, rRNA-minus RNA-seq datasets require at least 50% higher sequencing depth to achieve a similar exon coverage comparable to poly(A)^+^ RNA-seq, while being capable of identifying additional RNA classes ^9^.

Fusion genes such as *TMPRSS2*-*ERG* and *BCL*-*ABL* are frequently observed as drivers within their respective malignant tissue ^13^. Yet, many observed fusion genes are still of unknown consequence and seen in small frequencies in various cancer types. RNA-seq is highly suitable for fusion gene detection ^14–16^. Methods to integrate RNA and DNA fusions and breakpoints allow to further assess functional consequences ^7,8,17^, and are even capable of integrating higher order complex rearrangements, but remain dependent on the availability of matching DNA data. Current fusion-detection tools such as FusionMap, FusionCatcher and JAFFA focus on exon regions or splice junctions ^18–20^ which are the main target of poly(A)^+^ RNA-seq. Indeed, these tools also work well on rRNA-minus RNA-seq as these also include exonic reads. This efficient search space reduction in classical fusion gene detectors in turn reduces the overall complexity and processing time. However, using rRNA-minus RNA-seq, typically 30-40% of the aligned reads are intronic and a further 20-25% of all reads are found to be intergenic ^12^, which are often a priori neglected. This large number of intronic and intergenic reads provides an opportunity to identify additional cancer specific transcripts and the exact genomic breakpoints of fusion genes. We have previously shown in a proof-of-concept that rRNA-minus RNA-seq can identify genomic breakpoints ^10^. Here, we present an algorithm named Dr. Disco (https://github.com/yhoogstrate/dr-disco) which computationally identifies such genomic breakpoints and exon-to-exon junctions in a genome-wide fashion, taking into account the potential of rRNA-minus RNA-seq. We applied Dr. Disco on five large RNA-seq datasets spanning multiple malignant tissues (n=1.274) (**Table 1**). Indeed, we reveal exact causal genomic breakpoints as derived from RNA-sequencing alone but limited to regions sufficiently expressed. Furthermore, we show that rRNA-minus RNA-seq data can reveal more transcriptionally active rearrangements than poly(A)^+^ RNA-seq and therefore is a useful analysis to supplement WGS. Thus, rRNA-minus RNA-seq in combination with a suited analysis pipeline gives a more complete view on both the origin and effects of genomic rearrangements and their direct influence on the expression of associated genes.

**Table 1.**
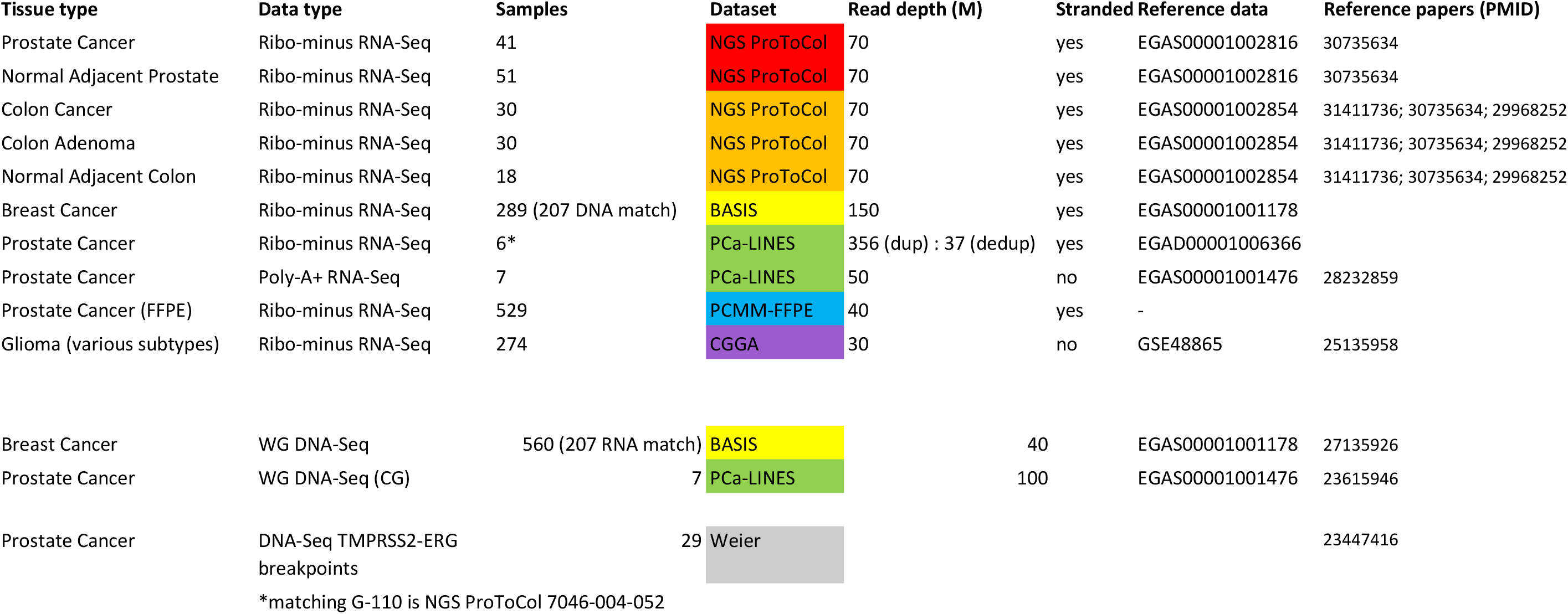

## Results

To identify exact genomic breakpoints from rRNA-minus RNA-seq, we developed a novel algorithm and implemented this in Python, termed Dr. Disco. The tool uses discordant reads ^21^, reads with a split alignment or read pairs with an inverted or large insert size. The method uses reads from not only exonic but also intronic and intergenic regions (**Figure 1** and **Supplementary Dr. Disco technical specification**). These split and spanning reads are converted and inserted into a breakpoint graph ^7^. The graph is analysed to find reads originating from the same junctions.

**Figure 1.**
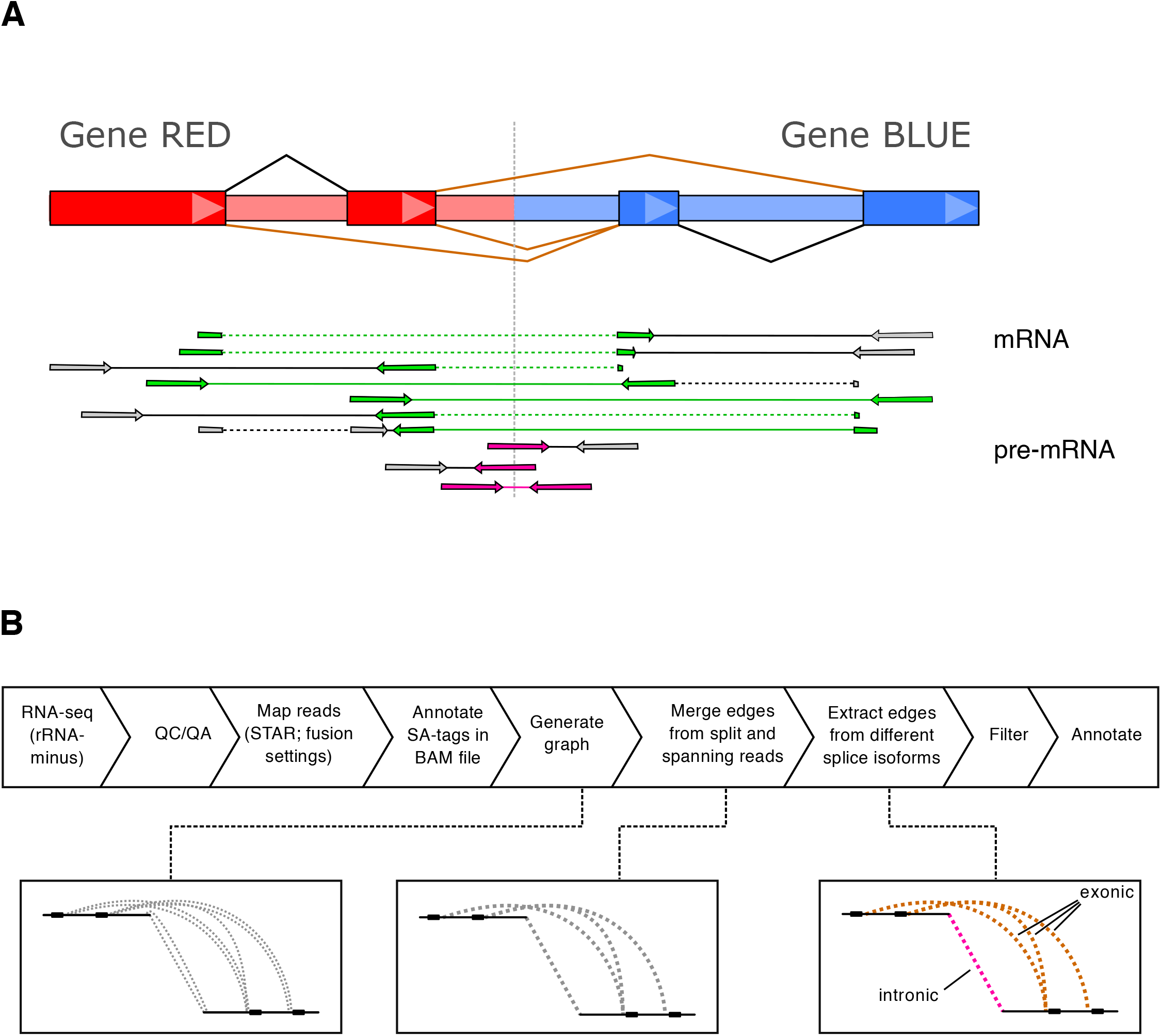
Overview intronic RNA and Dr. Disco algorithm. (A) Schematic representation of fusion-gene RED-BLUE. Due to relatively large intron sizes, in-gene genomic breakpoints are expected to occur most often intronic. The fusion could result in different isoforms of mature mRNA as indicated with fusion splice junctions (brown). Fusion splice junction spanning reads form the classical source of evidence for detecting mature mRNA fusion-events. In rRNA-minus data, intronic pre-mRNA reads (pink) may cover the causal genomic breakpoints. (B) Flowchart of the Dr. Disco pipeline. RNA-seq data is aligned to obtain discordant aligned reads; reads are transformed into edges that are inserted into a graph. In the graph, edges corresponding to either intronic or exonic junctions are kept separate. Detection of junctions is performed by analysing the graph for clusters. An additional splice variant correction is applied. Each identified junction variant is marked intronic or exonic and then filtered and annotated.

For terminology, we define exon-to-exon splice junctions as junctions of which it may be expected that they could be detected by classical fusion detection algorithms. Fusion transcripts which are not a result of (cryptic-)exon-to-exon splicing are typically intron-to-intron junctions located exactly at genomic breakpoints. In addition, it is possible that genomic breakpoints are located within exons and do not result in fused spliced junctions (**Figure S1**). Because these junctions do not match splice junctions and are not the primary target of classical fusion gene detection, we also consider them intronic. Corresponding detected junctions are marked exonic or intronic accordingly. The detailed computational methodology is described in **Supplementary Methods** and the **Supplementary Dr. Disco technical specification**.

### Comparison poly(A)^+^ and rRNA-minus RNA-seq

To determine the overlap of genomic breakpoints as detected from DNA-seq with those detected from RNA-seq using Dr. Disco, seven prostate cancer (PCa) samples (PCa-LINES dataset) were sequenced using the Complete Genomics WGS platform and with matching poly(A)^+^ and rRNA-minus RNA-seq. After filtering out the exon-exon junctions, we found that rRNA-minus RNA-seq identified more (3.4 times) intronic junctions, thus predicted genomic breakpoints, between chromosomes as compared to poly(A)^+^ RNA-seq (**Figure 2A**). Although poly(A)^+^ RNA-seq also harbours genomic breakpoints, they are less confidently called as they have fewer read counts and mostly lie in 3’ UTR terminal exons as in-exon located genomic breakpoints (**Figure S2**). Terminal exons are known for their relatively large size as they are approximately 6-7 times larger than internal exons ^22^. The number of exonic junctions, thus predicted mRNA fusions, identified by Dr. Disco is nearly identical for rRNA-minus and poly(A)^+^ RNA-seq (144 vs 155). Of the exonic junctions detected in rRNA-minus samples, 52% were also found in the poly(A)^+^ data. However, another 26% also matched the poly(A)^+^ data but did not pass filtering, mostly because of insufficient discordant reads.

**Figure 2.**
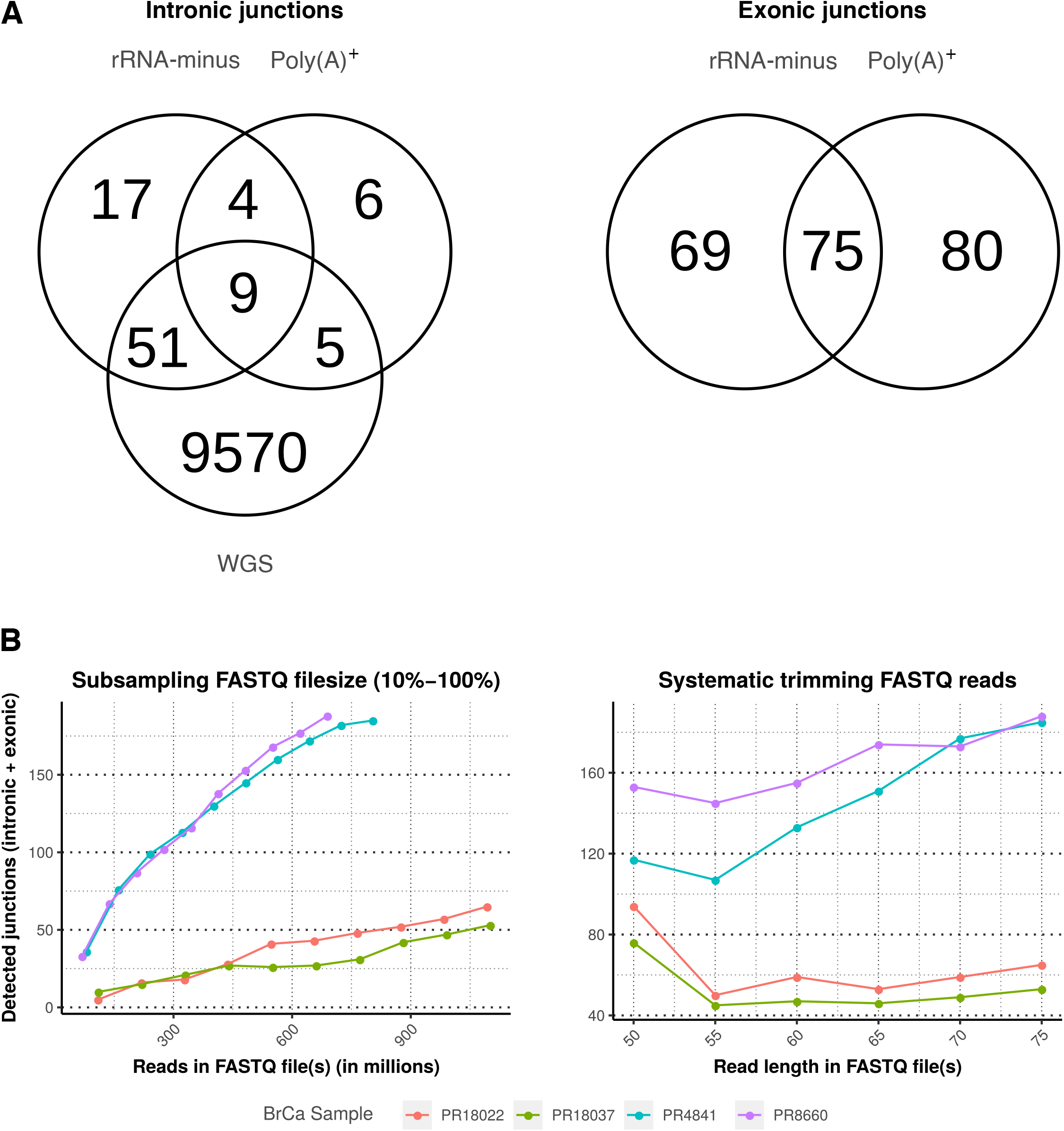
Overlap across sequencing types and library size influence. (A) Venn diagram with overlap of cumulative interchromosomal junctions of 7 WGS PCa samples rRNA-minus and poly(A)+ RNA-seq (PCa-LINES dataset). Overlap in only intronic junctions representing genomic breakpoints (left) and only exonic splice junctions (right). Of the 69 exonic junctions only found in rRNA-minus RNA-seq, 40 were detected in the matching poly(A)+ but did not pass filtering. Of the 80 poly(A)+-only exonic junctions, 58 were found in rRNA-minus but did not pass filtering. (B) The number of predicted junctions as function of sequencing depth (left) and read-length (right) reduction. BrCa samples were selected for high sequencing depth (PR18022 & PR18037) or a high number of junctions (PR4841 & PR8660). Left: The number of predicted junctions per sequencing-depth (10-100%) with the full read-length (2×75 bp). Reducing the sequencing depth, also for samples with a high sequencing depth, reduces the number of detected junctions. Only sample PR4841 reaches a plateau. Right: Each data point represents the number of predicted junctions per given read-length, with full sequencing-depth. Truncating sequencing-reads results in a lower number of predicted junctions. However, below 55 nucleotides this number of increases.

### Comparison of RNA-with DNA-seq data

Within these 7 PCa samples, the number of genomic breakpoints identified in WGS vastly outnumbered those extracted from the rRNA-minus RNA-seq (6.8%), indicating that only a fraction of the genomic rearrangements is expressed at a level to be detected by rRNA-minus RNA-seq. The intronic and exonic junctions as detected by rRNA-minus RNA-seq show overlap with the genomic breakpoints detected by WGS (**Figure S3**). Interestingly, 27 interchromosomal genomic breakpoints were only found by RNA-seq; 6 genomic breakpoints by poly(A)^+^ only, 17 by rRNA-minus only and 4 by both RNA sequencing methods (**Table S1**).

To identify the influence of sequencing coverage and read length on the number of intronic and exonic junctions, 4 breast cancer (BrCa) RNA-seq samples from the BASIS dataset ^23,24^ were systematically truncated (**Figure 2B**). The number of detected junctions dropped as sequencing reads became shorter, showing that a minimum length of 55 bases is necessary for accurate detection using Dr. Disco. We noticed an increase in discordantly-aligned reads when they were truncated to 50bp. This was due to an overall increase in misalignments that do not resemble actual evidence of genomic rearrangements. Irrespective of the number of genomic breakpoints present within a sample as determined by WGS, an increase in overall sequencing depth is positively correlated with an increase in detected junctions (**Figure 2B**).

Genomic breakpoints detected by WGS from 207 BrCa samples from the BASIS dataset ^23,24^ were compared to their matching rRNA-minus RNA-seq detected junctions. Only interchromosomal entries were compared to avoid fusion transcripts unrelated to genomic rearrangements such as read-throughs or circRNAs. WGS identified a total of 6531 interchromosomal genomic breakpoints and, similar to the seven prostate cancer samples, the majority of the genomic breakpoints were not detectable in the matching RNA-seq. Only 409 events (6.3%) were found in both assays, a similar percentage compared with our analysis on PCa samples (**Figure 3A**). Dr. Disco detected 377 unique genomic breakpoints (48%) which were only present within the RNA-seq data, of which 109 of these genomic breakpoints were identified within eight BrCa samples which also had an overall high number of (WGS-detected) genomic breakpoints (**Figure S4**). The density of WGS and RNA-seq detected junctions was highly similar (R^2^=0.72, **Figure 3B**, BrCa plots; **Figures S5-S7**), with prominent focal peaks near the genomic locus of *CCND1*, *SHANK2* and *FGFR1*.

**Figure 3.**
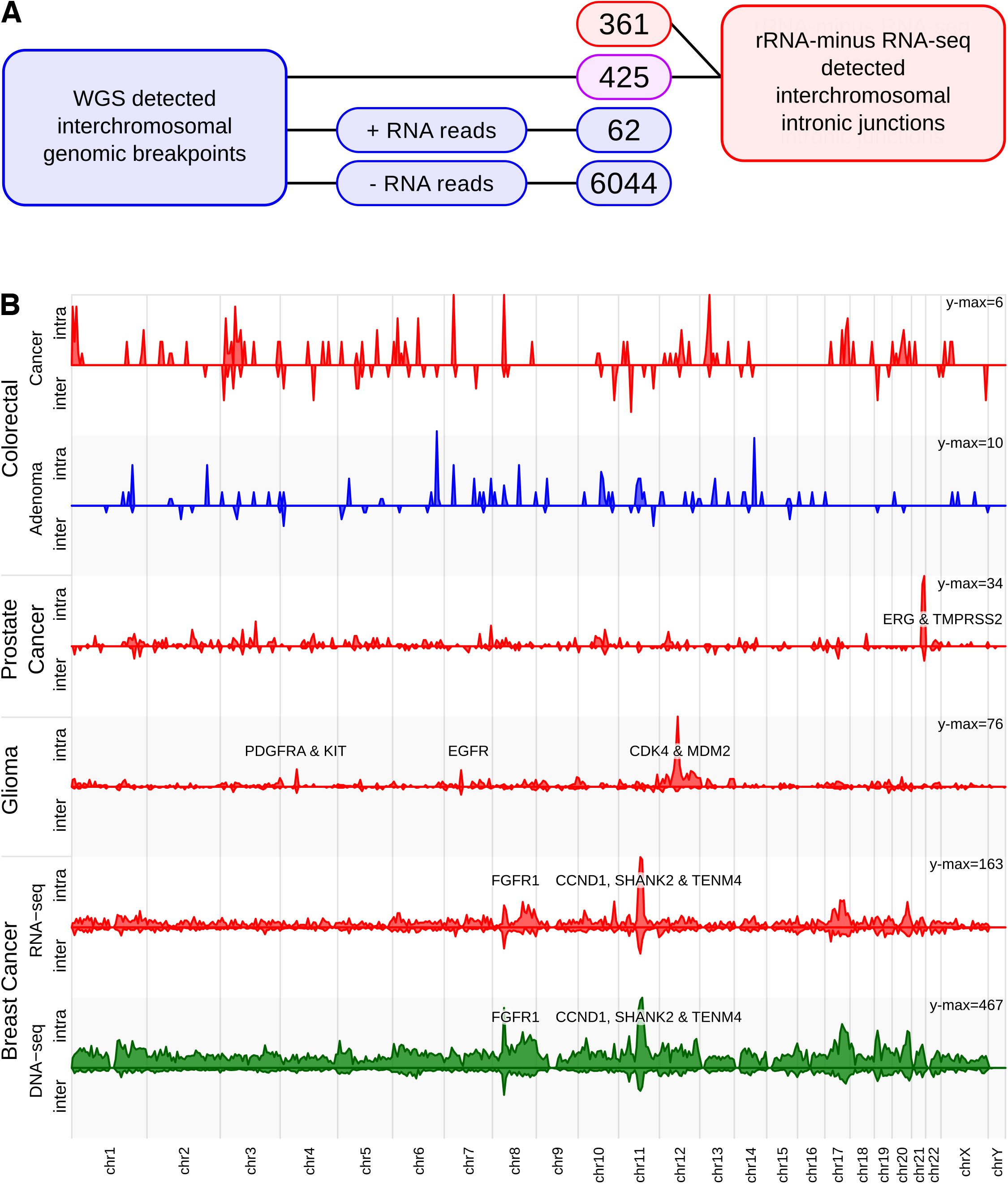
Integration RNA-seq analysis with WGS results. (A) Number of detected genomic breakpoints per subgroup in WGS and rRNA-minus RNA-seq data of 207 matching BrCa samples. Rectangles in blue indicate presence only in WGS data, in red only in RNA-seq data and in pink in both. To avoid artifacts from RNA post-processing such as circRNAs and read-throughs, only interchromosomal entries were interrogated. Of the interchromosomal WGS breakpoints, 6059 did not have sufficient discordant reads in the RNA-seq data. Of 62 genomic breakpoints, the threshold of sufficient discordant RNA-seq reads was exceeded, but it was not detected by Dr. Disco or did not pass filtering. 425 breakpoints were detected in both the assays and 361 RNA-seq detected breakpoints did not match a WGS entry. (B) Chromosome plot representing the density of inter and intrachromosomal genomic breakpoints. For the BrCa samples, Dr. Disco RNA-seq analysis (red) and WGS breakpoints (green) are depicted. The number of RNA-seq genomic breakpoints in the colorectal cancer and adenomas is low and no recurrent breakpoints were identified yet. The number of genomic breakpoints in colorectal adenomas was lower than in colorectal cancer. The observed peaks in colorectal cancer originated from multiple, sample specific, junctions (**Figure S9**).

### Pan-cancer analysis

We analysed rRNA-minus RNA-seq data (n=651) from different malignant tissues and datasets using the Dr. Disco algorithm (**Figures 3B** & **4**). This included the earlier described BrCa dataset BASIS (n=207), NGS-ProToCol (normal adjacent prostate; n=41, prostate cancer; n=51; normal adjacent colon; n=18, colorectal adenoma; n=30 and colorectal carcinoma; n=30) and the Chinese glioma atlas (CGGA) (various glioma types; n=274) (**Table 1**).

Intronic and exonic junctions were identified in each dataset. The different malignant tissue types showed distinct regions enriched with intronic and exonic junctions, as represented in a chromosome plot (**Figure 3B**). Known prominent events include *TMPRSS2*-*ERG* (chr21) in PCa, *EGFR* (chr7) in glioma and *CCND1* rearrangements (chr11) in BrCa. The number of breakpoints per sample with associated clinical parameters is provided in **Figure 4**. The lowest average number of genomic breakpoints per tissue type was found in normal adjacent samples (colon=0.5; prostate=0.9) followed by colorectal adenoma (1.1) (**Figures S8-S9**). The *TMPRSS2*-*ERG* fusion-event was observed in two normal adjacent prostate samples containing genomic breakpoints exactly identical to their matching malignant sample and were therefore contaminated with cancer cells (**Figure S8B**). Of the different malignant tissue types, colorectal cancer samples were characterized by the lowest average number of junctions (1.1) followed by combined low- and high-grade glioma (2.1) (**Figure S10**). Conversely, PCa (4.3) and BrCa (9.3) were characterized by relatively high numbers of genomic breakpoints per sample. Absolute numbers were used since not only sequencing depth but also read depth and library preparation differ per dataset.

**Figure 4.**
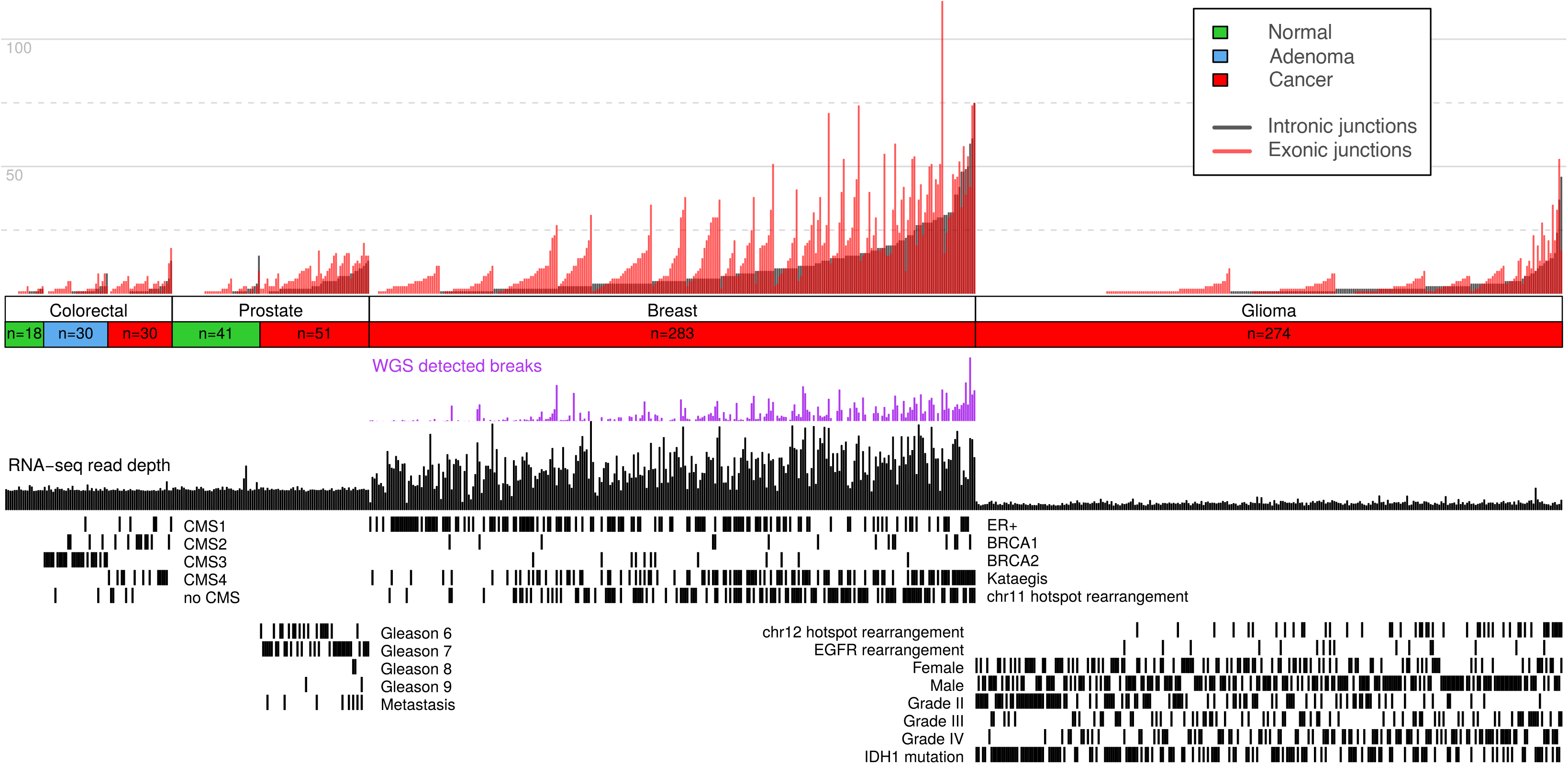
Results summary. Intronic and exonic junctions are given per sample for the NGS-ProToCol, BASIS and CGGA datasets with their associated clinical parameters. For the colon samples, the predicted CMS classes are provided, for the prostate cancer samples the Gleason grade and metastatic progression are provided, for the breast cancer samples the ER, BRCA1, BRCA2, kataegis and Dr. Disco detected chr11-hotspot status are provided and for the glioma samples the grading, recurrence, IDH1 mutation status, gender and the Dr. Disco detected EGFR and chr12 hotspot status are provided.

Several clinical parameters were associated with the number of Dr. Disco-detected genomic breakpoints per sample (**Figure 4**). In BrCa, kataegis (p=1.9e^−09^) was positively associated with the number of observed genomic breakpoints whereas ER+ BrCa revealed to be negatively associated (p=0.9e^−03^) with the number of genomic breakpoints. In glioma, tumour grade IV is positively associated with the number of genomic breakpoints per sample (p=1.1e^−05^), whereas tumour grade II (p=2.9e^−08^) and presence of an IDH1 mutation (p=0.8e^−03^) were negatively associated. The number of intronic junctions detected by Dr. Disco on RNA-seq correlates positively with the number of WGS-detected genomic breakpoints within BrCa (ρ=0.71, p=2.2e^−16^, **Figure S11**). Although trends within PCa were observed for the incidence of high Gleason grade (>=8; p=0.08; n=4/50) and metastasis (p=0.16; n=8/51) associated with the number of genomic breakpoints, it did not reach statistical significance. Because of the relative low number of genomic breakpoints per sample and the rather low number of colorectal cancer samples, further in-depth analysis on recurrent events could not be performed.

In the BASIS and NGS-ProToCol datasets approximately 65% of all intronic and exonic junctions have both sides located within an annotated gene (**Figure 5**). Thus, approximately 35% of these junctions have at least one side located within an intergenic region, regions that are often dismissed *a priori* by classical fusion gene detection tools ^19,20^. We found transcripts with incorporated cryptic (unannotated) exons. For instance, a BrCa sample harboured intergenic junctions in *SDC4* transcripts using 5 consecutive cryptic exons (**Figure S12**). In contrast, a PCa sample had an intergenic rearrangement lacking mRNA level transcripts, thus only visible by the presence of pre-mRNA (**FigureS13**).

**Figure 5.**
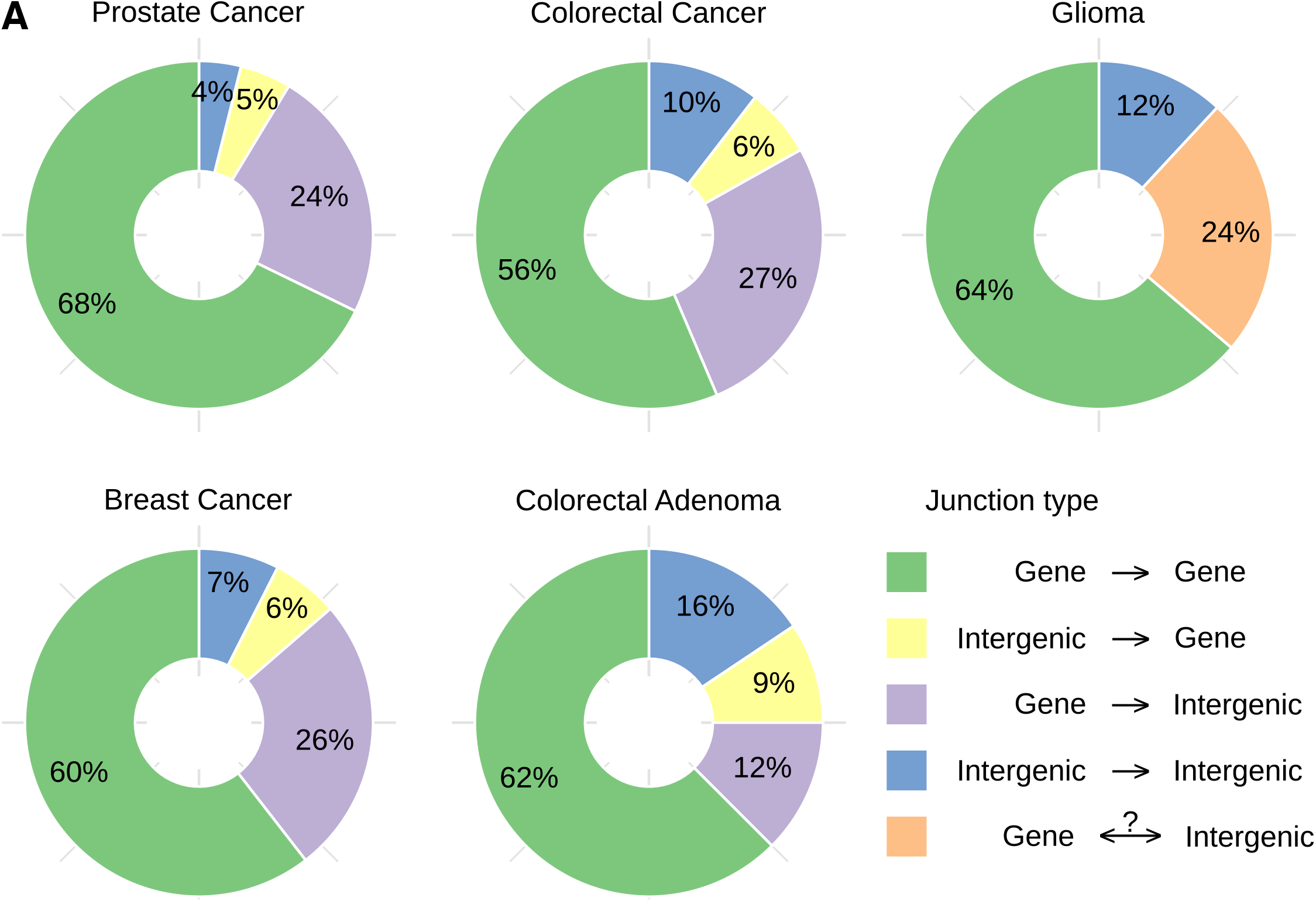
Genic/Intergenic junction status. (A) The frequency of intronic and exonic junctions with one or both breakpoints in gene and/or intergenic regions (Ensembl 89). Because the glioma dataset was sequenced unstranded, junctions with one intergenic side are grouped together. In all datasets, approximately 3/8 of the junctions have at least one intergenic side. Both inter- and intrachromosomal junctions were included, suspected circRNAs were discarded; unlocalized and unplaced sequences (chrUn_…) and alternate loci (chr…_alt) were discarded. Matching intronic and exonic predictions of the same variant were treated as a single entry.

### Genes associated with peaks in breakpoints

There were multiple, cancer type-specific, hotspots of junctions located near known oncogenes (**Figure 3**) such as *KIT*, *PDGFRA*, *EGFR*, *CDK4*, *MDM2* (glioma), *TMPRSS2*, *ERG* (PCa), *FGFR1* and *CCDN1* (BrCa). Recurrent gene fusions are depicted in **Table S2** and the list of all identified junctions is provided in an online data repository (**Table S3**; doi:10.5281/zenodo.4159414). Enrichment analysis was performed using HUGO symbols of genes recurrently hit per cohort, indicating the pathway “*Transcriptional misregulation in cancer [KEGG:05202]*” is significantly more frequently hit (p=1.6e^−04^) within PCa due to *TMPRSS2*, *ERG*, *ETV1*, *H3FA3*, *SLC45A3* and *ELK4*. Within BrCa, pathways *ETF* and *E2F* are significantly enriched (p=6.75e^−10^, p=2.8e^−06^) in ER+ BrCa and “*Proteoglycans in cancer*” in ER-BrCa (p=1.4e^−05^). Genes that are recurrently hit in glioma samples were found more often in pathways “*Rap1 signaling pathway*” (p=3.2e^−04^), “*Glioma*” (p=5.9e^−03^) and “*Ras Signaling*” (p=2.6e^−03^) (**Table S4**).

### TMPRSS2-ERG

In 32 of the 51 NGS-ProToCol PCa samples Dr. Disco detected the mRNA fusion-transcripts of *TMPRSS2*-*ERG* fusions, including a genomic breakpoint in 27/32 samples (**Figure 6**). These fusions were in concordance with high *ERG* expression in those samples only. The detection rate for genomic breakpoints for this oncogenenic fusion gene is thus markedly higher than for the overall number of genomic breakpoints. The genomic breakpoint did not pass filtering in sample 072, was marked exonic in sample 027 and was merged with closely adjacent (<450 bp; insert size) exonic junctions in three samples (053, 050 and 065); indicating that breakpoint-spanning reads were present in all 32 samples. Three other samples (075, 054 and 048) had their *ERG*-flanking genomic breakpoint located in an intergenic region upstream to *ERG*’s first exon (**Figures S14** & **S16**). In these samples, cryptic exons were identified in *TMPRSS2*-*ERG* fusion mRNA transcripts (chr21:38,692,521-38,692,797 and chr21:38,701,593-38,701,947; hg38). Two of the three samples with their breakpoint before *ERG* had additional deletions in *ERG*, removing exon 2. The most abundant intronic junctions were T1-E4 and T1-E5 (**Figures 6** & **S15-S16**) which is in concordance with previous reports ^25^. Genomic breakpoints were indeed located in hotspot-regions within the first two introns of *TMPRSS2* and the last half of *ERG* intron 3 ^26^. Additional shallow sequenced FFPE RNA-seq samples which were subsequently analysed by Dr. Disco revealed the *TMPRSS2*-*ERG* fusion in 181 samples (**Figures S15-S16**) and confirmed this remarkable breakpoint preference region within *ERG* intron 3 more precisely.

**Figure 6.**
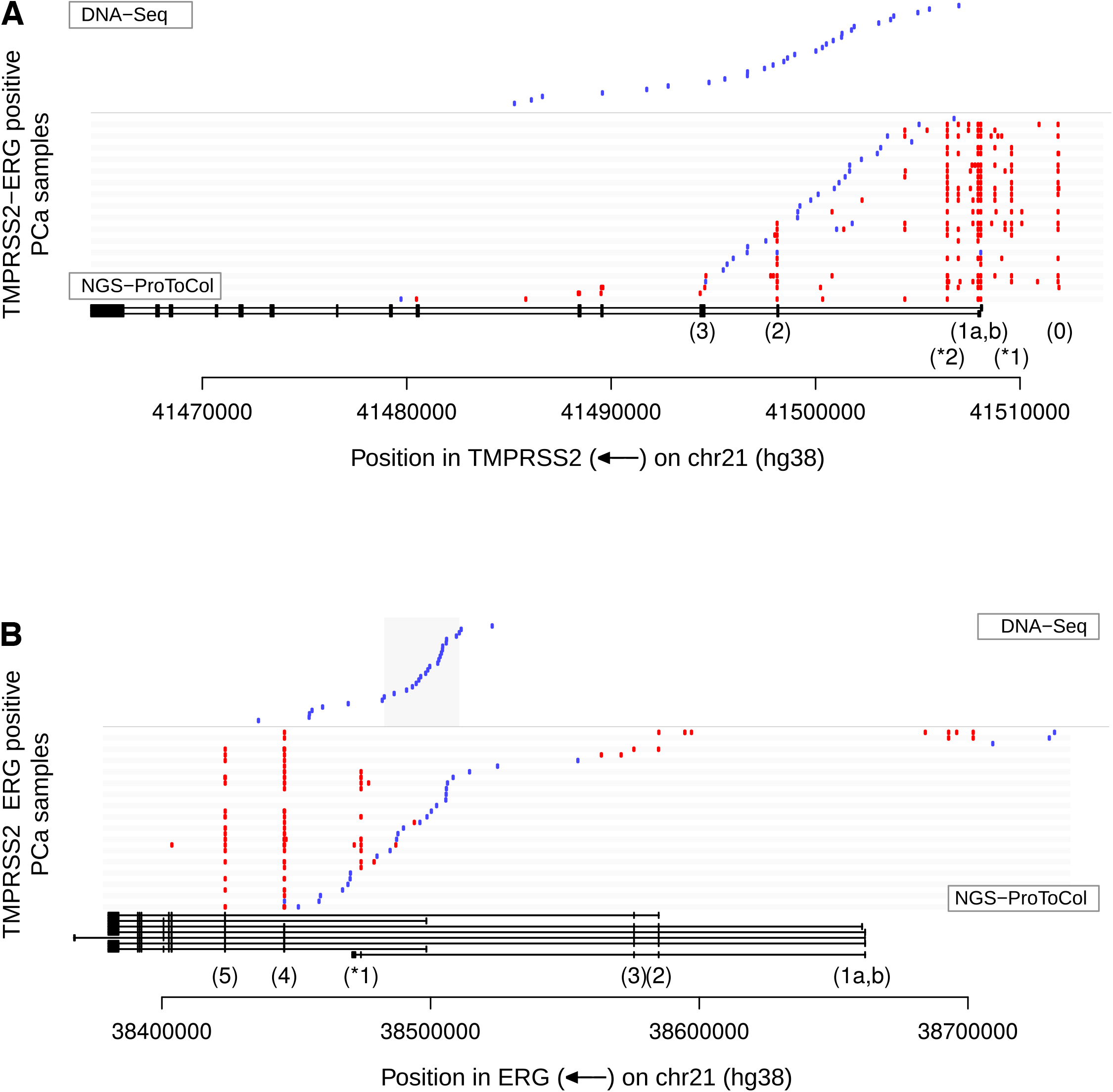
TMPRSS2-ERG junction map. Summary of TMPRSS2 and ERG junctions and breakpoints in NGS-ProToCol RNA-seq and non-matching targeted DNA-seq (Weier dataset). Gene structures are indicated at the bottom. Intronic Dr. Disco detected junctions (representing genomic breakpoints) and genomic breakpoints from the Weier dataset are indicated in blue and exonic junctions in red. (A) For TMPRSS2, most breakpoints are detected after exon 1, up to exon 3. At mRNA level, apart from the first exons (1a and 1b), exon 0 and exon 2 were commonly included in fusion transcripts. Also, two novel recurrent cryptic exons (*1 and *2) were often observed in fusion transcripts. (B) In ERG we observe in the NGS-ProToCol data three samples (048, 054 & 075) that have their genomic breakpoints before ERG and result in transcripts with additional, novel, intergenic cryptic exons.

PCa cell line VCaP is known to have *TMPRSS2*-*ERG* with two additional rearrangements ^26,27^. We interrogated the fusion in VCaP using Dr. Disco on both rRNA-minus and poly(A)^+^ RNA-seq data. Poly(A)^+^ RNA-seq shows that only the first exon of *TMPRSS2* splices to *ERG*, even though the genomic breakpoint to *ERG* is located in the 5^th^ intron (**Figure S17A**). The rRNA-minus data confirms this splice junction but also reveals all the other genomic breakpoints spanning *TMPRSS2* and *ERG*. Read stranding indicates that a region containing the 4^th^ and 5^th^ exon is inverted, and that its breakpoint-A is an inversion. Breakpoint-B is an amplification and the junction from TMPRSS2 to *ERG* is again inverted such that *ERG* is in its original orientation, which deletes the genomic region containing exons 2 and 3 of *ERG*. Thus, only *TMPRSS2* exon 1 splices to *ERG* since exon 2 and 3 are deleted and exon 4 and 5 are inverted (**Figure S17B**). The small proportion of reads within the deleted *TMPRSS2* exons 2 and 3 in the rRNA-minus data originated from the non-fusion allele(s). The rRNA-minus RNA-seq data not only revealed both intronic and exonic junctions but also clarifies the complex downstream effects on transcription. As expected, analysing the rRNA-minus RNA-seq data with FusionCatcher^15^ resulted only in the exonic *TMPRSS2*-*ERG* junction, similar to the Dr. Disco results in poly(A)^+^ RNA-seqdata.

Other PCa-related and detected *TMPRSS2* fusions were *TMPRSS2*-*RERE*, *SERINC5*-*TMPRSS2*, *TMPRSS2*-*TBX3*, *TMPRSS2*-*PADI4*, *MGA*-*TMPRSS2* and *TMPRSS2*-*CATSPER2* (**Table S5**). Two novel exons in *TMPRSS2* were observed in both fusion and wild-type transcripts (**Figure 6**). These cryptic exons were both lowly expressed as they represented ~3% of all *TMPRSS2*-*ERG* reads in samples having the splice variant. Additionally, intergenic TMPRSS2 exon-0 ^28^ was detected by Dr. Disco in fusion mRNA-transcripts within 18/32 *TMPRSS2*-*ERG* positive samples.

In one sample we identified an exonic junction originating in *ERG* and spanning to *TMPRSS2* in which the gene order and included exons indicated that this *ERG*-*TMPRSS2* fusion was caused by a reciprocal translocation instead of the common 3 Mb deletion (**Figure S18**).

### Large gene amplifications

Hotspot regions (20-30 Mb) enriched with RNA-seq detected breakpoints were observed in the BrCa (chr11) and glioma (chr12) datasets. These hotspots differ from focal events (e.g.*TMPRSS2*-*ERG*) in the sense that they were larger, had no consistent fusion-partners and often contained multiple hotspot junctions per sample. To understand their function and what triggers their selective advantage, the transcriptional effects of these rearrangements were investigated by performing differential gene expression analysis between BrCa and glioma samples with and without a chr11 and chr12 hotspot rearrangement (BrCa: n=122/283; glioma: n=45/274, respectively). BrCa samples having a chr11 hotspot rearrangement were characterized by a large stretch of significant up-regulated genes within the respective hotspot region (**Figures 7A-C** & **S19**). The large genes SHANK2 and TENM4, both located in the hotspot region, were the most frequently hit genes (25 and 13 BrCa samples, respectively), yet were not among the strongest up-regulated genes of the overall region. Instead, genes with an extreme fold-change were *FGF4* and *CCND1*, the cluster *KCTD21*, *ALG8* & *GAB2* and genes downstream of *TENM4*. Up-regulation of the overall region indicates amplifications of *CCND1* and/or the gene cluster, which is in concordance with previous reports ^29^. The high frequency of junctions in the relatively large, yet not heavily upregulated *SHANK2* (785 kb) and *TENM4* (788 kb), suggests they are ‘collateral damage’ of the amplifications; a hypothesis that has been described in glioma previously ^30^. This hypothesis is further supported by the lack of consistent fusion partners, consistency in acting as acceptor or donor and the absence of a clear spike in cumulative breakpoints (**Figure 7A-B**; **TableS6**).

**Figure 7.**
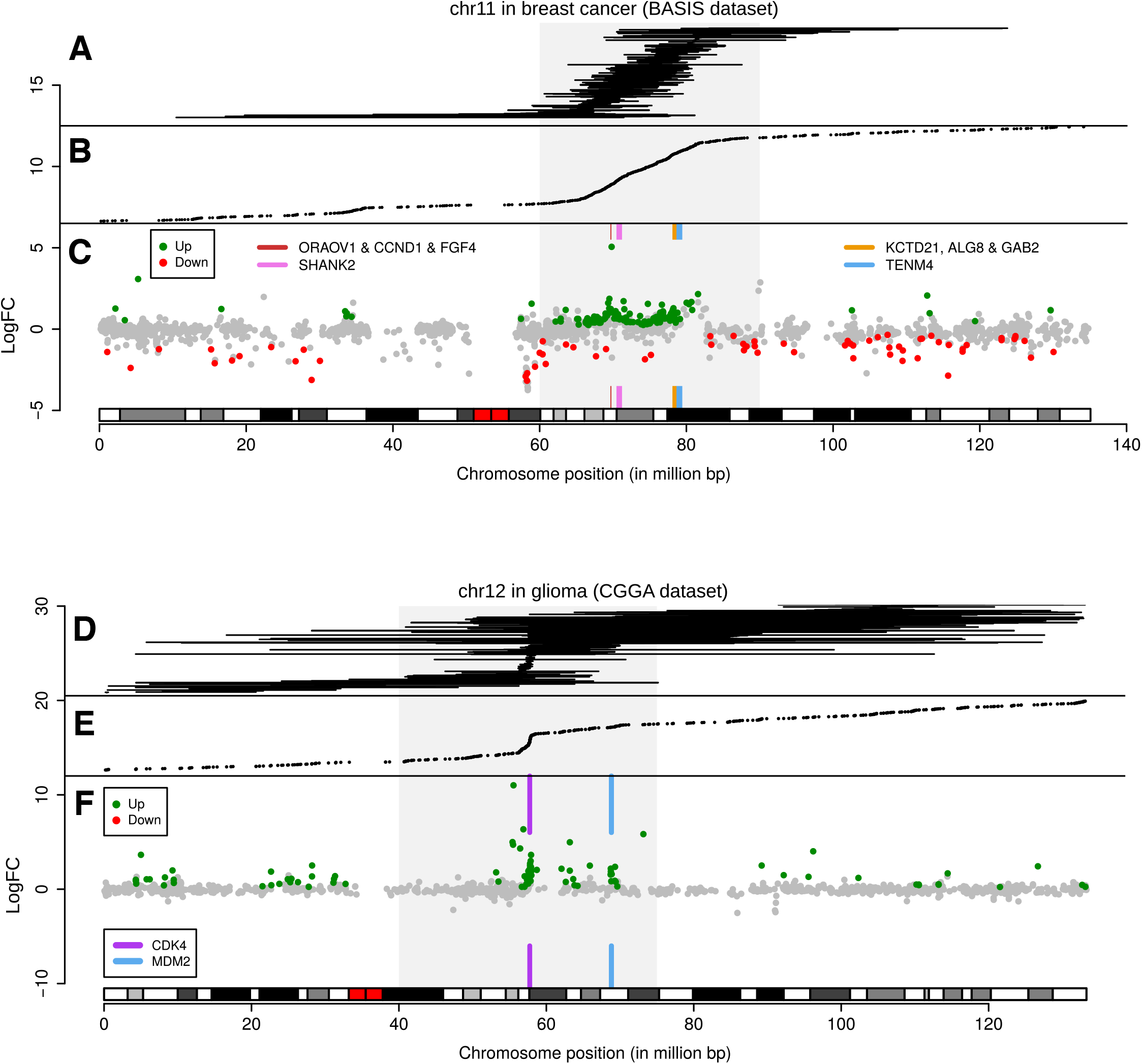
Differential gene expression in junction hotspot regions. (A-C) Overview of chr11 junctions, breakpoint positions and hotspot associated differential gene expression in BrCa, using RNA-seq data only. (A) Intrachromosomal junctions not marked as putative circRNA, indicated by horizontal lines. (B) Breakpoint positions from intronic and exonic, inter- and intrachromosomal junction. (C) Chromosomal differential expression plot for locus chr11:60,000,000-90,000,000 (grey square) with a q-value threshold of 0.001. Genes with the highest number of rearrangements, SHANK2 and TENM4, were illustrated with coloured boxes. Peaks in fold-change were observed surrounding ORAOV1, CCND1 & FGF4 and surrounding TENM4. (D-F) Overview of chr12 junctions, breakpoint positions and hotspot associated differential gene expression in glioma. (D) Intrachromosomal junctions not marked as putative circRNA are indicated with lines. (E) Breakpoint positions from intronic and exonic, inter- and intrachromosomal junction not marked as putative circRNA are included. The breakpoint enriched region chr12:40,000,000-75,000,000 is indicated with a grey square. (F) Chromosomal differential expression plot for locus chr12:40,000,000-75,000,000 with a q-value threshold of 0.01. Peaks in fold change from up-regulated genes are found near CDK4 and MDM2.

Glioma samples having a junction harbouring the chr12 hotspot region (**Figure 7D-F**) were analysed similarly and also showed up-regulation of genes in the hotspot locus, with an increased fold-change of *CDK4, MDM2* and neighbouring genes. Both *CDK4* and *MDM2* are known to be hyper-amplified in glioblastoma ^31^, often by double minute chromosomes ^32^. The Dr. Disco detected junctions showed a sharp increase in close proximity of *CDK4* (**Figure 7D-E**), likely indicating a common start of the amplification event. These breakpoints and up-regulated genes ceased just prior to *LRIG3*. Similarly, glioma samples harbouring rearrangement near the common hyper-amplified *EGFR* showed up-regulation of the surrounding locus (**Figure S20**). These results show that using RNA-seq data only, Dr. Disco can identify genomic breakpoints, which can thereafter be used to reveal associated over-expression of oncogenes which have resulted from high copy gene amplifications.

### Chromothripsis

In VCaP, the q-arm of chr5 has been subjected to chromothripsis as revealed by 468 intrachromosomal WGS-detected breakpoints ^27^. Seventeen intronic and exonic junctions were detected by Dr. Disco in rRNA-minus RNA-seq, identifying evidence for chromothripsis events in VCaP at the (pre-)mRNA level (**Figure S21**). In three BrCa samples, high numbers of WGS-detected genomic breakpoints were identified on the q-arm of chr17 which recurrently involved the genes *BCAS3*, *APPBP2*, *MED13*, *USP32* and *VMP1* (**Figure S22**). RNA-seq analysis revealed intronic and exonic junctions in concordance with the WGS data, which demonstrates the possibility to observe chromothripsis derived junctions in RNA-seq.

### CircRNA detection

Head-to-tail aligned reads (**Figure S23**) are marked as chimeric (discordant) by STAR and are used as input for Dr. Disco. Such reads are not only observed in transcripts from genomic tandem duplications, but also from circular mitochondrial DNA and circular RNAs. Using the PCA-Lines rRNA-minus samples, we found that 88.6% of the junctions with a head-to-tail orientation were located exactly on exon-junctions corresponded to annotated circRNAs from circBase 31 (**Figure S24**). This indicates that Dr. Disco is also capable of identifying circRNAs within rRNA-minus RNA-seq data.

## Discussion

RNA-seq is generally performed on poly(A)^+^ RNA-seq and fusion gene detection algorithms are mostly focused on annotated exons or splice junctions. For a broader understanding of the transcriptome, it has become common practice to sequence ribosome-depleted total RNA (rRNA-minus RNA) ^12^, especially used for partially degraded RNA samples. rRNA-minus RNA-seq is interesting as it yields also non-polyadenylated transcripts and pre-mRNA-derived intronic sequences. As a result, there is more genomic coverage in rRNA-minus RNA-seq alignments and it is closer to whole genome sequencing compared to poly(A)^+^ RNA-seq (**Figure S25**). Because genomic breakpoints are often harboured within introns ^6^ and intergenic regions (**Figure S26**), we interrogated to what extend rRNA-minus RNA-seq can be used to reveal genomic breakpoints as this also captures intronic (pre-mRNA) reads ^10^. Here, we show by utilizing Dr. Disco that rRNA-minus RNA-seq data can indeed reveal exact genomic breakpoints of expressed transcripts, including intergenic translocations. Detection was limited to approximately 10% of all present breakpoints but markedly higher for the driver TMPRSS2-ERG fusion gene (85% detected; 100% presence). Discovering these genomic breakpoints at transcriptional level (RNA) on top of exonic junctions requires an analysis strategy keeping these two levels of information separated. We show that the increased search space combined with graph transformation as implemented in Dr. Disco is a solution to this challenge by providing a unique view on the transcriptome.

CircRNAs are a relatively new group of non-polyadenylated transcripts with more than 90,000 different human circRNAs identified so far ^33,34^. The distinctive signature of proximate exonic head-to-tail junctions sets them apart from other junctions, except for small tandem duplications. A useful addition to the algorithm could be annotation of the junctions using a circRNA database such as circBase^33^. As Dr. Disco is not specifically designed to identify circRNAs and has stringent cut-off levels, the number of circRNAs identified by Dr. Disco is much lower as compared with dedicated detection software such as CIRI^35,36^.

The number of intronic RNA-seq junctions varied largely between the four different cancer types (PCa, BrCa, CRC and glioma). This variation is in line with the omics-reported number of structural variants; low in colorectal cancer ^37^ while high in breast cancer ^38,39^, but is influenced by sequencing depth, read length and library preparation which vary per dataset.

The comparison with WGS data indicated that only a fraction of all genomic rearrangements is transcribed. It is expected that non-transcribed genomic breakpoints more often involve passenger events than transcribed genomic breakpoints. Conversely, oncogene driver events such as TMPRSS2-ERG are characterized by high expression and thus high breakpoint detection rates, as do their mRNA level fusion genes. Known exceptions that can be considered driver events include promoter and enhancer rearrangements such as known for *AR* and *FOXP1* ^40^,but also tumour suppressor gene deletions^41,42^.

Although WGS depth surpasses 40x coverage, Dr. Disco showed that 26% and 48% of all RNA-seq intronic breaks in PCa and BrCa, respectively, were not identified by WGS. Multiple reasons may explain this discrepancy; high RNA-seq coverage of highly expressed genes (up-to 1000x), clonality as this difference was in particular high for a small subset of samples, local low coverage in DNA-seq, intergenic exonic junctions not spanning canonical splice junctions, and selection criteria in software such as conservative cut-offs for genomic breakpoint detection and read mapping rulings but they may also contain false positives. For Dr. Disco, both read-length and coverage are directly linked to the number of detected genomic breakpoints and fusion splice junctions. In the PCa-LINES FFPE dataset, we found that samples with low insert sizes or short read lengths resulted in insufficient split-reads whilst resulting in many false positive read-pairs in the full transcriptome analysis, but could still be used effectively in identifying the targeted, highly expressed, *TMPRSS2*-*ERG* fusion events.

From our Dr. Disco analyses, we were able to resolve the genomic breakpoints and splice variants for various known and novel fusion events. The PCa-specific *TMPRSS2*-*ERG* fusions and breakpoints were investigated in detail and revealed additional cryptic and intergenic exons including TMPRSS2 exon-0 ^28^ and breakpoints located before *ERG*. For some of these fusion events (e.g. VCaP cell line), the genomic rearrangement is complex and consists of insertions, deletions and inversions. The use of stranded RNA-seq provides an advantage in deciphering complex genomic rearrangements. In VCaP, an inversion results in partial anti-sense transcription from which the chronological order of events can be deduced. The manual unravelling of the complex TMRPSS2-ERG variant in VCaP shows the importance of automatic resolution of complex genomic rearrangements or poly-fusions. The current implementation of Dr. Disco does not offer top-level integration for poly-fusions but there are methods available with that aim ^7,43^. In addition, the effect of enhancer/promoter rearrangements and head-to-head gene fusions on the local transcriptome landscape can be resolved by stranded RNA-seq. Besides their unique genomic breakpoints, complex genomic rearrangements harbouring inversions are also characterized by regions with opposite strand transcription. Since the current Dr. Disco algorithm uses discordant reads exclusively, extending it with the detection of regions enriched with concordant opposite stranded reads may strengthen detection of genomic rearrangements having insufficient breakpoint coverage. RNA-seq data can reveal genomic breakpoints, (cryptic and/or intergenic) splicing and gene expression information, which together can reveal consequences and their selective advantage for cancer development and progression and be a useful supplement to DNA-seq.

In both BrCa and glioma, RNA-seq data revealed hotspot regions of junctions with the subsequent up-regulation of known amplified oncogenes within these regions. This integrated RNA-seq analysis utilizing recurrent junctions coupled with gene expression analysis of neighbouring genes directly uncovered known oncogenes. This shows which changes at RNA level are most prominent, and thus which genes are most strongly influenced by these genomic aberrations. Then the direction of transcription provides additional context, by showing that there are no consistent acceptor/donor genes. Indeed, as DNA detection of translocations is the golden standard, a combined RNA DNA analysis would yield more comprehensive results. Furthermore, the expression analysis indicated that certain detected fusion transcripts such as TEM4 and SHANK2 fusions are likely not driving cancer in these cases. In BrCa, the RNA detected junctions originating from driver gene amplifications were often located within the sizeable genes *SHANK2* and *TENM4*. It is likely that selection of breakpoints near *SHANK2* is influenced by being adjacent to *CTTN*, a gene containing an enhancer often co-amplified with *CCND1* ^41^. The hotspots found in *TMPRSS2*-*ERG*, *CCDN1*, *CKD4*/*MDM2* were based on frequent events, but also rare and single combinations of transcribed rearrangements and aberrant gene expression can be extracted from Dr. Disco employed on RNA-seq data.

In the VCaP cell line and three BrCa samples, chromothripsis derived junctions were observed at RNA level. Similar to the observation of regular genomic rearrangements, the majority of the chromothripsis rearranges were not detected on RNA level. Solely based on RNA-seq data, it will be difficult to prove presence of chromothripsis as not all parameters that define this specific process can readily be extracted (e.g. copy-number variations, short insertions, loss of heterozygosity) ^44,45^. However, potential indicators for chromothriptic events within cancer cells can be extracted using Dr. Disco.

Approximately 35% of the junctions in rRNA-minus datasets were full or partial intergenic events, of which exonic junctions often included cryptic exons. Also, for well-known in-frame gene-gene fusions such as *TMPRSS2*-*ERG*, many novel cryptic exons were identified that, although often rare, can result in sections of nonsense protein. Cryptic exons may encode completely novel neo-antigens that are more divergent than point mutation-based neo-antigens and could therefore likely be more immunogenic ^46^. Deciphering the consequence of rearrangements, annotation of novel cryptic exons and their coding potential for nonsense protein sequences is therefore relevant for therapeutic interventions using tumour-specific antigens^47^.

Facilitated by Dr. Disco, we set out to extract both intronic and exonic junctions from comprehensive rRNA-minus RNA-seq datasets and identified novel DNA breakpoints, circRNAs, gene fusions, cryptic exons, chromothripsis events and were able to link expressed rearrangements to transcriptional outcome. Performing analysis as presented can be an informative supplement to WGS analysis because of stranding, expression levels and analysis of gene structures. However, in case of lacking WGS data such analysis can provide additional information as compared to poly(A)^+^ RNA-seq, but will require deeper coverage to achieve similar exon depth. Thus, rRNA-minus RNA-seq provides unique and more complete information on non-polyadenylated and aberrant transcripts and, if the pre-mRNA is sequenced, the genomic breakpoints that underlie transcriptional changes.

## Methods

### Sequencing and datasets

Datasets analysed in this study are the BrCa dataset BASIS (n=207) ^23,24^, NGS-ProToCol (normal adjacent prostate; n=41, prostate cancer; n=51; normal adjacent colon; n=18, colorectal adenoma; n=30 and colorectal carcinoma; n=30) ^34,48^ and the Chinese glioma atlas CGGA (various glioma types; n=274) ^49^ of which the data accession identifiers are given in **Table 1**.

For the NGS-ProToCol cohort, RNA was extracted using RNA-Bee (Campro Scientific, Berlin, Germany) and the library prepared for RNA-seq used the NEBNext Ultra Directional RNA Library Prep Kit for Illumina with rRNA reduction. The sample preparation was performed according to the protocol ‘*NEBNext Ultra Directional RNA Library Prep Kit for Illumina*’ (NEB, Cat. #E7420S/L and E6310S/L/X). Briefly, rRNA was reduced using RNase H-based method. Then, fragmentation of the rRNA reduced RNA and a cDNA synthesis was performed. This was used for ligation with the sequencing adapters and PCR amplification of the resulting product. The quality and yield after sample preparation were measured with the Fragment Analyzer (Advanced Analytical). Clustering and DNA sequencing using the Illumina cBot and HiSeq 2500 was performed according to manufacturer’s protocols. A concentration of 16.0 pM of DNA was used as input. HiSeq control software HCS (v2.2.58) was used. Image analysis, base calling, and quality check was performed with the Illumina data analysis pipeline RTA (v1.18.64) and Bcl2fastq (v2.17). The 126 bp stranded Illumina HiSeq 2500 paired-end reads have a peak in fragment size of 300-600 bp and the samples have an average depth of 70 million paired-end reads.

The PCa-LINES dataset consists of PCa cell lines PC346C and VCaP and additional PCa patient samples G-089, G-110, G-295, G-316 and G-346. Each of these samples were WGS DNA sequenced and processed using the complete genomics platform ^27,50^. The matching poly(A)^+^ RNA-seq samples were taken from the TraIT-Cell Line Use Case ^51,52^. The matching rRNA-minus samples of G-089, G-295, G-316, G-346, VCaP and PC346C were processed similarly as the rRNA-minus samples from the NGS-ProToCol dataset. rRNA-minus RNA-seq sample G-110 was sequenced in the NGS-ProToCol study as sample7046-004-052.

In the BASIS RNA-seq dataset, total RNA was extracted and cleaned from abundant RNAs such as rRNA and tRNA using duplex-specific nuclease treatment prior to random primed cDNA synthesis ^53^. The BASIS DNA-seq data preparation and analysis is described elsewhere ^23^ and coordinates were converted to hg38 using *pyliftover* where needed.

The detection of genomic breakpoints from additional TMPRSS2-ERG fusions determined by targeted DNA-seq was described elsewhere ^26^ and genomic coordinates were obtained from this study accordingly. DNA breakpoints of TMPRSS2-ERG and chromothripsis on chr5 in VCaP were described elsewhere ^26,27^. The predicted CMS classes for the NGS-ProToCol colon samples were described elsewhere ^48^.

### Computational data analysis

RNA-seq data was aligned with STAR ^54^ version 2.4.2 with fusion settings using hg38 as reference genome. A more detailed description of the used methodology is given in **Supplementary methods**. Dr. Disco version 0.17.8 (git commit 2a9ff32950b71029b124ff4d16544b2953c57dbe) was used for all analysis. Dr. Disco is available under a Free Open-Source Software license at the website: https://github.com/yhoogstrate/dr-disco. For this study, we designed a free software package to generate Lorenz and coverage plots: https://github.com/yhoogstrate/bam-lorenz-coverage. Processed bam files used to estimate general genome coverage statistics were obtained from EGAS00000000052 ^55^. Pathway enrichment was performed with g:Profiler (https://biit.cs.ut.ee/gprofiler/gost) ^56^ using gene identifiers as a non-ordered query. For differential expression analysis, the annotation of the results of Dr. Disco and further integration with gene sets for determining intergenic and protein coding status, Ensembl gene annotation 89 was used.

Plots were made with R 3.6.2 (base R, ggplot2, plotrix and circlize) and illustrations with Inkscape. Differential gene expression analysis was performed using the edgeR 3.2.8 library ^57^. Associations between the frequency of breakpoints per sample and clinical parameters were tested using the Mann Whitney U test in R. For the Venn diagrams describing overlap across intronic, exonic and WGS junctions, both sides of the junctions must be within 40 genomic nucleotides in proximity to be considered a match.

Chromosomal differential expression plots were made using base R. For a given locus and q-value threshold, a cohort is separated in a mutant and wildtype group by having one or more intronic or exonic junctions within the given locus. Differential expression analysis is performed across these groups using edgeR. Every gene located on the chromosome on which the locus is located, is plotted with its genomic centre as defined by Ensembl 89 on the x-axis and with edgeR’s log fold change on the y-axis. A gene that is up-regulated in the mutant group has a positive log fold change and a gene that is down regulated a negative log fold change. When the gene is not significantly differentially expressed across the wildtype and mutant group (q-value below predetermined threshold) the gene will be coloured grey. If the difference is significant, it will be coloured green (up) or red (down).

## Supporting information

Table S1 - RNA only SVs 7x PCa

Table S2 - Recurrent Fusion Genes

Table S3 - All detected SVs

Table S4 - gProfiler

Table S5 - TMPRSS2-ERG and related

Table S6 - SHANK2 and TENM4 related

Dr. Disco technical specification

## Data Access

Dr. Disco is available at the following url: https://github.com/yhoogstrate/dr-disco.

Raw sequencing is accessible at the following public repositories: EGAS00001002816, EGAS00001002854, EGAS00001001178, EGAD00001006366, GSE48865, EGAS00001001178, EGAS00001001476 (**Table 1**).

The concatenated results on all samples (**Table S3**) using the Dr. Disco v.0.17.8 pipeline is available at: https://doi.org/10.5281/zenodo.4159414.

## Disclosure Declaration

For the CTMM NGS-ProToCol study (NGS-ProToCol, Next Generation Sequencing from Prostate to Colorectal Cancer - Center for Translational Molecular Medicine (2014-2015); https://www.lygature.org/ctmm-portfolio), 51 prostate cancers from the Erasmus MC were snap-frozen and stored in liquid nitrogen as previously described ^58^. Use of the samples for research purposes was approved by the Erasmus MC Medical Ethics Committee according to the Medical Research Involving Human Subjects Act (MEC-2004-261; MEC-2010-176).

Other data was obtained from publicly available studies.

The authors declare no competing interests.

## Funding

This study was performed within the framework of the CTMM (Center for Translational Molecular Medicine) research program; NGS-ProToCol [grant 03O-402]; PCMM [grant 03O-203-1]; Translational Research IT (TraIT); the Complete Genomics Inc. grant [EMC GL 083111]; the FP7 Marie Curie Initial Training Network PRO-NEST [grant number 238278] and Support for the Cancer Computational Biology Center was provided by the Daniel den Hoed Foundation. Funding for open access charge: NGS-ProToCol [grant 03O-402].

## Legends supplementary files

**Additional file 1** – Supplementary Figures S1-S26

**Additional file 2** – Table S1

Dr. Disco detected intronic junctions expected to be genomic breakpoints but not matching with WGS detected breakpoints in the 7 PCa samples with matching poly(A)^+^ and rRNA-minus RNA-seq data (PCa-LINES dataset). Results are ordered by presence in either rRNA-minus, poly(A)^+^ or both datasets.

**Additional file 3** – Table S2

Recurrent fusion genes as found by Dr. Disco in the NGS-ProToCol prostate cancer and colon datasets and the BASIS breast cancer dataset. Glioma samples were excluded because they were sequenced unstranded. Fusion genes present in at least 2 samples of the same tumour type are considered recurrent; only entries that passed filtering and were marked as ‘linear’ to avoid circRNA entries were included; both intronic and exonic entries were included but were de-duplicated per sample; only 1 unique occurrence of a fusion gene per sample; no self-fusions (TMPRSS2-TMPRSS2); no intergenic fusions; no fusions involving chrM or alternate loci. If there are multiple genes spanning the breakpoint, the Cartesian product of the gene names is used; when A,B -> C is found, this is expanded to: 1x A->C and 1x B->C.

**Additional file 4** – Table S3

Large concatenated results table on all samples of the Dr. Disco study. Available online because of the large file size: https://doi.org/10.5281/zenodo.4159414.

**Additional file 5** – Table S4

G:Profiler pathway enrichment analysis on genes that are recurrently hit. (**A**) ER-negative BrCa samples from the BASIS cohort; (**B**) ER-positive BrCa samples from the BASIS cohort; (**C**) glioma samples from the CGGA and (**D**) PCa samples from the NGS-ProToCol cohort. Colon samples were not included because of the relatively small number of recurrently hit genes. For the BrCa dataset only genes that were hit in 3 or more distinct samples were used in the analysis. For the glioma and PCa samples, only genes that were hit in 2 distinct samples were used in the analysis. Entries suspected to be circRNAs were excluded.

**Additional file 6** – Table S5

The concatenated Dr. Disco detected junctions related to **TMPRSS2**-**ERG** in the NGS-ProToCol PCa samples.

**Additional file 7** – Table S6

Dr. Disco output of detected junctions related to **SHANK2** and/or **TENM4** as found in the BASIS BrCa dataset.

**Additional file 8** – Supplementary methods

**Additional file 9** – Dr. Disco technical specification

## Notes

https://doi.org/10.5281/zenodo.4159414

### Competing Interest Statement

The authors have declared no competing interest.

