## Supplementary material for "Detection of fusion transcripts and their genomic breakpoints from RNA sequencing data": Supplemenatry Figures

### Slide 1
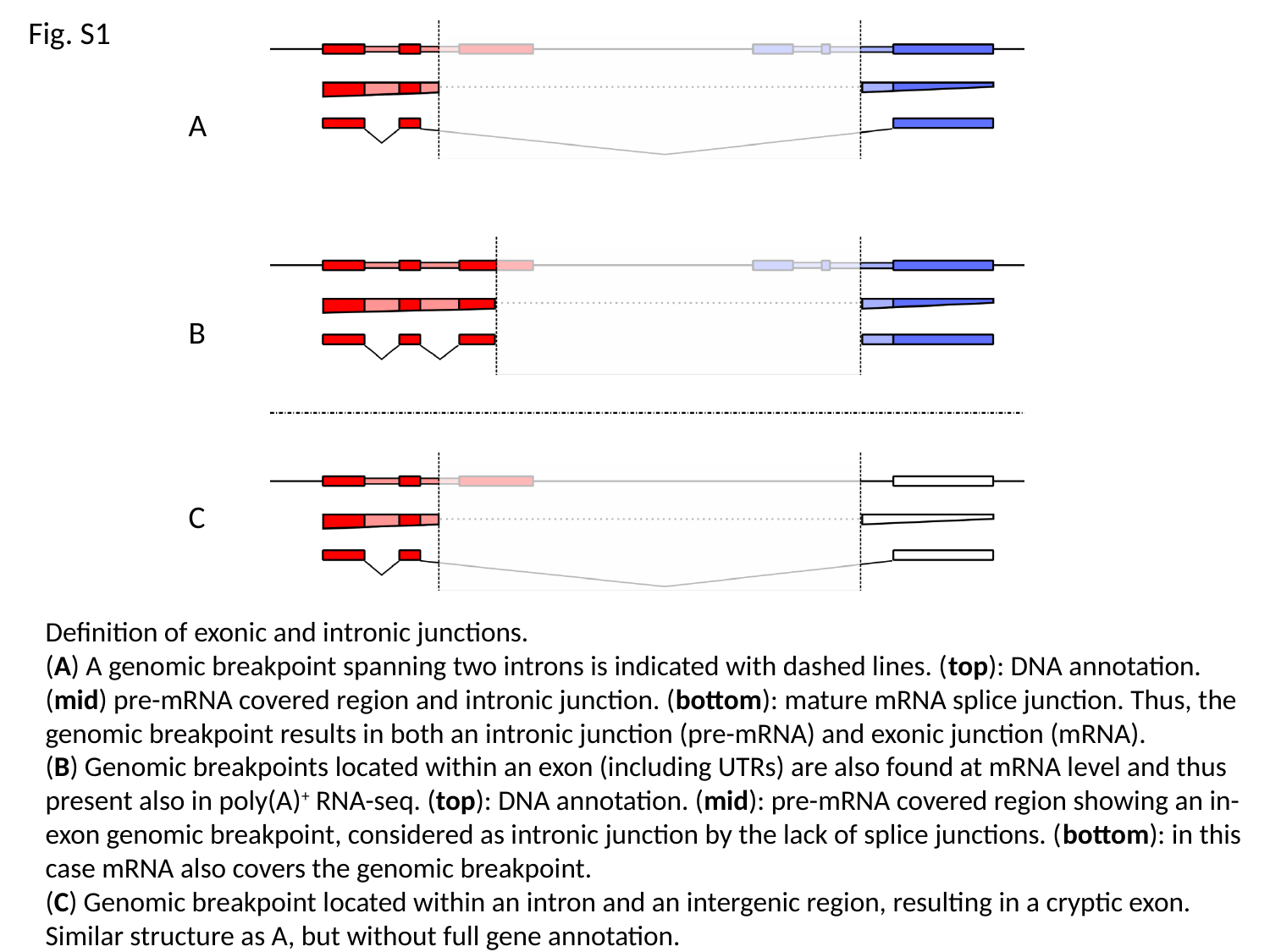

Fig. S1
A
B
C
Definition of exonic and intronic junctions.
(A) A genomic breakpoint spanning two introns is indicated with dashed lines. (top): DNA annotation. (mid) pre-mRNA covered region and intronic junction. (bottom): mature mRNA splice junction. Thus, the genomic breakpoint results in both an intronic junction (pre-mRNA) and exonic junction (mRNA).
(B) Genomic breakpoints located within an exon (including UTRs) are also found at mRNA level and thus present also in poly(A)^+^ RNA-seq. (top): DNA annotation. (mid): pre-mRNA covered region showing an in-exon genomic breakpoint, considered as intronic junction by the lack of splice junctions. (bottom): in this case mRNA also covers the genomic breakpoint.
(C) Genomic breakpoint located within an intron and an intergenic region, resulting in a cryptic exon. Similar structure as A, but without full gene annotation.

### Slide 2
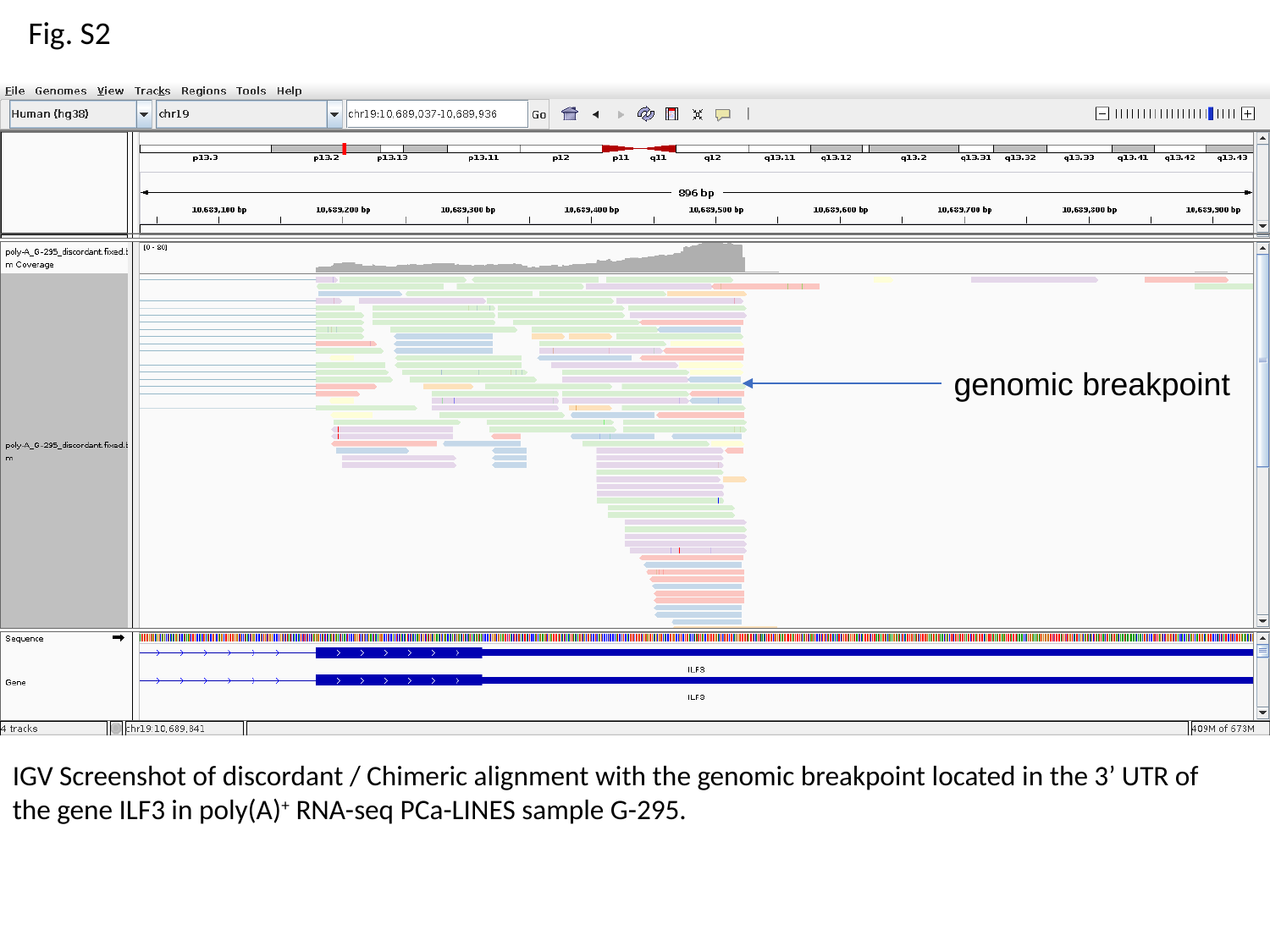

Fig. S2
genomic breakpoint
IGV Screenshot of discordant / Chimeric alignment with the genomic breakpoint located in the 3’ UTR of the gene ILF3 in poly(A)^+^ RNA-seq PCa-LINES sample G-295.

### Slide 3
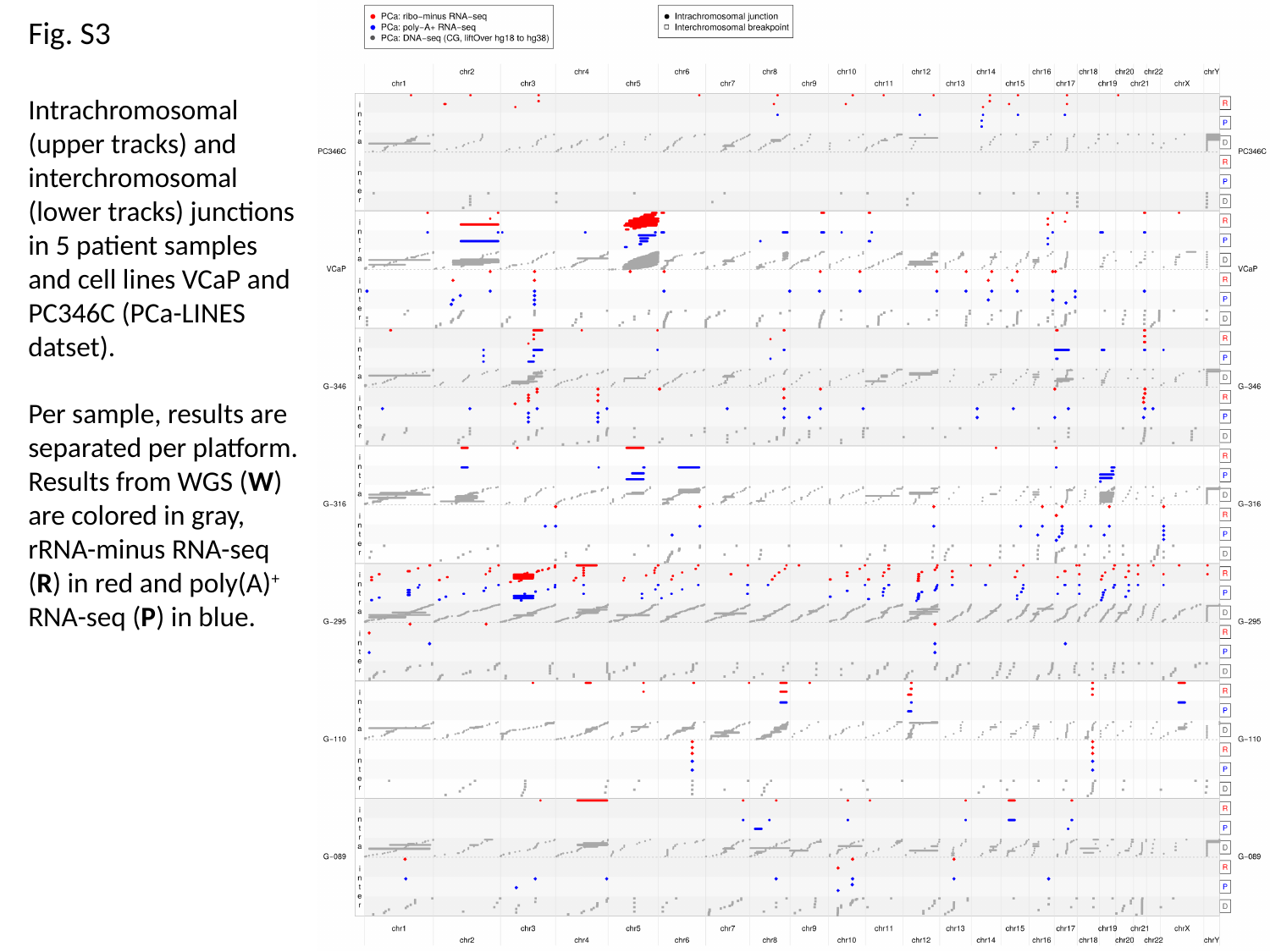

Fig. S3
Intrachromosomal (upper tracks) and interchromosomal (lower tracks) junctions in 5 patient samples and cell lines VCaP and PC346C (PCa-LINES datset).
Per sample, results are separated per platform. Results from WGS (W) are colored in gray, rRNA-minus RNA-seq (R) in red and poly(A)^+^ RNA-seq (P) in blue.

### Slide 4
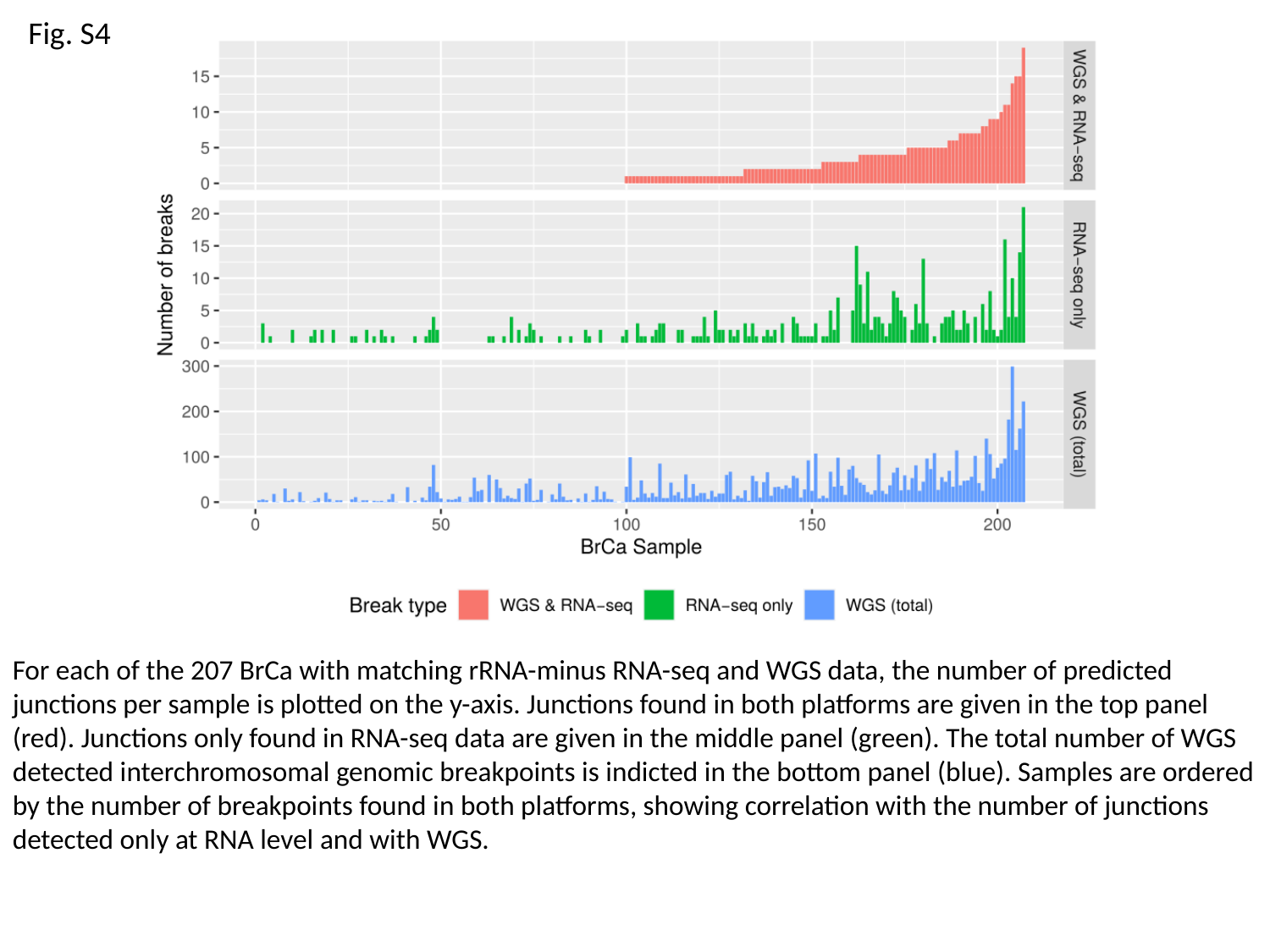

Fig. S4
For each of the 207 BrCa with matching rRNA-minus RNA-seq and WGS data, the number of predicted junctions per sample is plotted on the y-axis. Junctions found in both platforms are given in the top panel (red). Junctions only found in RNA-seq data are given in the middle panel (green). The total number of WGS detected interchromosomal genomic breakpoints is indicted in the bottom panel (blue). Samples are ordered by the number of breakpoints found in both platforms, showing correlation with the number of junctions detected only at RNA level and with WGS.

### Slide 5
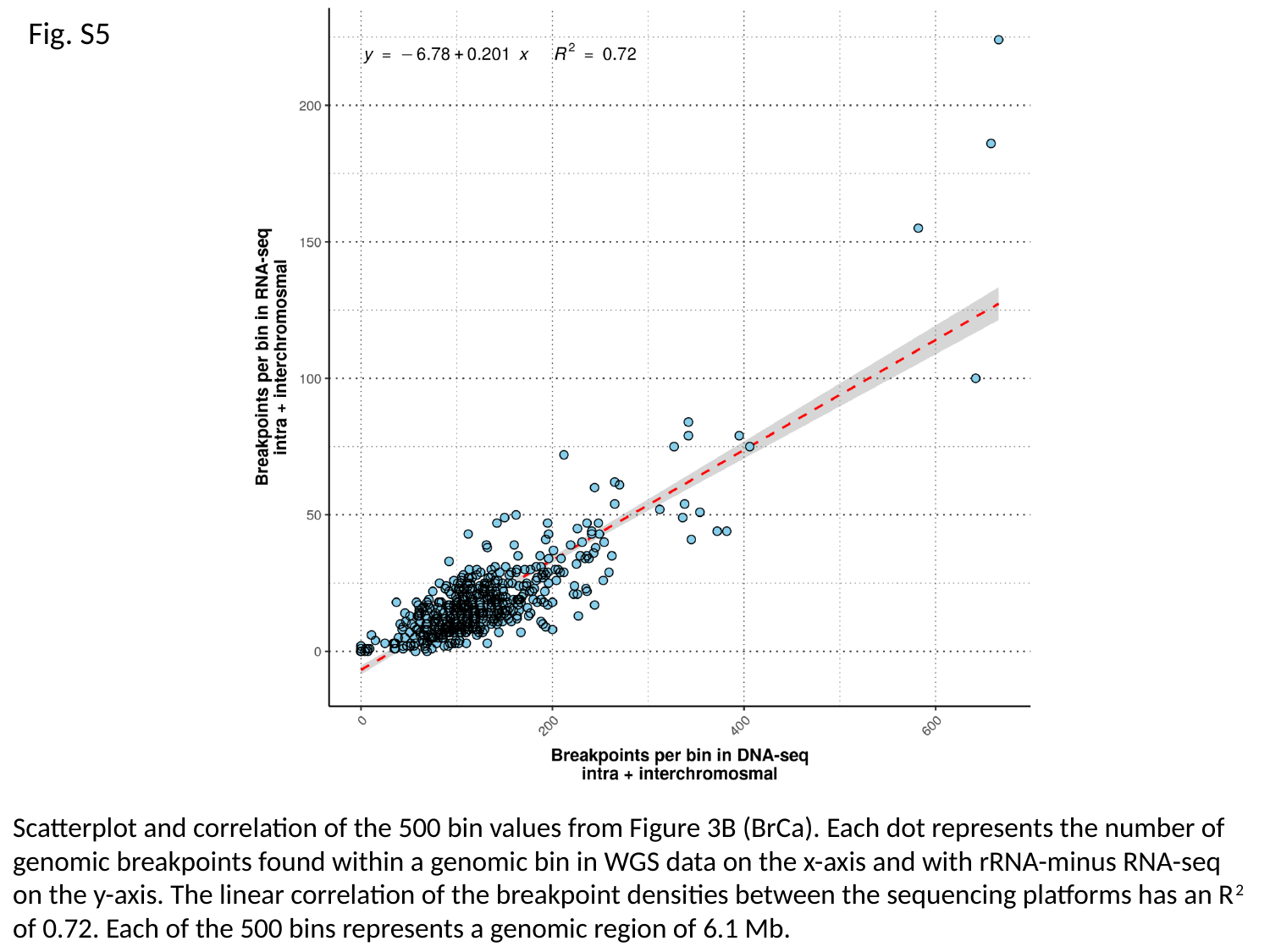

Fig. S5
Scatterplot and correlation of the 500 bin values from Figure 3B (BrCa). Each dot represents the number of genomic breakpoints found within a genomic bin in WGS data on the x-axis and with rRNA-minus RNA-seq on the y-axis. The linear correlation of the breakpoint densities between the sequencing platforms has an R^2^ of 0.72. Each of the 500 bins represents a genomic region of 6.1 Mb.

### Slide 6
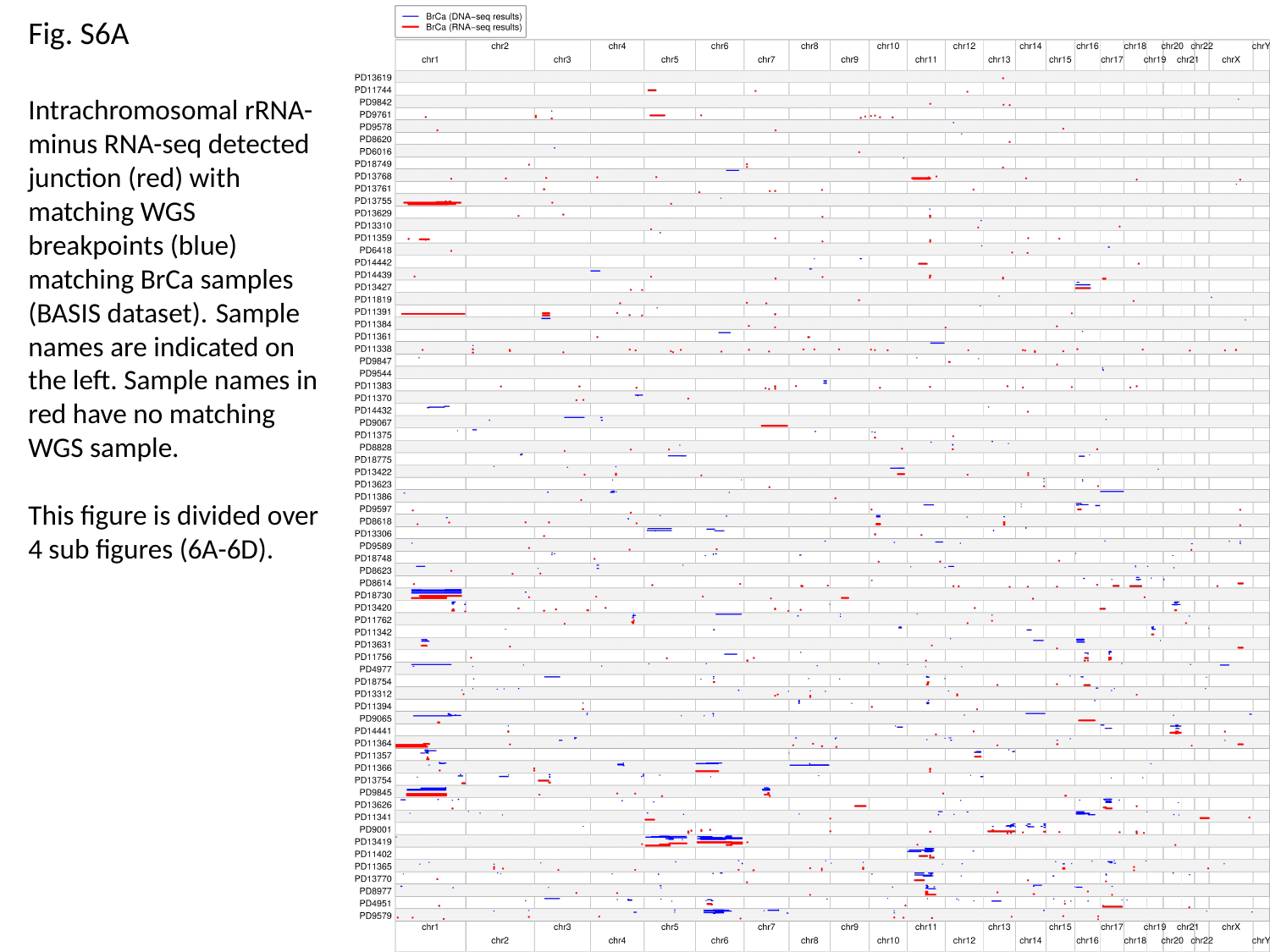

Fig. S6A
Intrachromosomal rRNA-minus RNA-seq detected junction (red) with matching WGS breakpoints (blue) matching BrCa samples (BASIS dataset). Sample names are indicated on the left. Sample names in red have no matching WGS sample.
This figure is divided over 4 sub figures (6A-6D).

### Slide 7
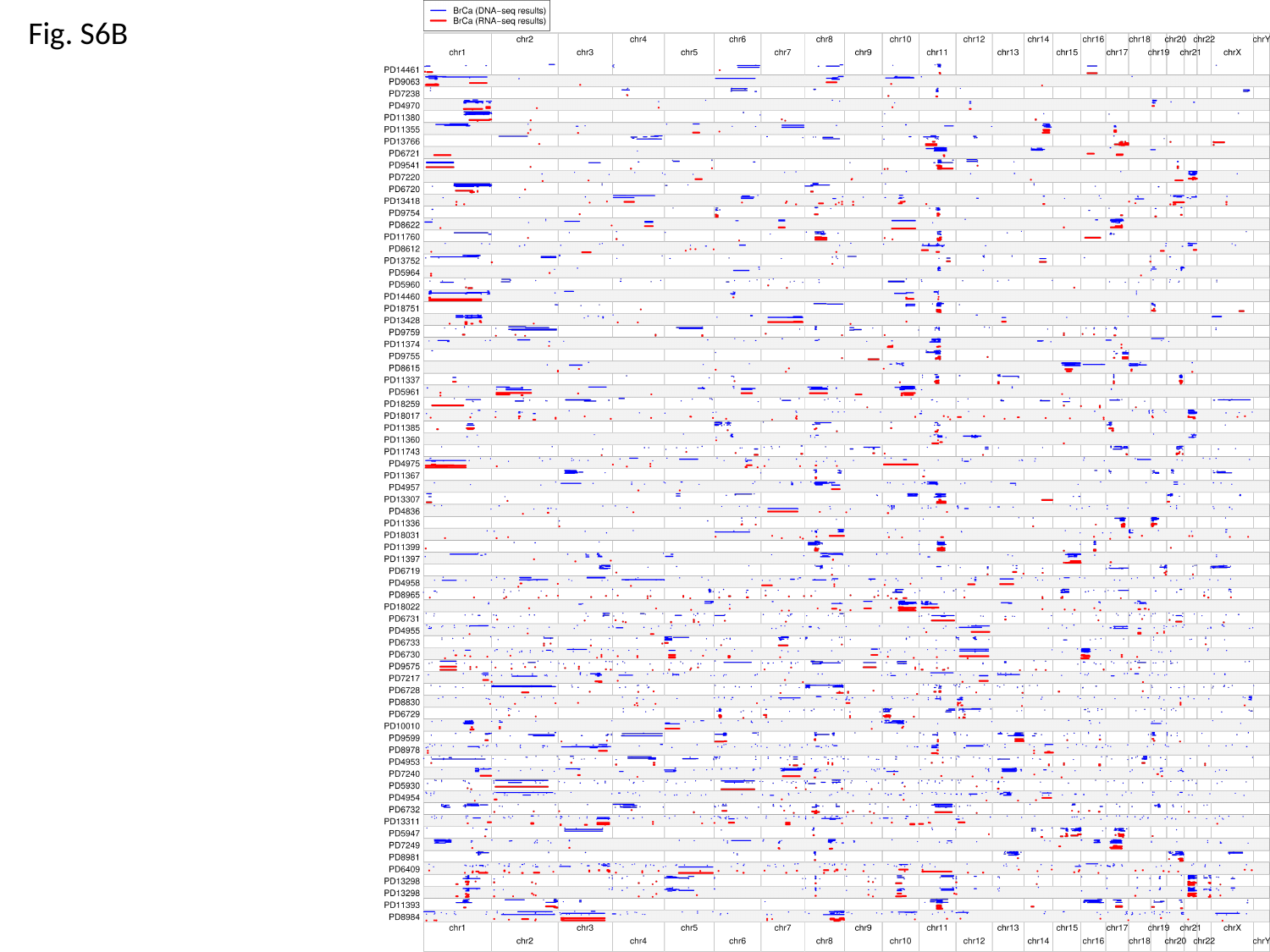

Fig. S6B

### Slide 8
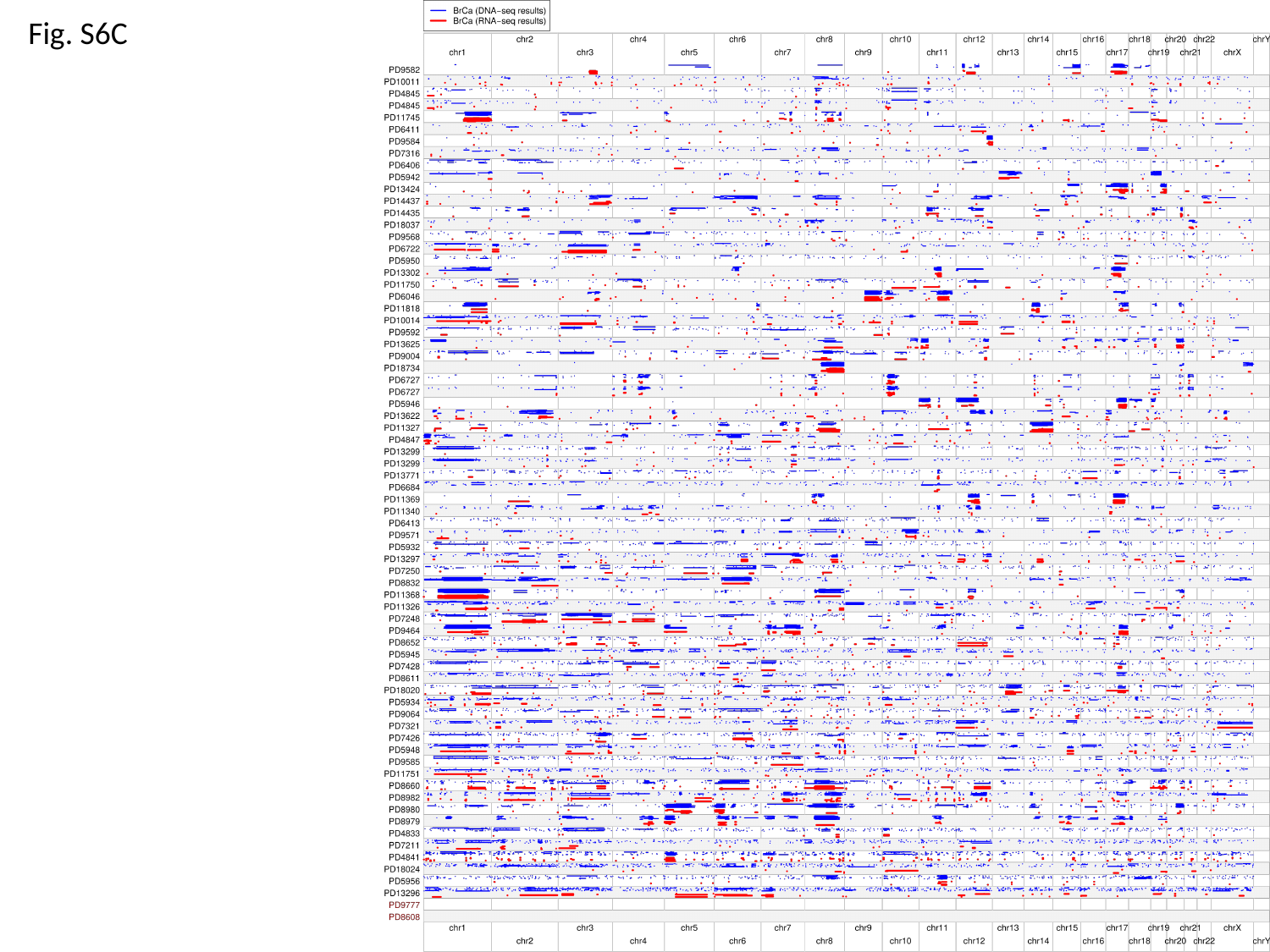

Fig. S6C

### Slide 9
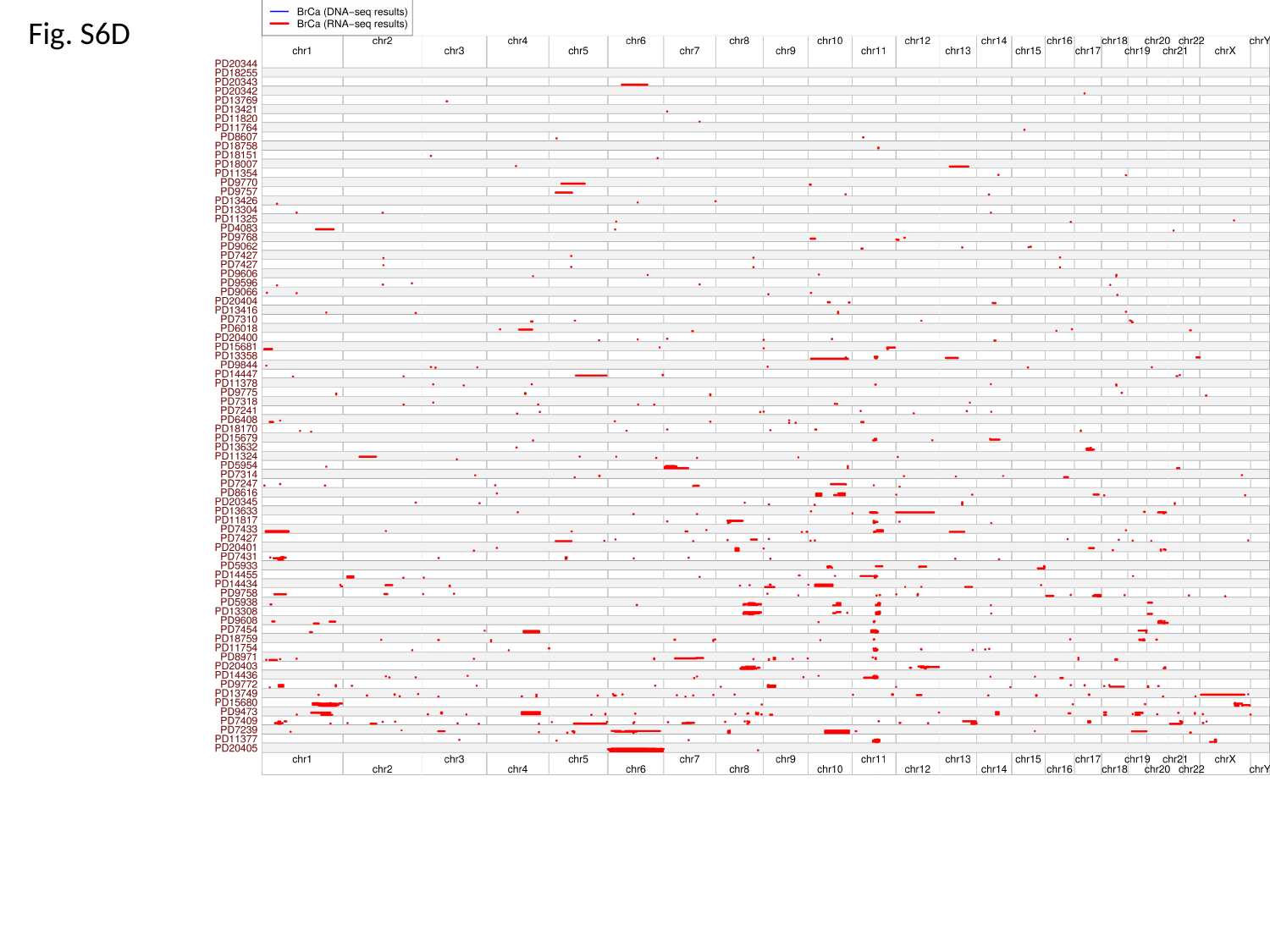

Fig. S6D

### Slide 10
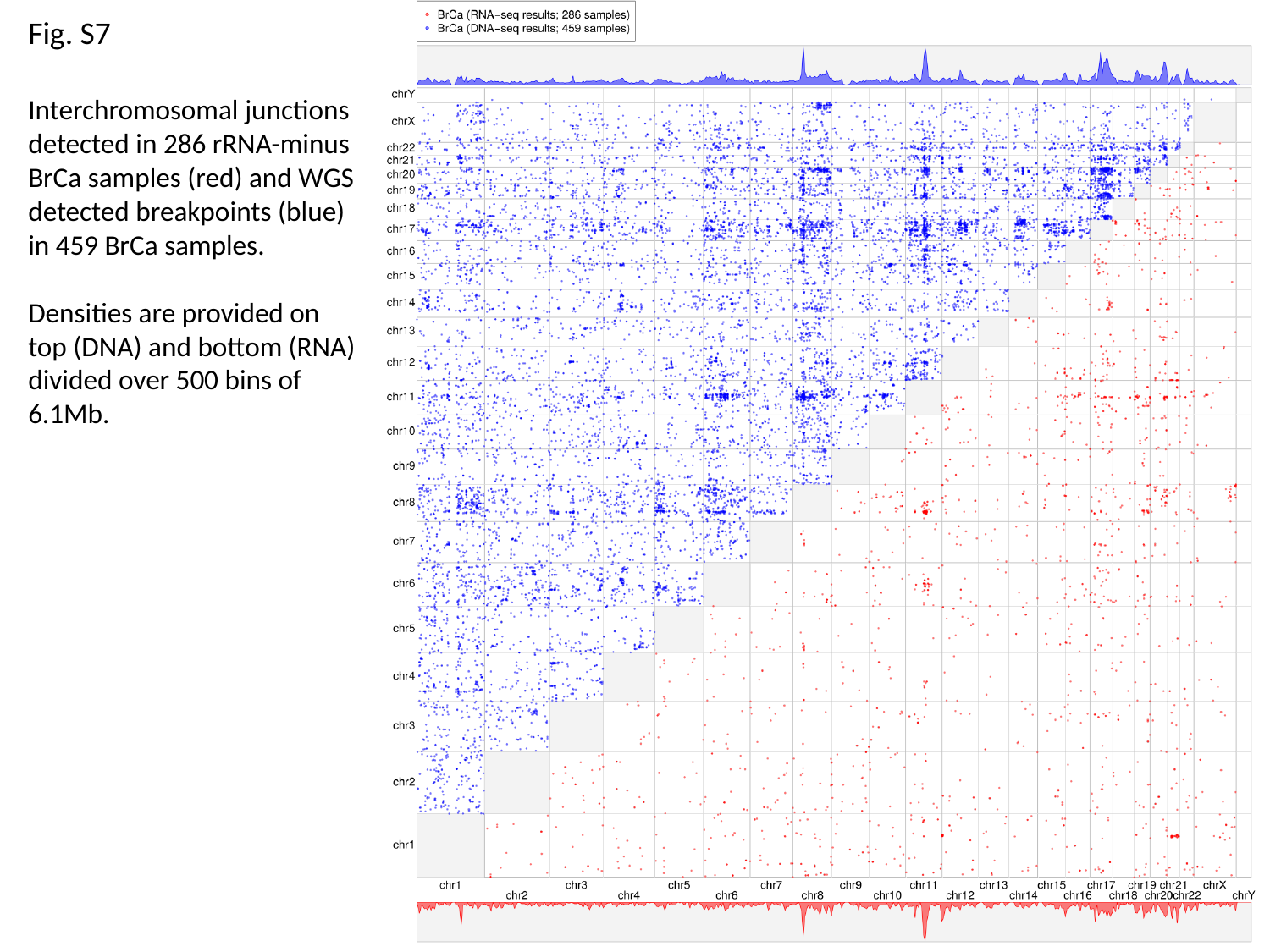

Fig. S7
Interchromosomal junctions detected in 286 rRNA-minus BrCa samples (red) and WGS detected breakpoints (blue) in 459 BrCa samples.
Densities are provided on top (DNA) and bottom (RNA) divided over 500 bins of 6.1Mb.

### Slide 11
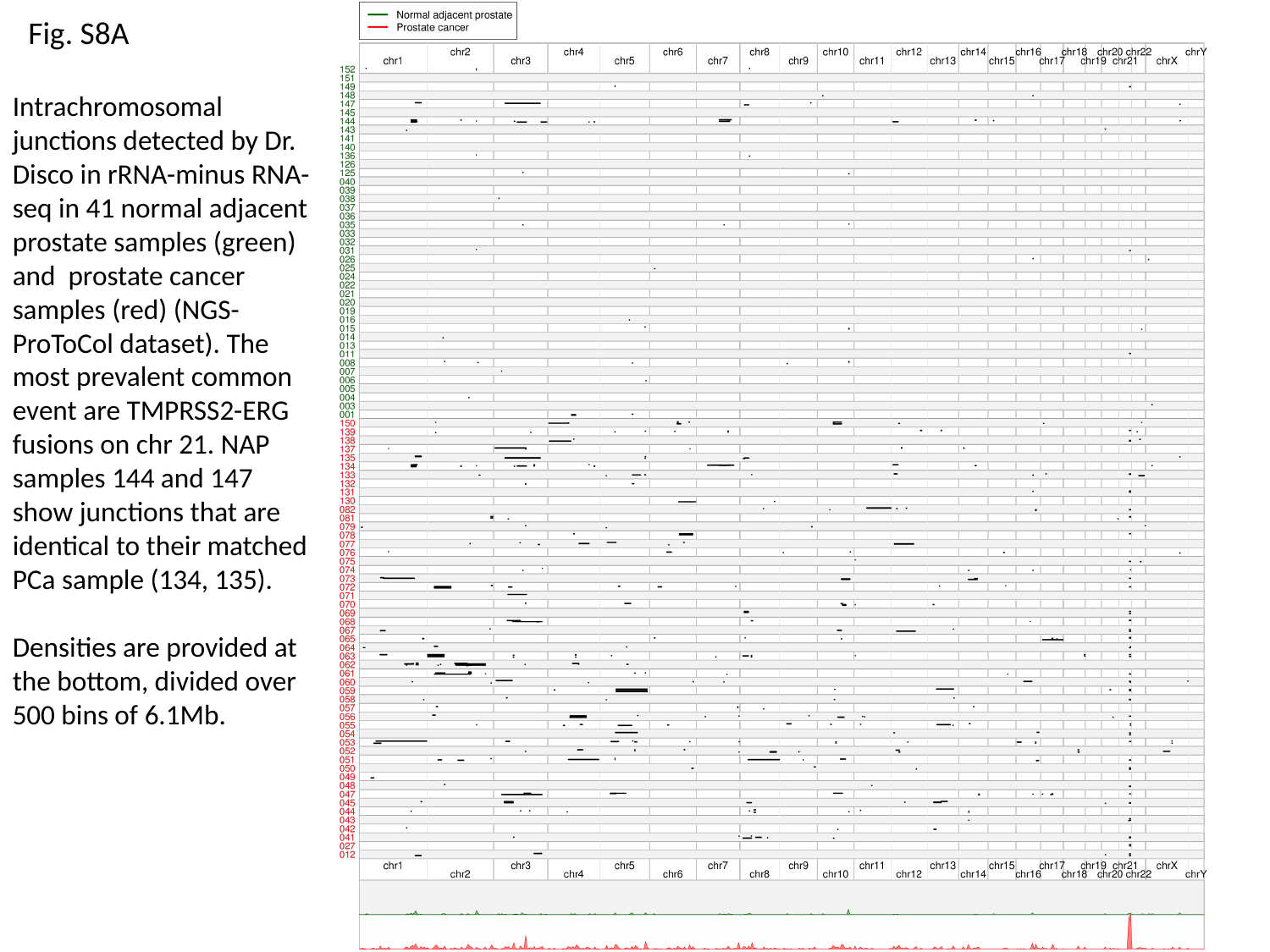

Fig. S8A
Intrachromosomal junctions detected by Dr. Disco in rRNA-minus RNA-seq in 41 normal adjacent prostate samples (green) and prostate cancer samples (red) (NGS-ProToCol dataset). The most prevalent common event are TMPRSS2-ERG fusions on chr 21. NAP samples 144 and 147 show junctions that are identical to their matched PCa sample (134, 135).
Densities are provided at the bottom, divided over 500 bins of 6.1Mb.

### Slide 12
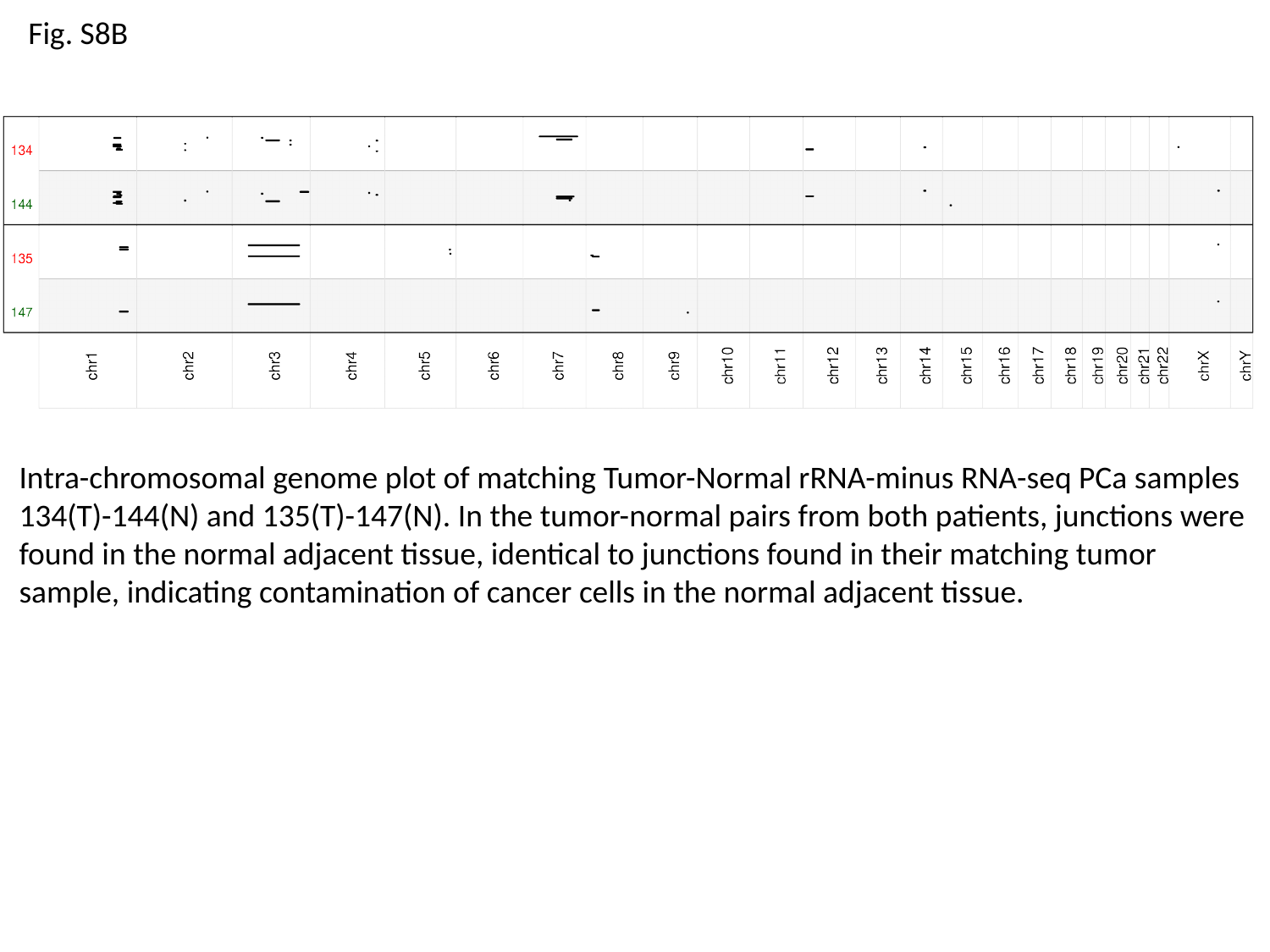

Fig. S8B
Intra-chromosomal genome plot of matching Tumor-Normal rRNA-minus RNA-seq PCa samples 134(T)-144(N) and 135(T)-147(N). In the tumor-normal pairs from both patients, junctions were found in the normal adjacent tissue, identical to junctions found in their matching tumor sample, indicating contamination of cancer cells in the normal adjacent tissue.

### Slide 13
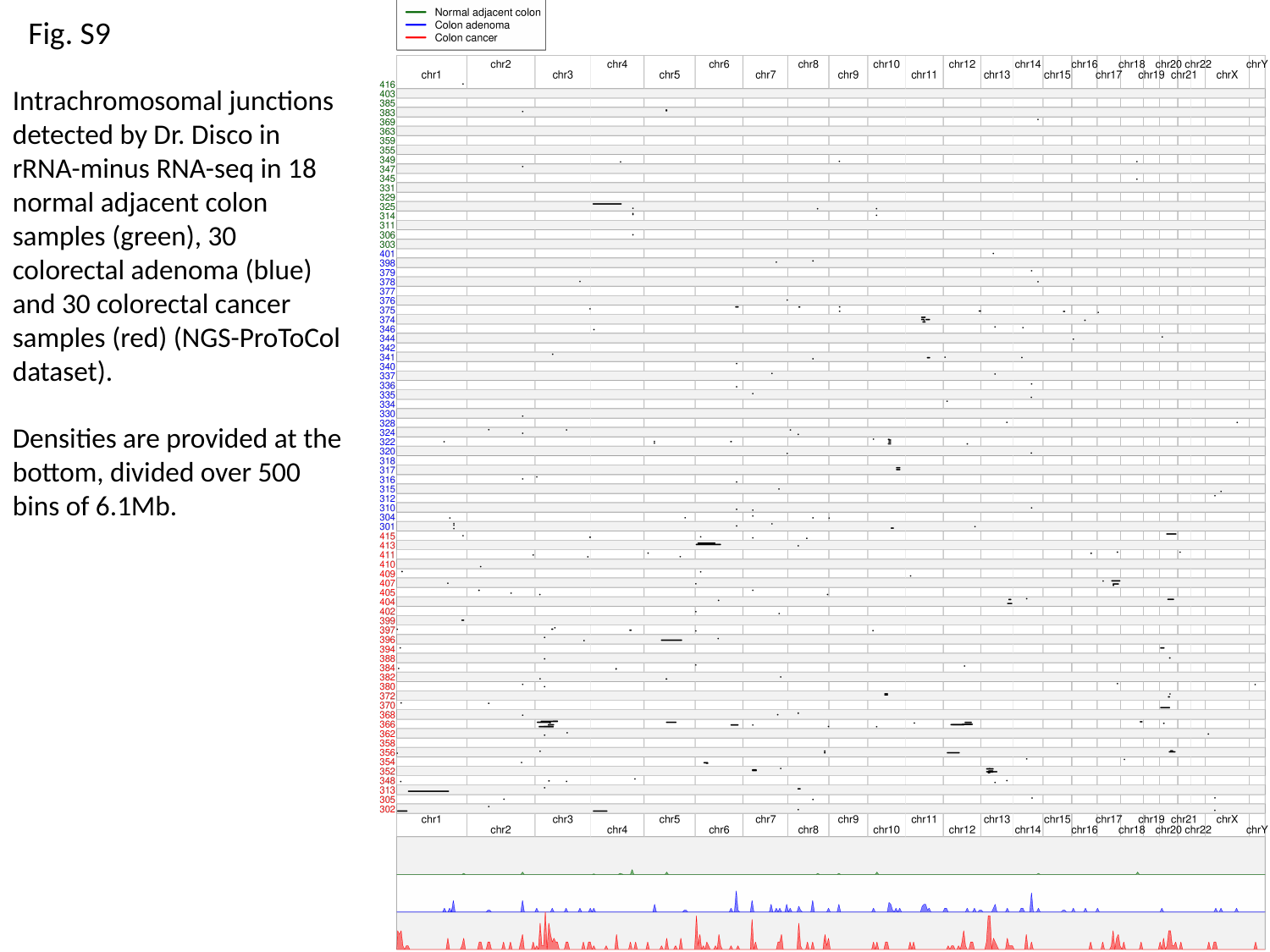

Fig. S9
Intrachromosomal junctions detected by Dr. Disco in rRNA-minus RNA-seq in 18 normal adjacent colon samples (green), 30 colorectal adenoma (blue) and 30 colorectal cancer samples (red) (NGS-ProToCol dataset).
Densities are provided at the bottom, divided over 500 bins of 6.1Mb.

### Slide 14
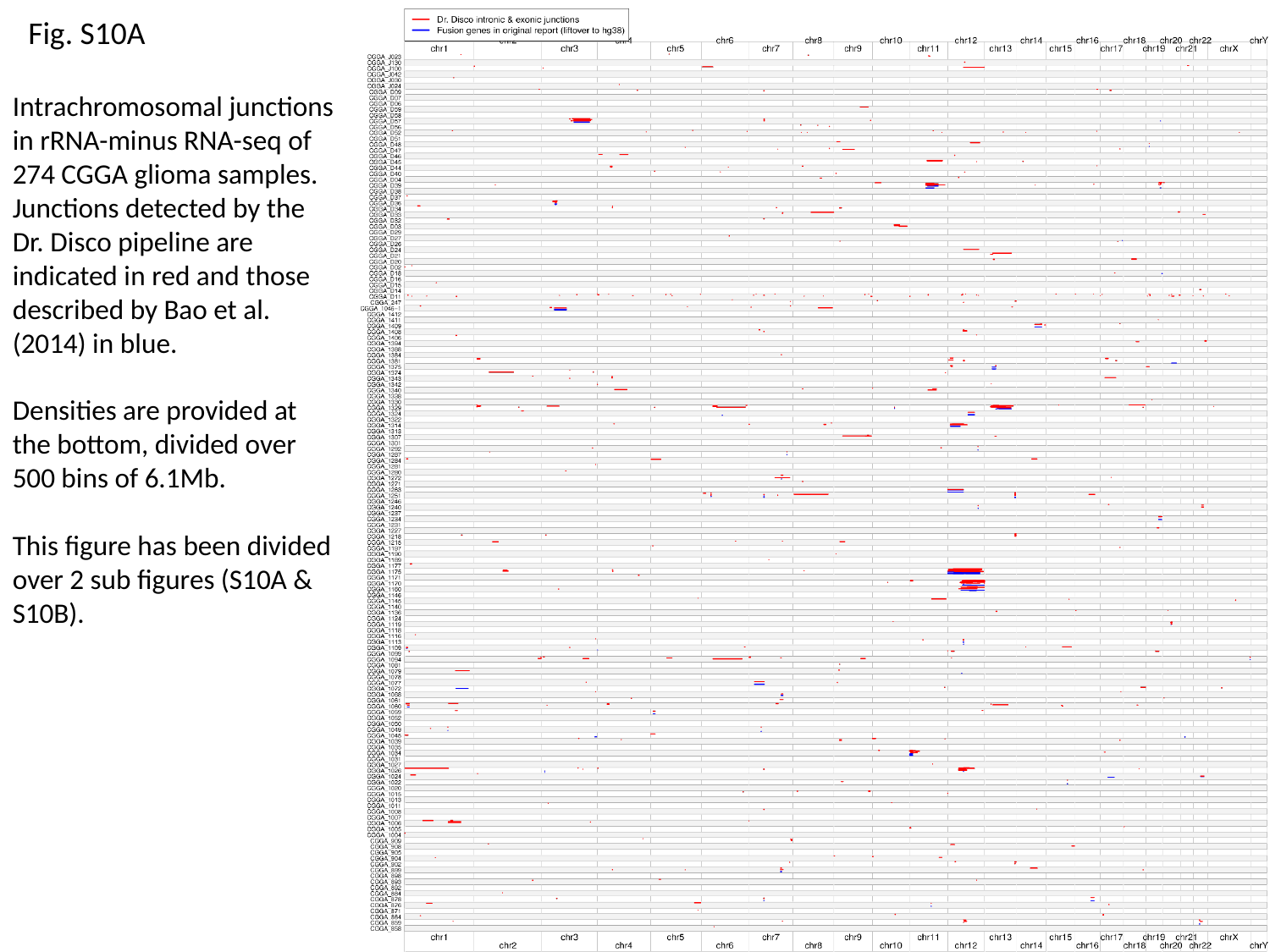

Fig. S10A
Intrachromosomal junctions in rRNA-minus RNA-seq of 274 CGGA glioma samples. Junctions detected by the Dr. Disco pipeline are indicated in red and those described by Bao et al. (2014) in blue.
Densities are provided at the bottom, divided over 500 bins of 6.1Mb.
This figure has been divided over 2 sub figures (S10A & S10B).

### Slide 15
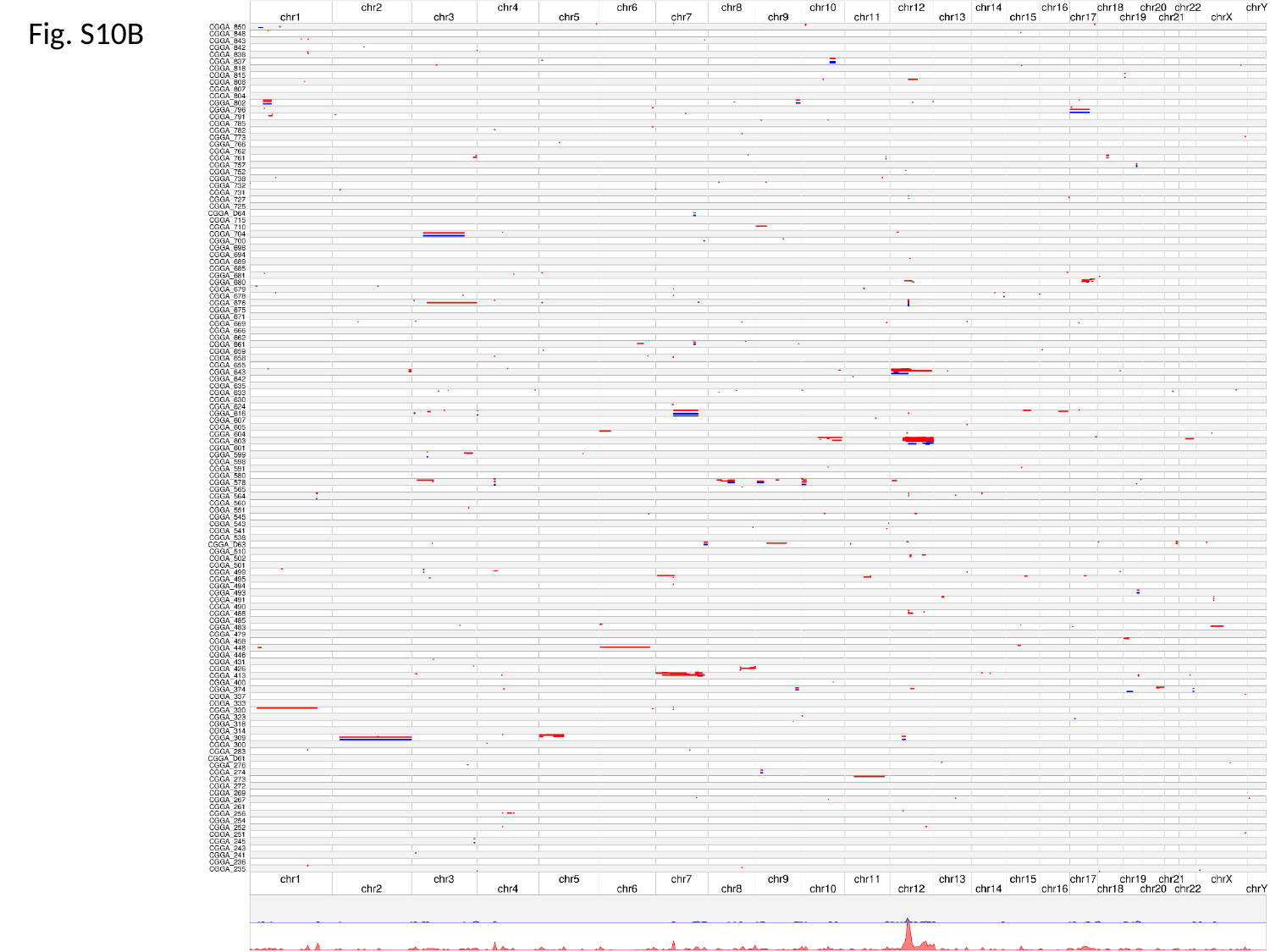

Fig. S10B

### Slide 16
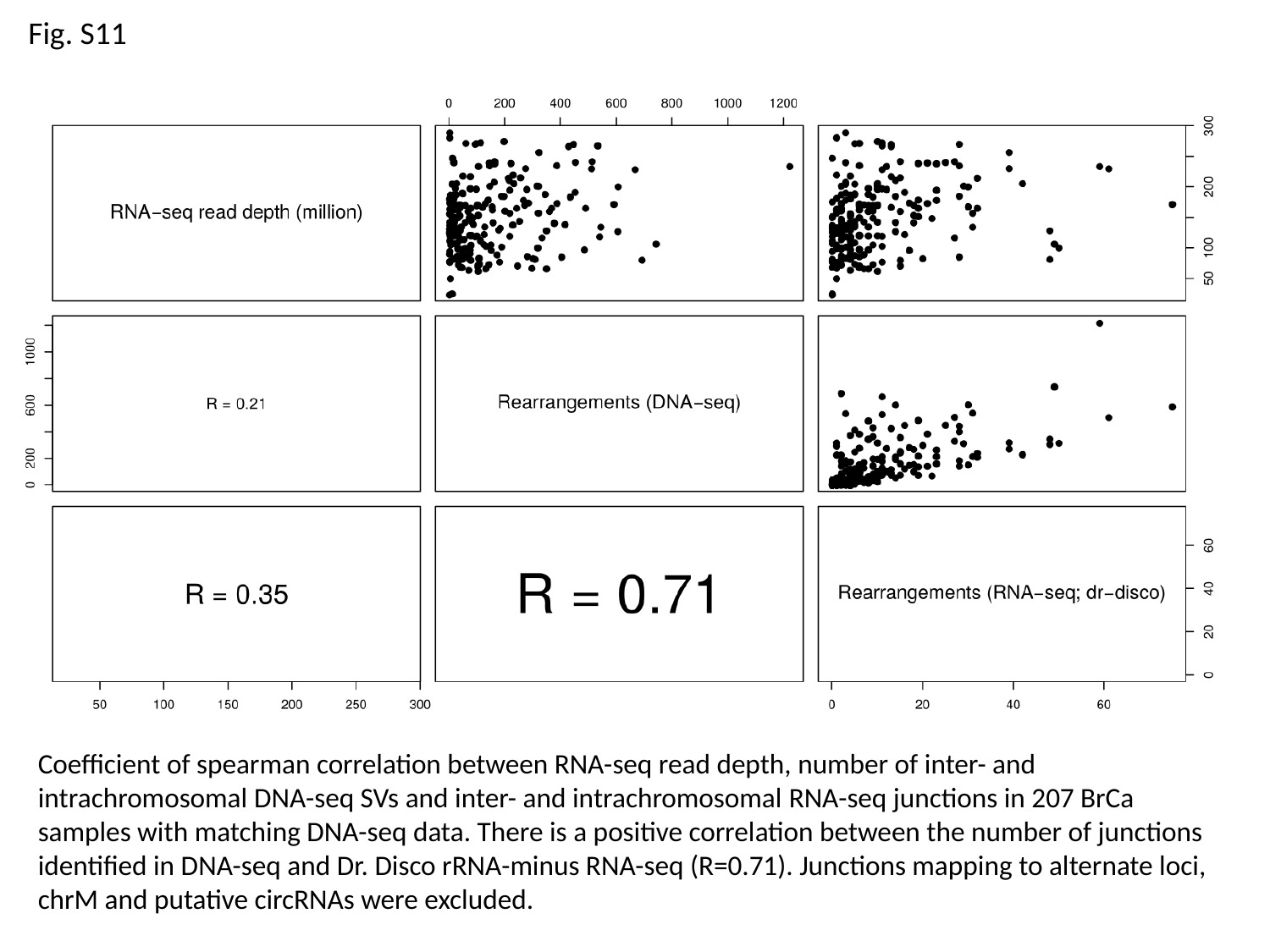

Fig. S11
Coefficient of spearman correlation between RNA-seq read depth, number of inter- and intrachromosomal DNA-seq SVs and inter- and intrachromosomal RNA-seq junctions in 207 BrCa samples with matching DNA-seq data. There is a positive correlation between the number of junctions identified in DNA-seq and Dr. Disco rRNA-minus RNA-seq (R=0.71). Junctions mapping to alternate loci, chrM and putative circRNAs were excluded.

### Slide 17
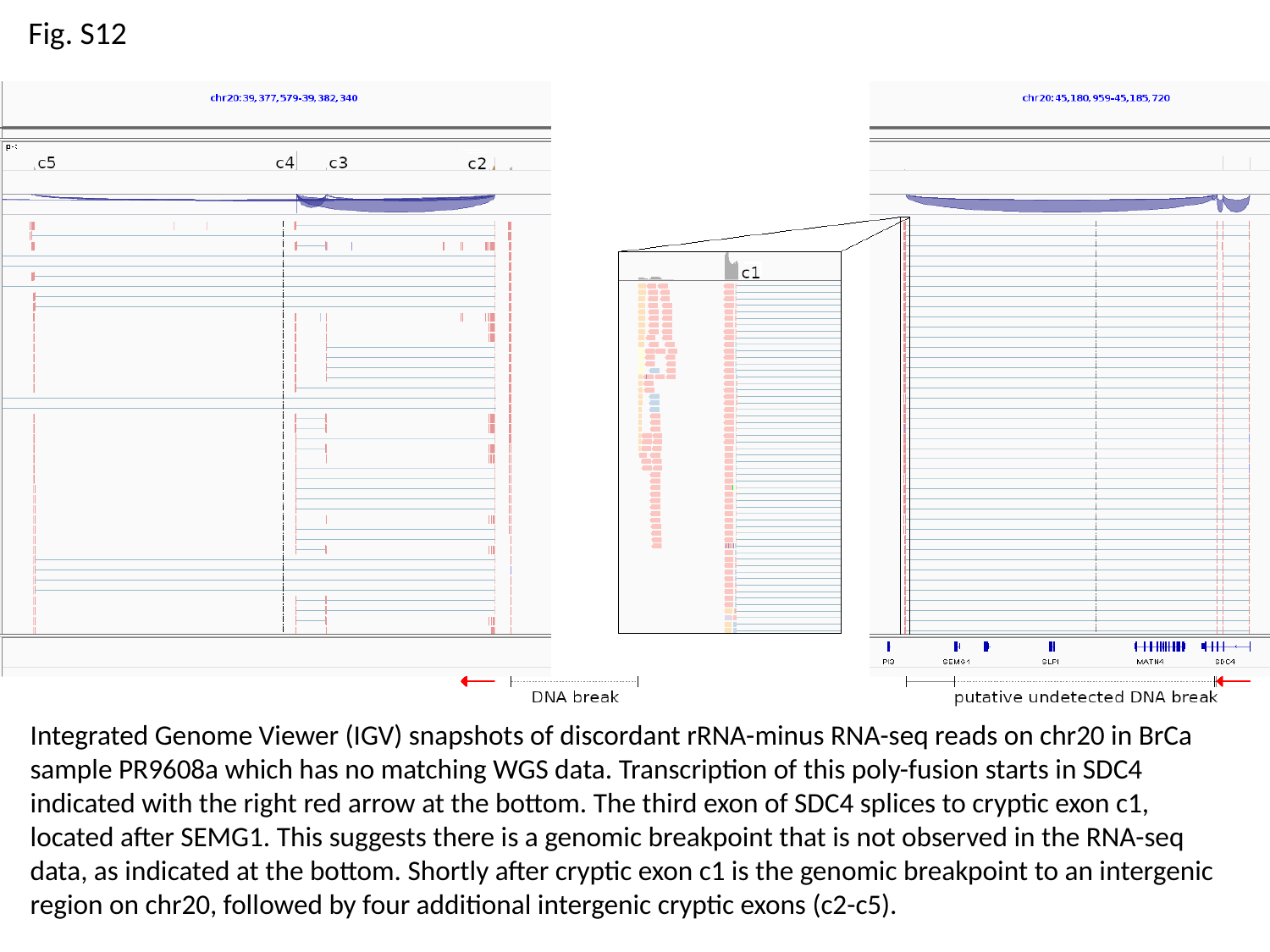

Fig. S12
Integrated Genome Viewer (IGV) snapshots of discordant rRNA-minus RNA-seq reads on chr20 in BrCa sample PR9608a which has no matching WGS data. Transcription of this poly-fusion starts in SDC4 indicated with the right red arrow at the bottom. The third exon of SDC4 splices to cryptic exon c1, located after SEMG1. This suggests there is a genomic breakpoint that is not observed in the RNA-seq data, as indicated at the bottom. Shortly after cryptic exon c1 is the genomic breakpoint to an intergenic region on chr20, followed by four additional intergenic cryptic exons (c2-c5).

### Slide 18
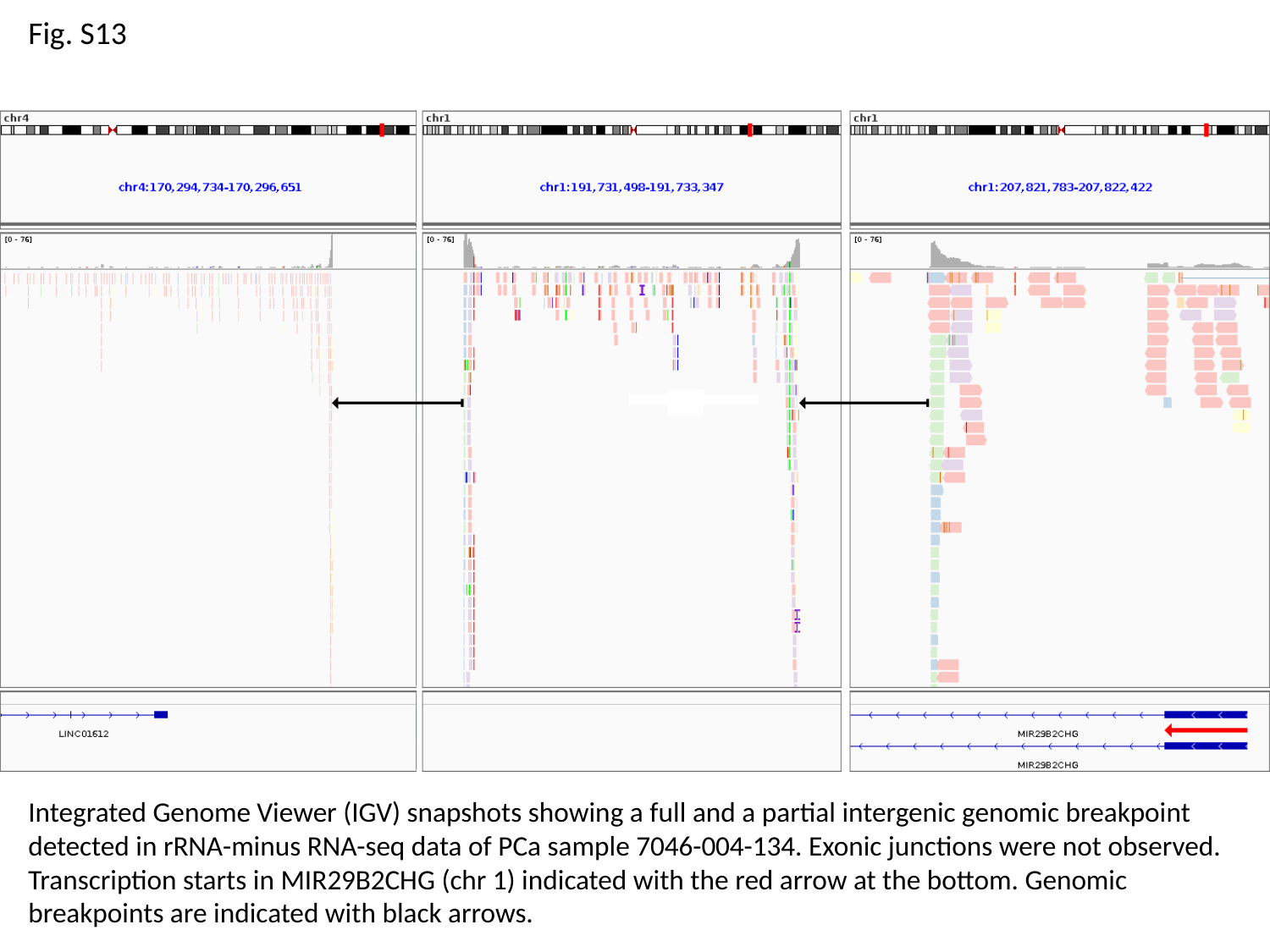

Fig. S13
Integrated Genome Viewer (IGV) snapshots showing a full and a partial intergenic genomic breakpoint detected in rRNA-minus RNA-seq data of PCa sample 7046-004-134. Exonic junctions were not observed. Transcription starts in MIR29B2CHG (chr 1) indicated with the red arrow at the bottom. Genomic breakpoints are indicated with black arrows.

### Slide 19
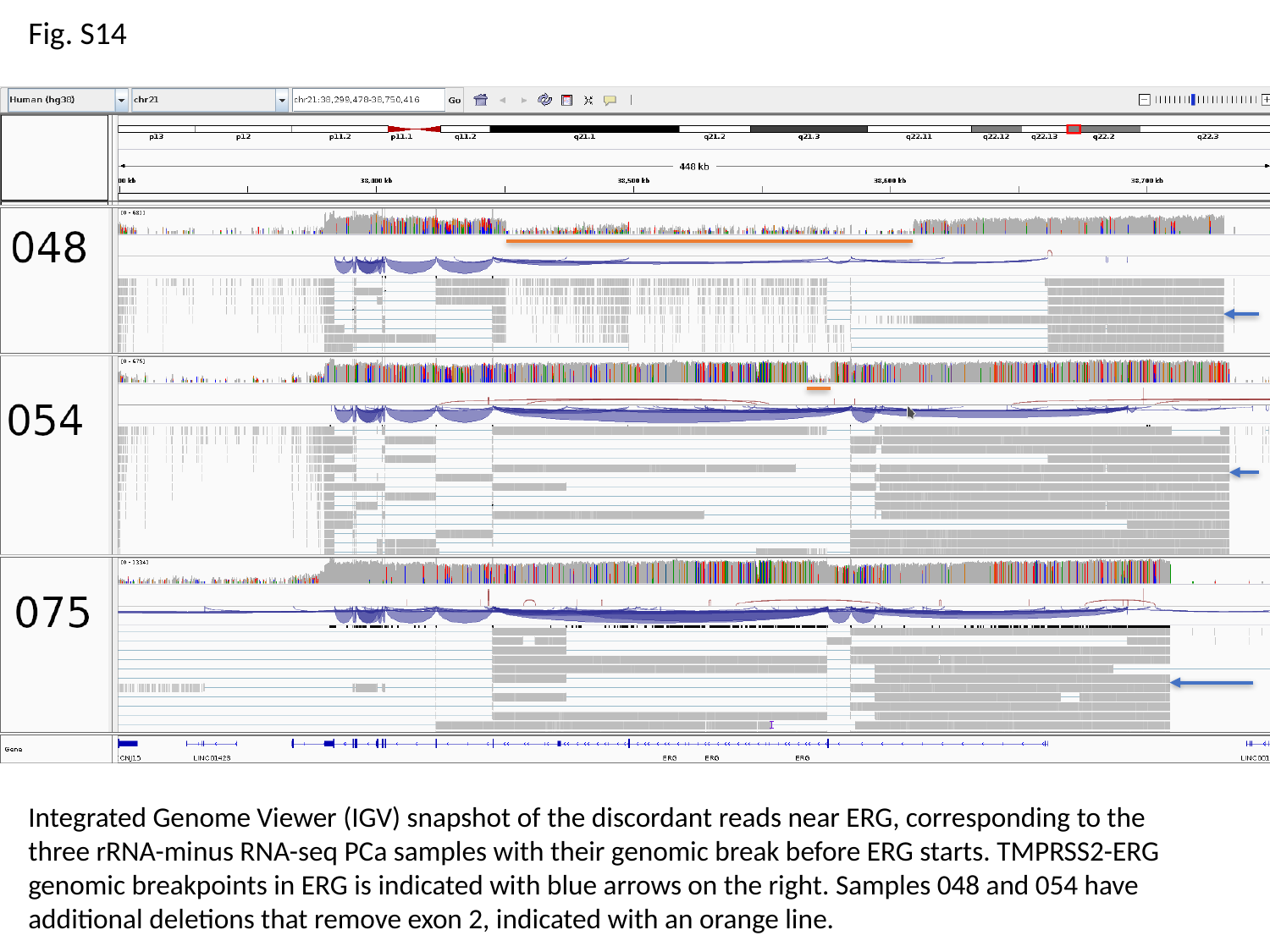

Fig. S14
Integrated Genome Viewer (IGV) snapshot of the discordant reads near ERG, corresponding to the three rRNA-minus RNA-seq PCa samples with their genomic break before ERG starts. TMPRSS2-ERG genomic breakpoints in ERG is indicated with blue arrows on the right. Samples 048 and 054 have additional deletions that remove exon 2, indicated with an orange line.

### Slide 20
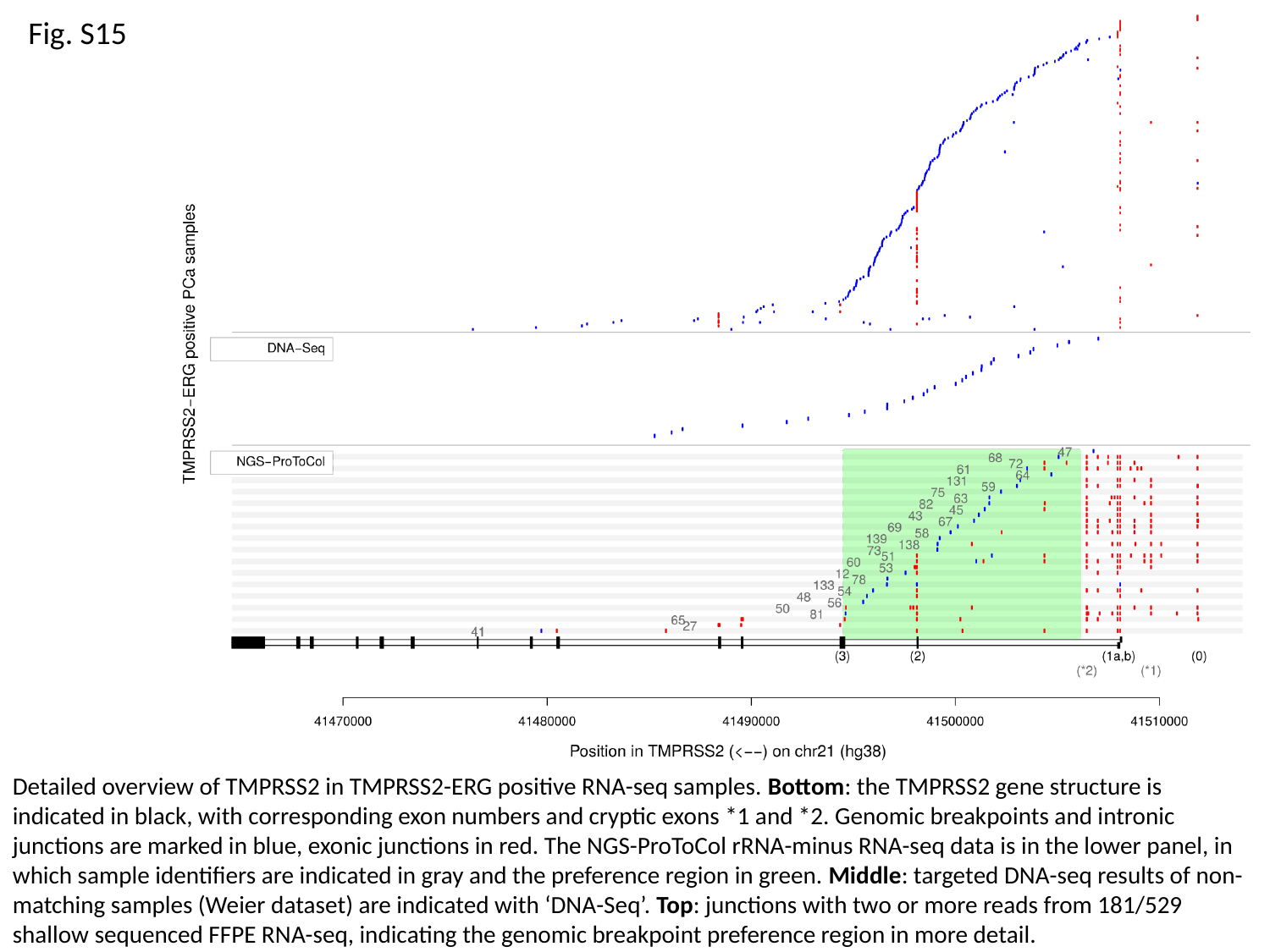

Fig. S15
Detailed overview of TMPRSS2 in TMPRSS2-ERG positive RNA-seq samples. Bottom: the TMPRSS2 gene structure is indicated in black, with corresponding exon numbers and cryptic exons *1 and *2. Genomic breakpoints and intronic junctions are marked in blue, exonic junctions in red. The NGS-ProToCol rRNA-minus RNA-seq data is in the lower panel, in which sample identifiers are indicated in gray and the preference region in green. Middle: targeted DNA-seq results of non-matching samples (Weier dataset) are indicated with ‘DNA-Seq’. Top: junctions with two or more reads from 181/529 shallow sequenced FFPE RNA-seq, indicating the genomic breakpoint preference region in more detail.

### Slide 21
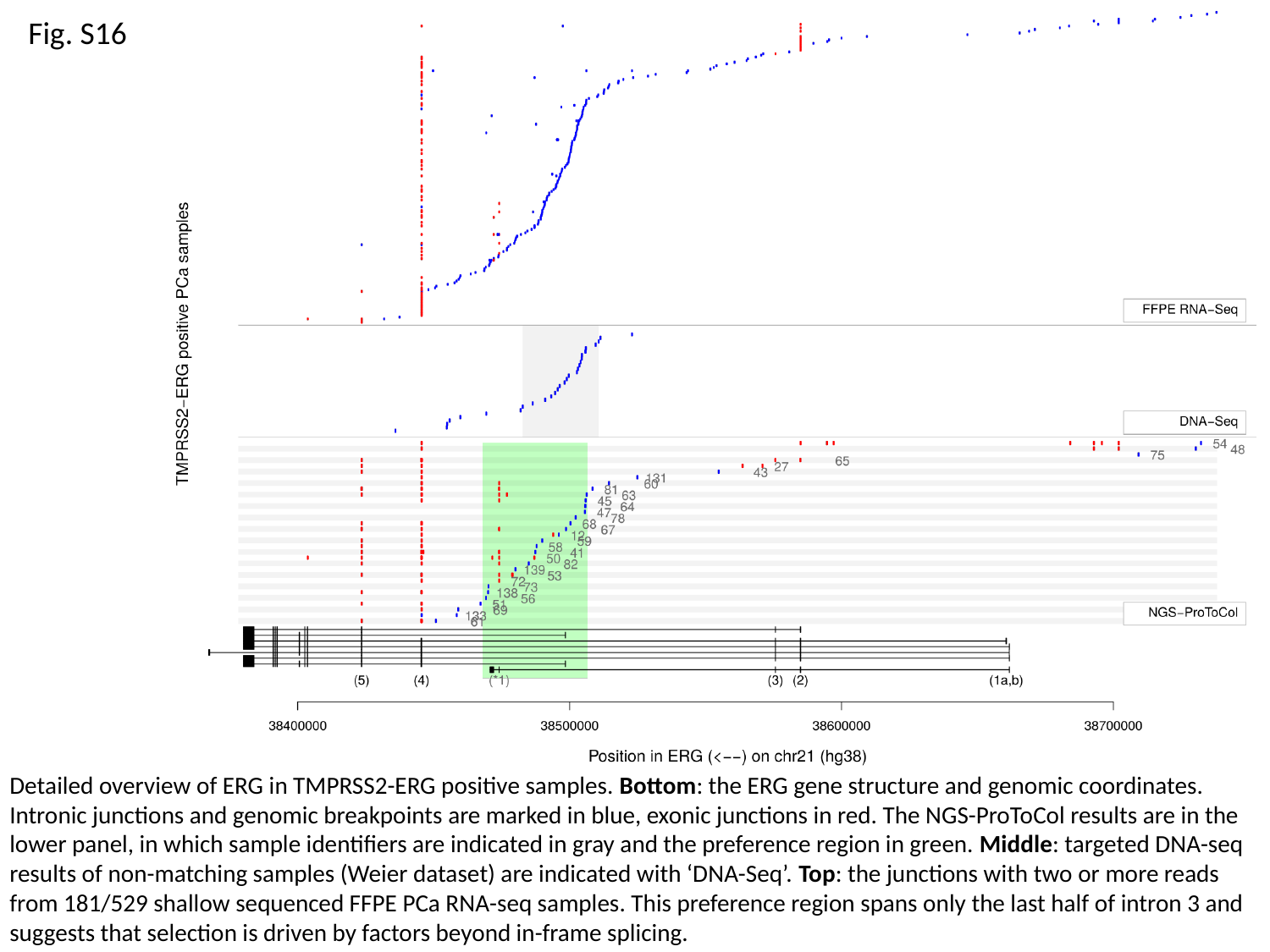

Fig. S16
Detailed overview of ERG in TMPRSS2-ERG positive samples. Bottom: the ERG gene structure and genomic coordinates. Intronic junctions and genomic breakpoints are marked in blue, exonic junctions in red. The NGS-ProToCol results are in the lower panel, in which sample identifiers are indicated in gray and the preference region in green. Middle: targeted DNA-seq results of non-matching samples (Weier dataset) are indicated with ‘DNA-Seq’. Top: the junctions with two or more reads from 181/529 shallow sequenced FFPE PCa RNA-seq samples. This preference region spans only the last half of intron 3 and suggests that selection is driven by factors beyond in-frame splicing.

### Slide 22
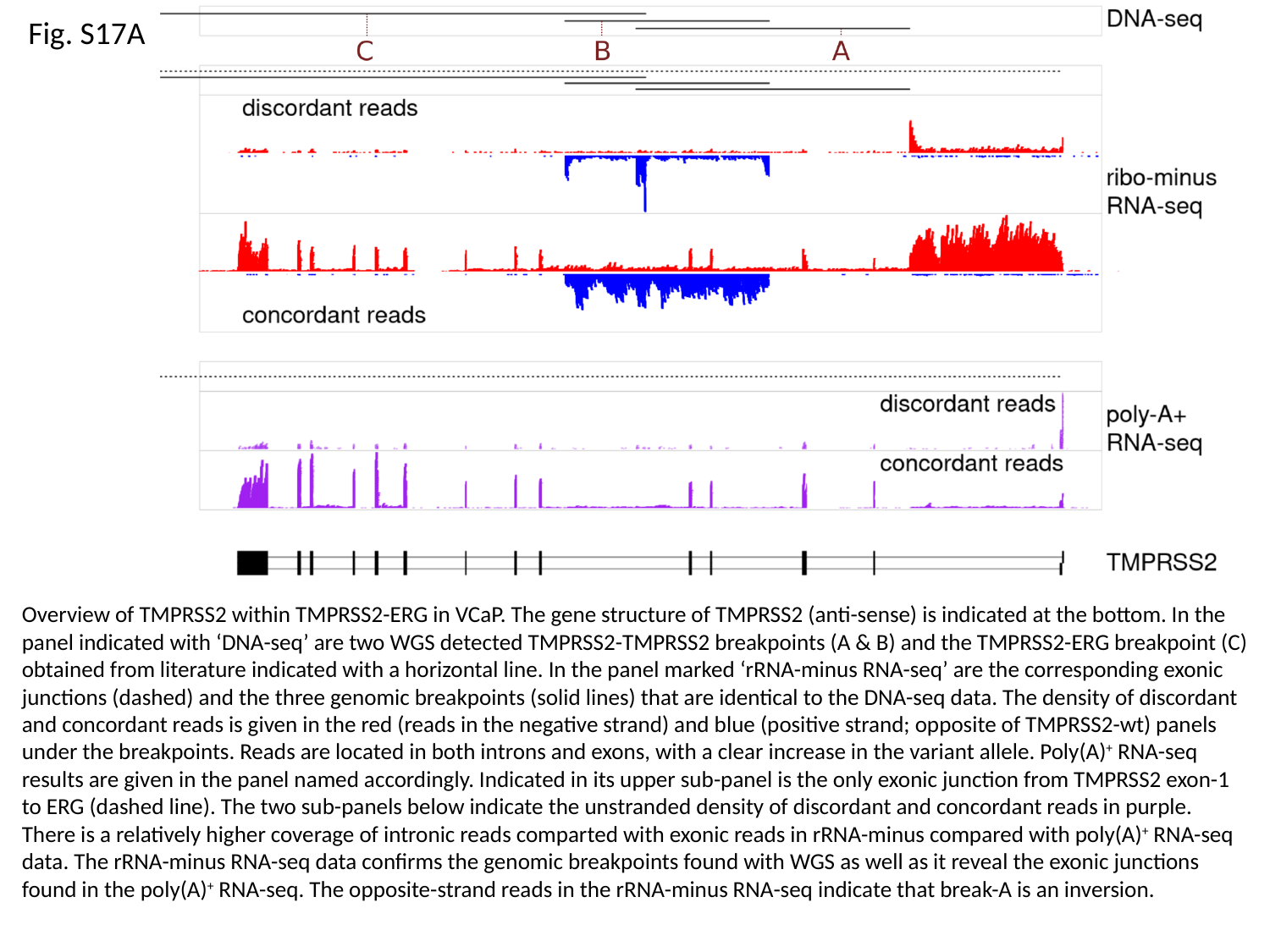

Fig. S17A
Overview of TMPRSS2 within TMPRSS2-ERG in VCaP. The gene structure of TMPRSS2 (anti-sense) is indicated at the bottom. In the panel indicated with ‘DNA-seq’ are two WGS detected TMPRSS2-TMPRSS2 breakpoints (A & B) and the TMPRSS2-ERG breakpoint (C) obtained from literature indicated with a horizontal line. In the panel marked ‘rRNA-minus RNA-seq’ are the corresponding exonic junctions (dashed) and the three genomic breakpoints (solid lines) that are identical to the DNA-seq data. The density of discordant and concordant reads is given in the red (reads in the negative strand) and blue (positive strand; opposite of TMPRSS2-wt) panels under the breakpoints. Reads are located in both introns and exons, with a clear increase in the variant allele. Poly(A)^+^ RNA-seq results are given in the panel named accordingly. Indicated in its upper sub-panel is the only exonic junction from TMPRSS2 exon-1 to ERG (dashed line). The two sub-panels below indicate the unstranded density of discordant and concordant reads in purple. There is a relatively higher coverage of intronic reads comparted with exonic reads in rRNA-minus compared with poly(A)^+^ RNA-seq data. The rRNA-minus RNA-seq data confirms the genomic breakpoints found with WGS as well as it reveal the exonic junctions found in the poly(A)^+^ RNA-seq. The opposite-strand reads in the rRNA-minus RNA-seq indicate that break-A is an inversion.

### Slide 23
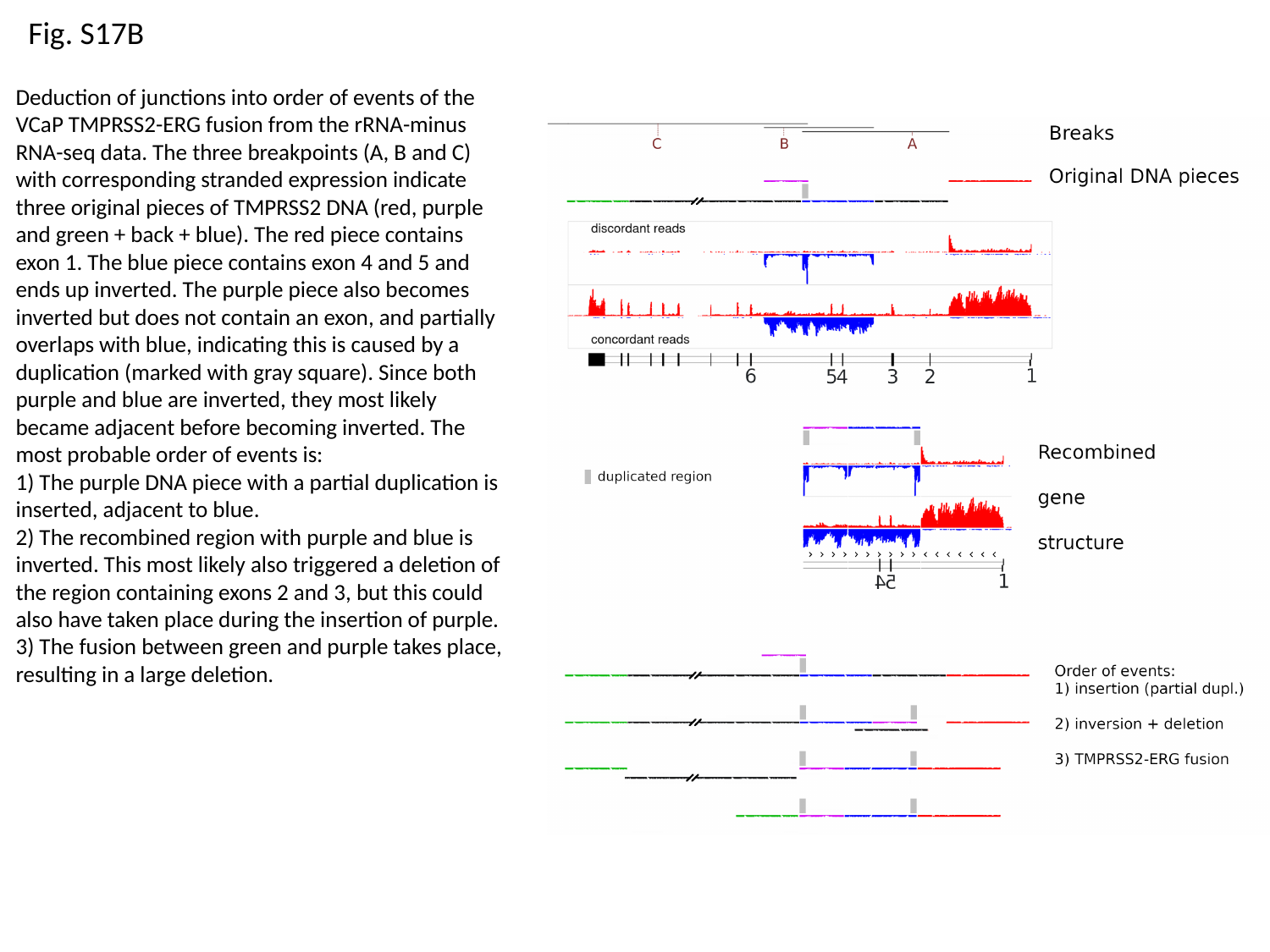

Fig. S17B
Deduction of junctions into order of events of the VCaP TMPRSS2-ERG fusion from the rRNA-minus RNA-seq data. The three breakpoints (A, B and C) with corresponding stranded expression indicate three original pieces of TMPRSS2 DNA (red, purple and green + back + blue). The red piece contains exon 1. The blue piece contains exon 4 and 5 and ends up inverted. The purple piece also becomes inverted but does not contain an exon, and partially overlaps with blue, indicating this is caused by a duplication (marked with gray square). Since both purple and blue are inverted, they most likely became adjacent before becoming inverted. The most probable order of events is:
1) The purple DNA piece with a partial duplication is inserted, adjacent to blue.
2) The recombined region with purple and blue is inverted. This most likely also triggered a deletion of the region containing exons 2 and 3, but this could also have taken place during the insertion of purple.
3) The fusion between green and purple takes place, resulting in a large deletion.

### Slide 24
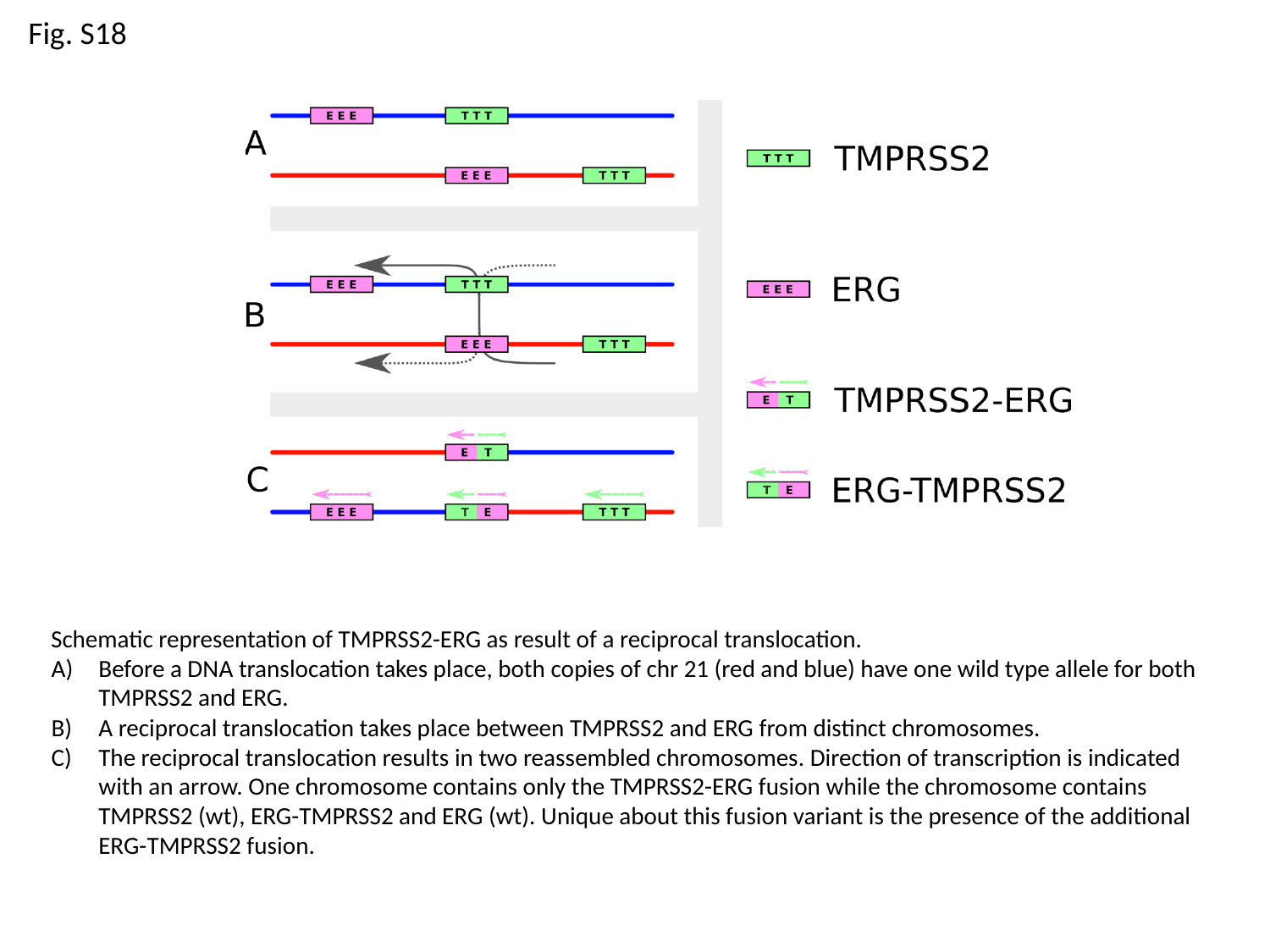

Fig. S18
Schematic representation of TMPRSS2-ERG as result of a reciprocal translocation.
Before a DNA translocation takes place, both copies of chr 21 (red and blue) have one wild type allele for both TMPRSS2 and ERG.
A reciprocal translocation takes place between TMPRSS2 and ERG from distinct chromosomes.
The reciprocal translocation results in two reassembled chromosomes. Direction of transcription is indicated with an arrow. One chromosome contains only the TMPRSS2-ERG fusion while the chromosome contains TMPRSS2 (wt), ERG-TMPRSS2 and ERG (wt). Unique about this fusion variant is the presence of the additional ERG-TMPRSS2 fusion.

### Slide 25
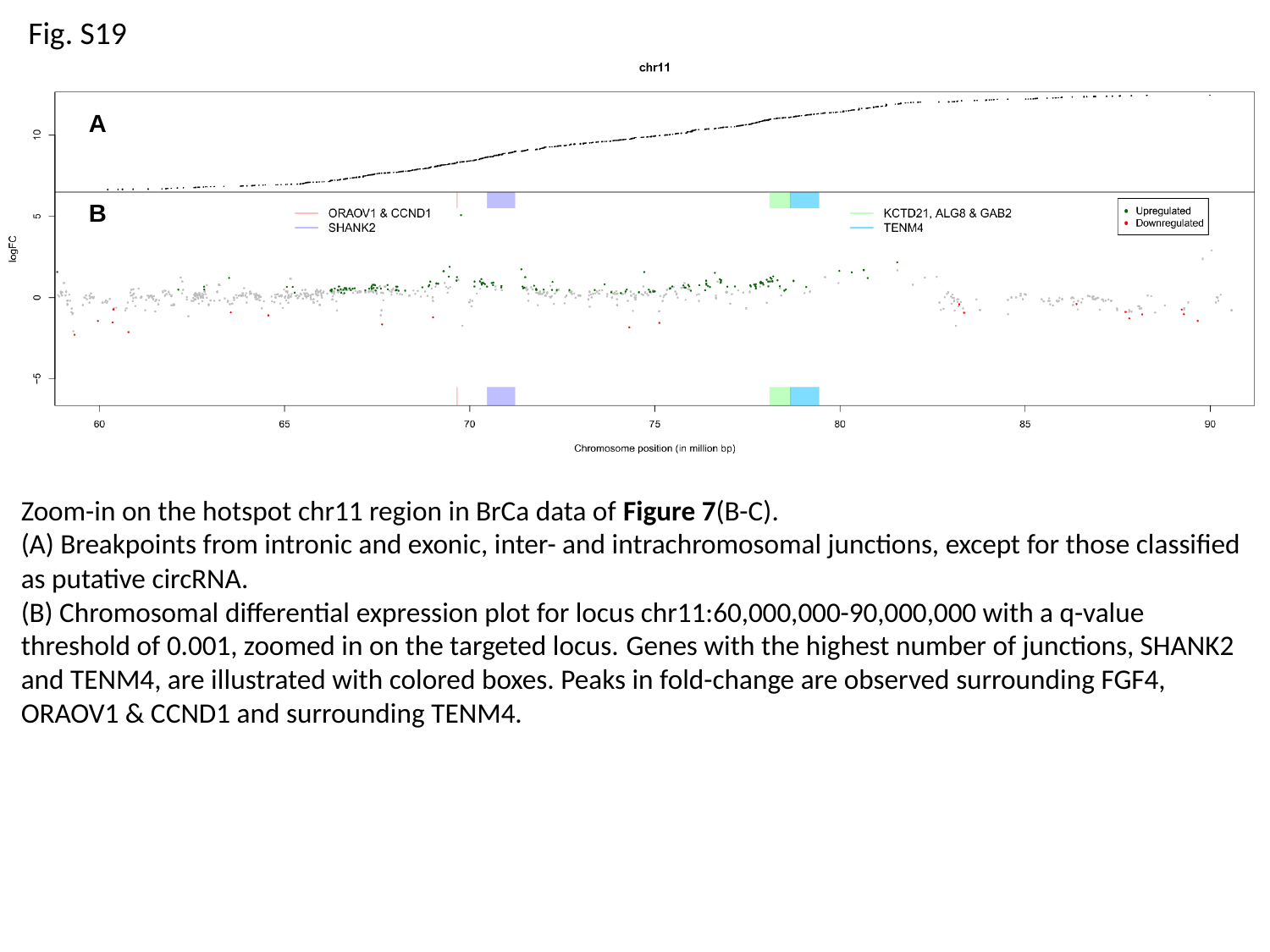

Fig. S19
A
B
Zoom-in on the hotspot chr11 region in BrCa data of Figure 7(B-C).
(A) Breakpoints from intronic and exonic, inter- and intrachromosomal junctions, except for those classified as putative circRNA.
(B) Chromosomal differential expression plot for locus chr11:60,000,000-90,000,000 with a q-value threshold of 0.001, zoomed in on the targeted locus. Genes with the highest number of junctions, SHANK2 and TENM4, are illustrated with colored boxes. Peaks in fold-change are observed surrounding FGF4, ORAOV1 & CCND1 and surrounding TENM4.

### Slide 26
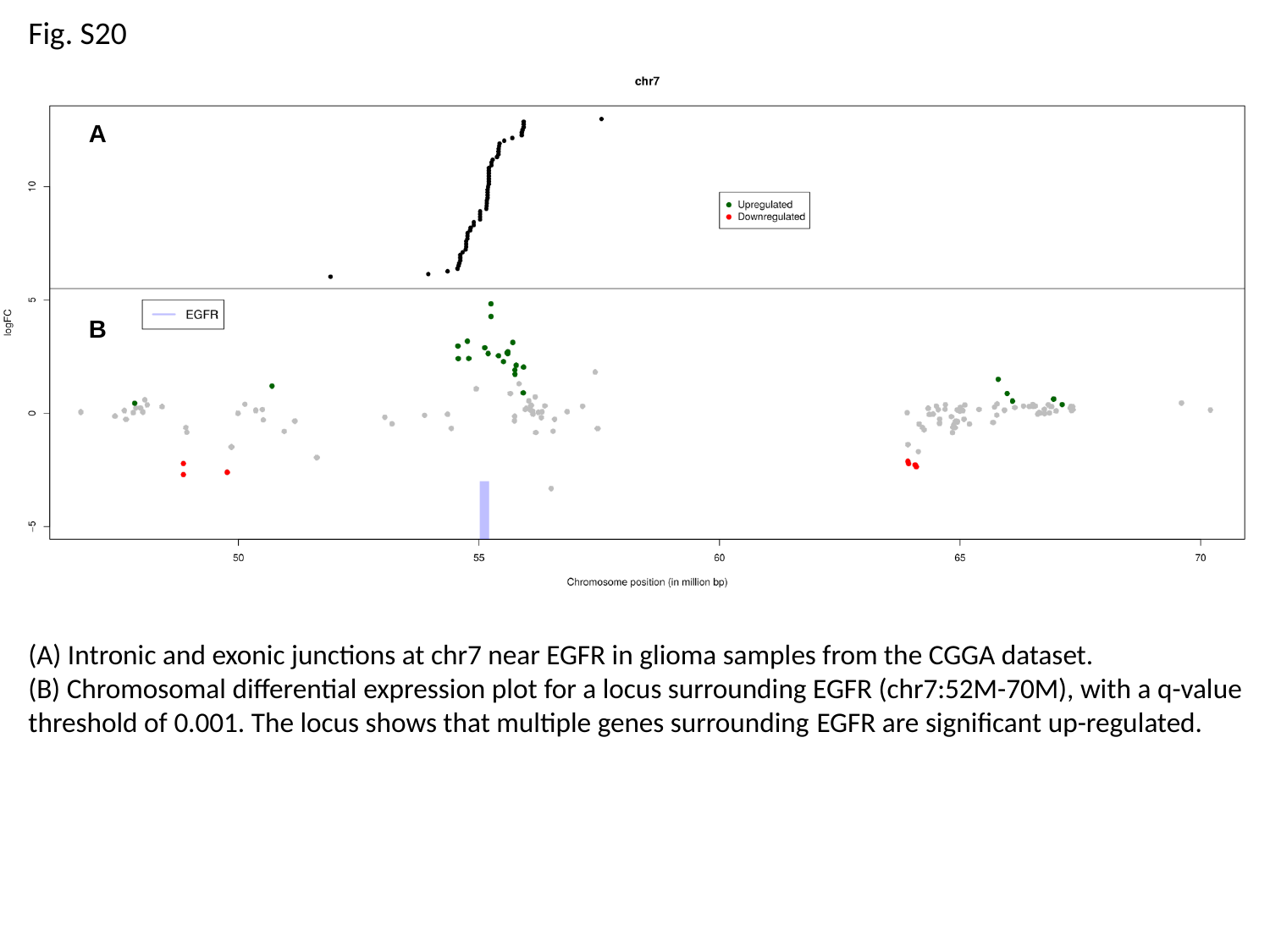

Fig. S20
A
B
(A) Intronic and exonic junctions at chr7 near EGFR in glioma samples from the CGGA dataset.
(B) Chromosomal differential expression plot for a locus surrounding EGFR (chr7:52M-70M), with a q-value threshold of 0.001. The locus shows that multiple genes surrounding EGFR are significant up-regulated.

### Slide 27
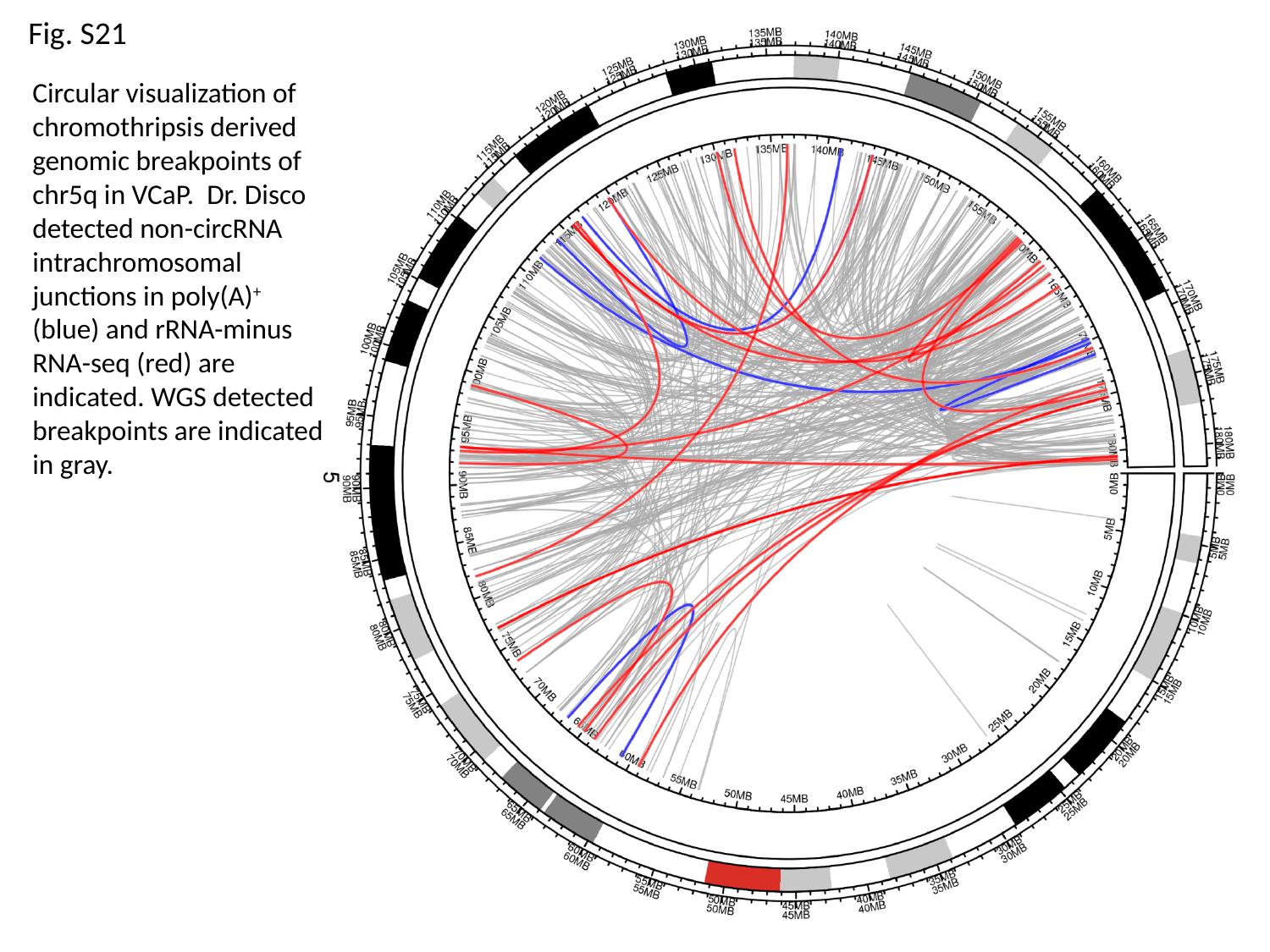

Fig. S21
Circular visualization of chromothripsis derived genomic breakpoints of chr5q in VCaP. Dr. Disco detected non-circRNA intrachromosomal junctions in poly(A)^+^ (blue) and rRNA-minus RNA-seq (red) are indicated. WGS detected breakpoints are indicated in gray.

### Slide 28
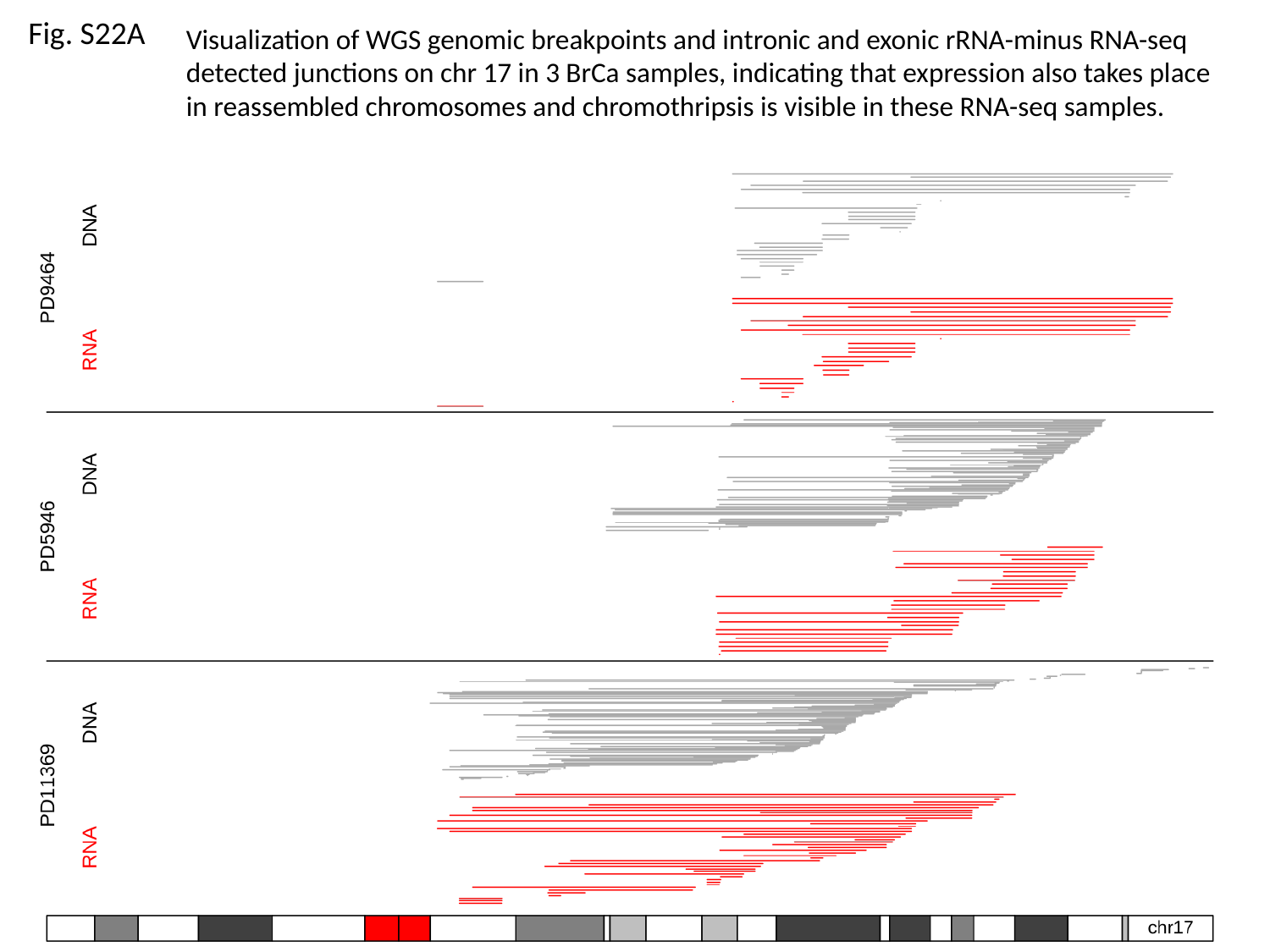

Fig. S22A
Visualization of WGS genomic breakpoints and intronic and exonic rRNA-minus RNA-seq detected junctions on chr 17 in 3 BrCa samples, indicating that expression also takes place in reassembled chromosomes and chromothripsis is visible in these RNA-seq samples.

### Slide 29
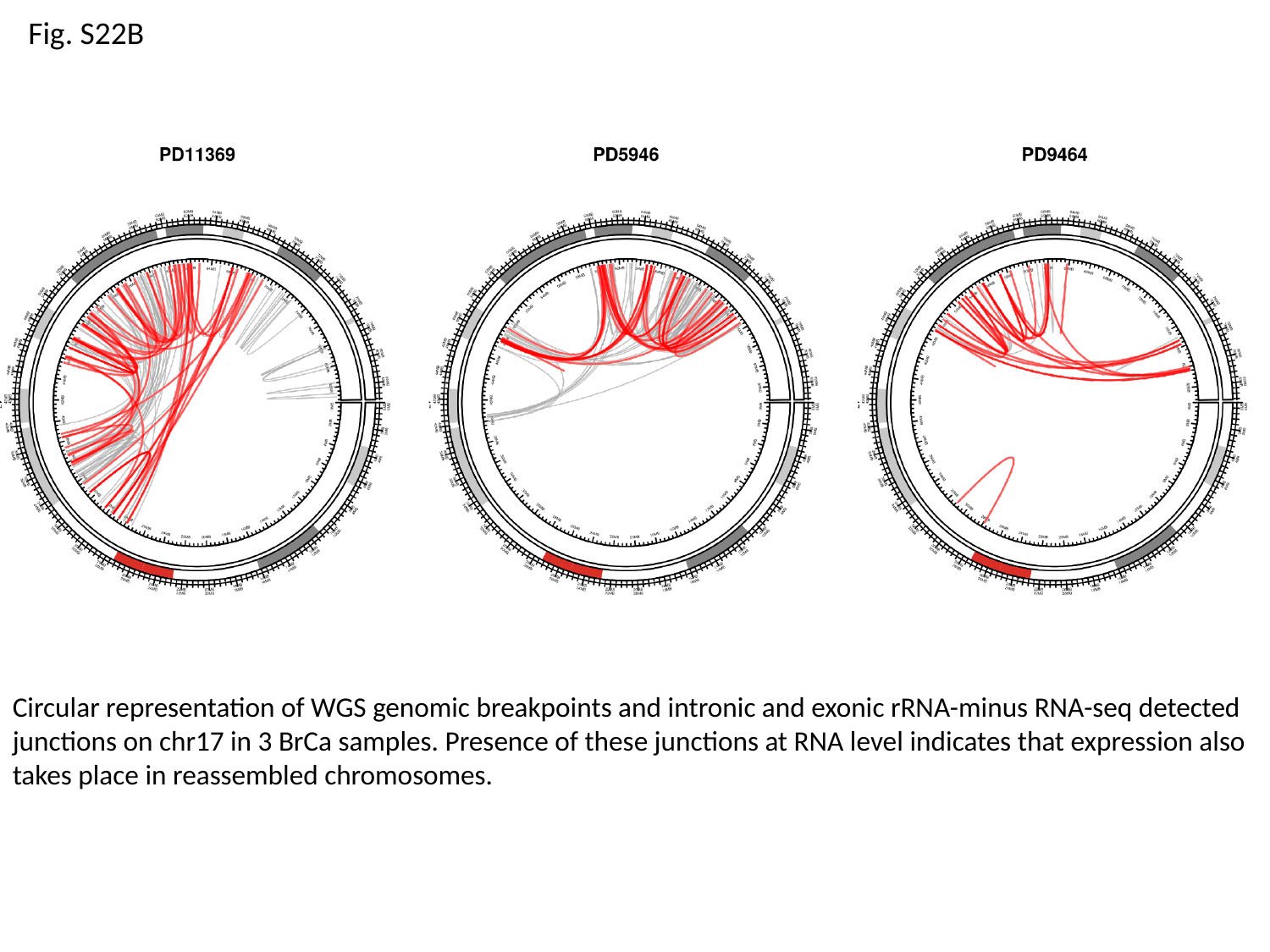

Fig. S22B
Circular representation of WGS genomic breakpoints and intronic and exonic rRNA-minus RNA-seq detected junctions on chr17 in 3 BrCa samples. Presence of these junctions at RNA level indicates that expression also takes place in reassembled chromosomes.

### Slide 30
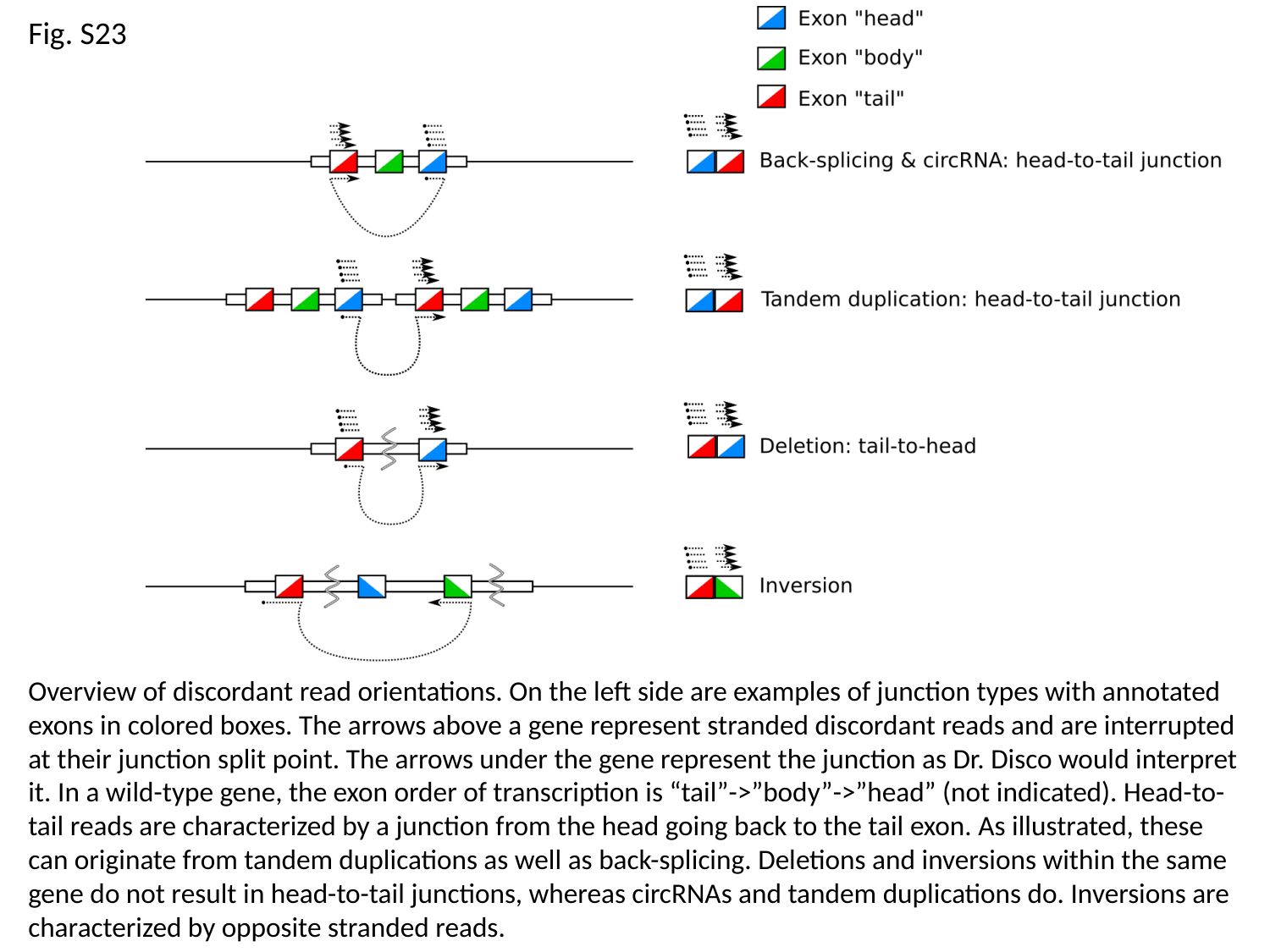

Fig. S23
Overview of discordant read orientations. On the left side are examples of junction types with annotated exons in colored boxes. The arrows above a gene represent stranded discordant reads and are interrupted at their junction split point. The arrows under the gene represent the junction as Dr. Disco would interpret it. In a wild-type gene, the exon order of transcription is “tail”->”body”->”head” (not indicated). Head-to-tail reads are characterized by a junction from the head going back to the tail exon. As illustrated, these can originate from tandem duplications as well as back-splicing. Deletions and inversions within the same gene do not result in head-to-tail junctions, whereas circRNAs and tandem duplications do. Inversions are characterized by opposite stranded reads.

### Slide 31
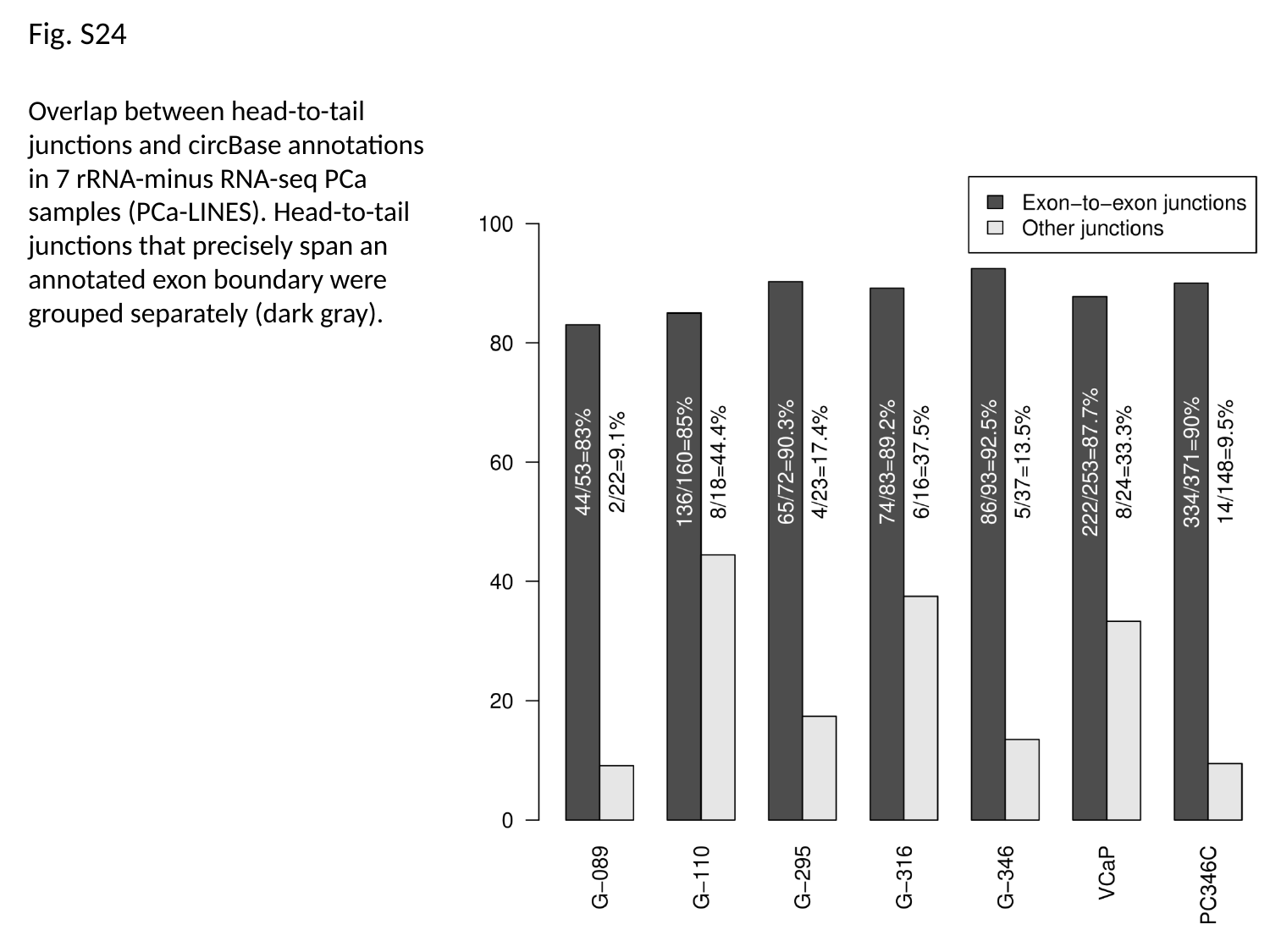

Fig. S24
Overlap between head-to-tail junctions and circBase annotations in 7 rRNA-minus RNA-seq PCa samples (PCa-LINES). Head-to-tail junctions that precisely span an annotated exon boundary were grouped separately (dark gray).

### Slide 32
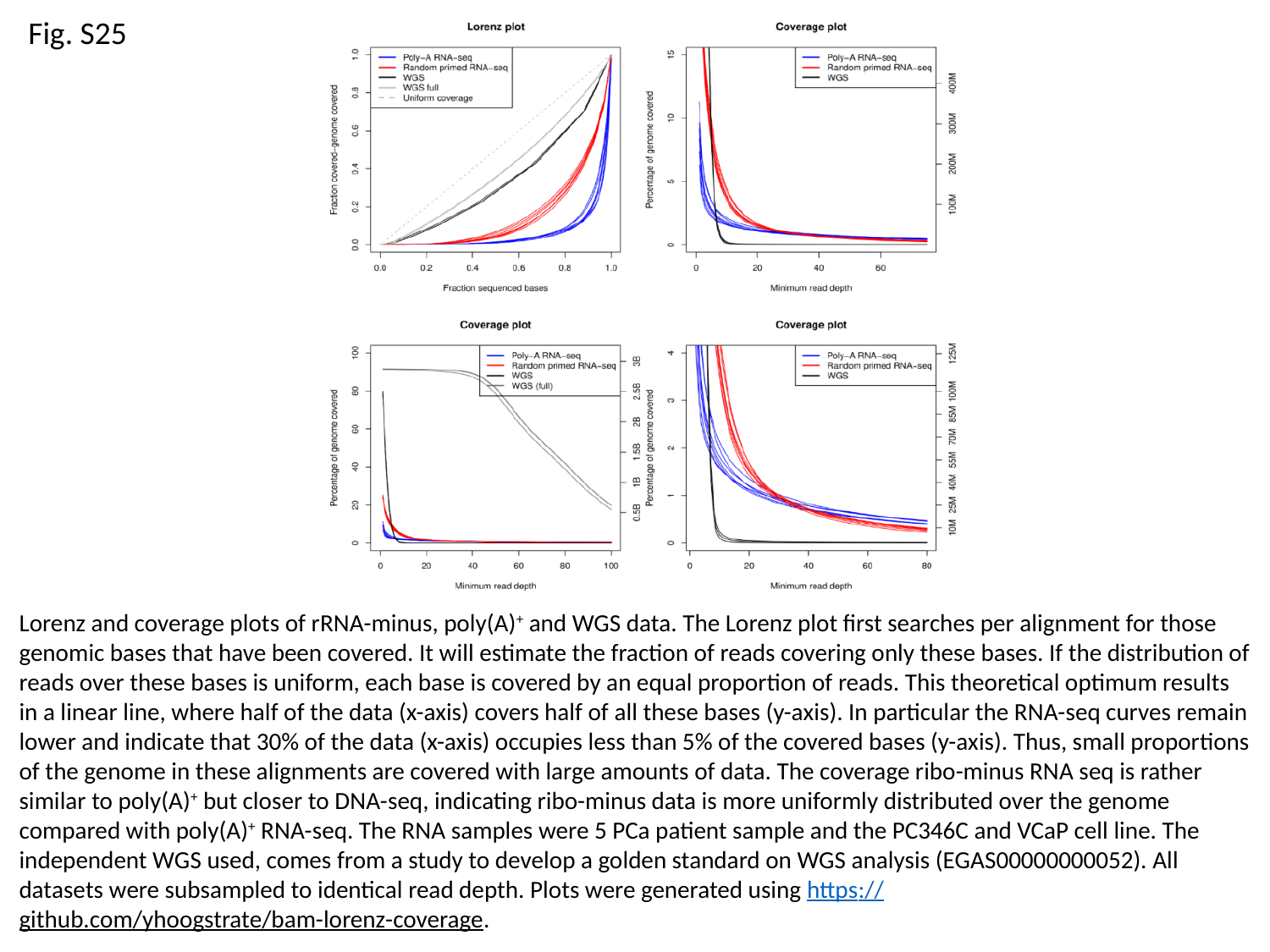

Fig. S25
Lorenz and coverage plots of rRNA-minus, poly(A)^+^ and WGS data. The Lorenz plot first searches per alignment for those genomic bases that have been covered. It will estimate the fraction of reads covering only these bases. If the distribution of reads over these bases is uniform, each base is covered by an equal proportion of reads. This theoretical optimum results in a linear line, where half of the data (x-axis) covers half of all these bases (y-axis). In particular the RNA-seq curves remain lower and indicate that 30% of the data (x-axis) occupies less than 5% of the covered bases (y-axis). Thus, small proportions of the genome in these alignments are covered with large amounts of data. The coverage ribo-minus RNA seq is rather similar to poly(A)^+^ but closer to DNA-seq, indicating ribo-minus data is more uniformly distributed over the genome compared with poly(A)^+^ RNA-seq. The RNA samples were 5 PCa patient sample and the PC346C and VCaP cell line. The independent WGS used, comes from a study to develop a golden standard on WGS analysis (EGAS00000000052). All datasets were subsampled to identical read depth. Plots were generated using https://github.com/yhoogstrate/bam-lorenz-coverage.

### Slide 33
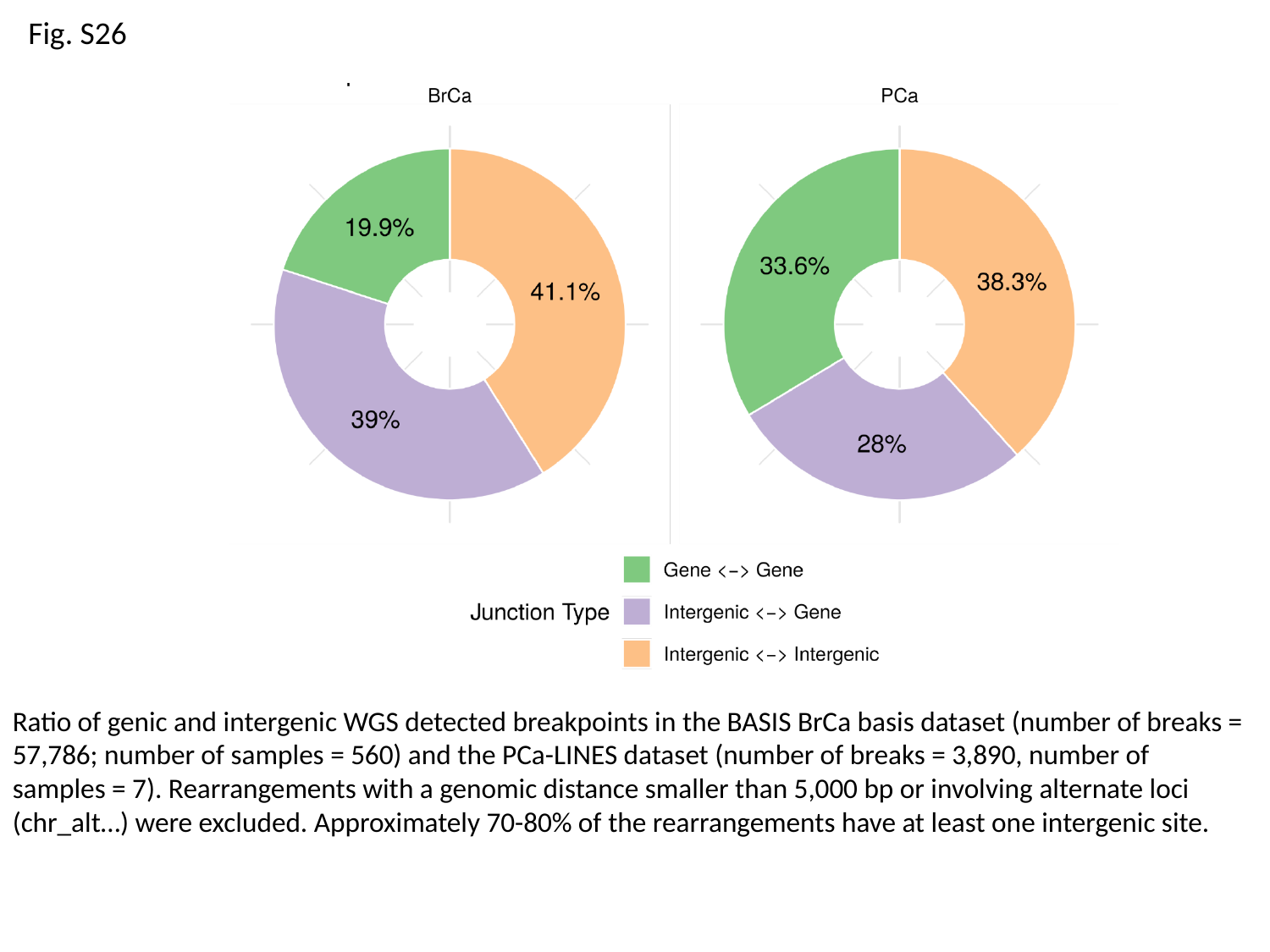

Fig. S26
Ratio of genic and intergenic WGS detected breakpoints in the BASIS BrCa basis dataset (number of breaks = 57,786; number of samples = 560) and the PCa-LINES dataset (number of breaks = 3,890, number of samples = 7). Rearrangements with a genomic distance smaller than 5,000 bp or involving alternate loci (chr_alt…) were excluded. Approximately 70-80% of the rearrangements have at least one intergenic site.
