## Supplementary figures and images for "Detection of fusion transcripts and their genomic breakpoints from RNA sequencing data"

### Table S2 - Recurrent Fusion Genes

#### Recurrent fusion genes

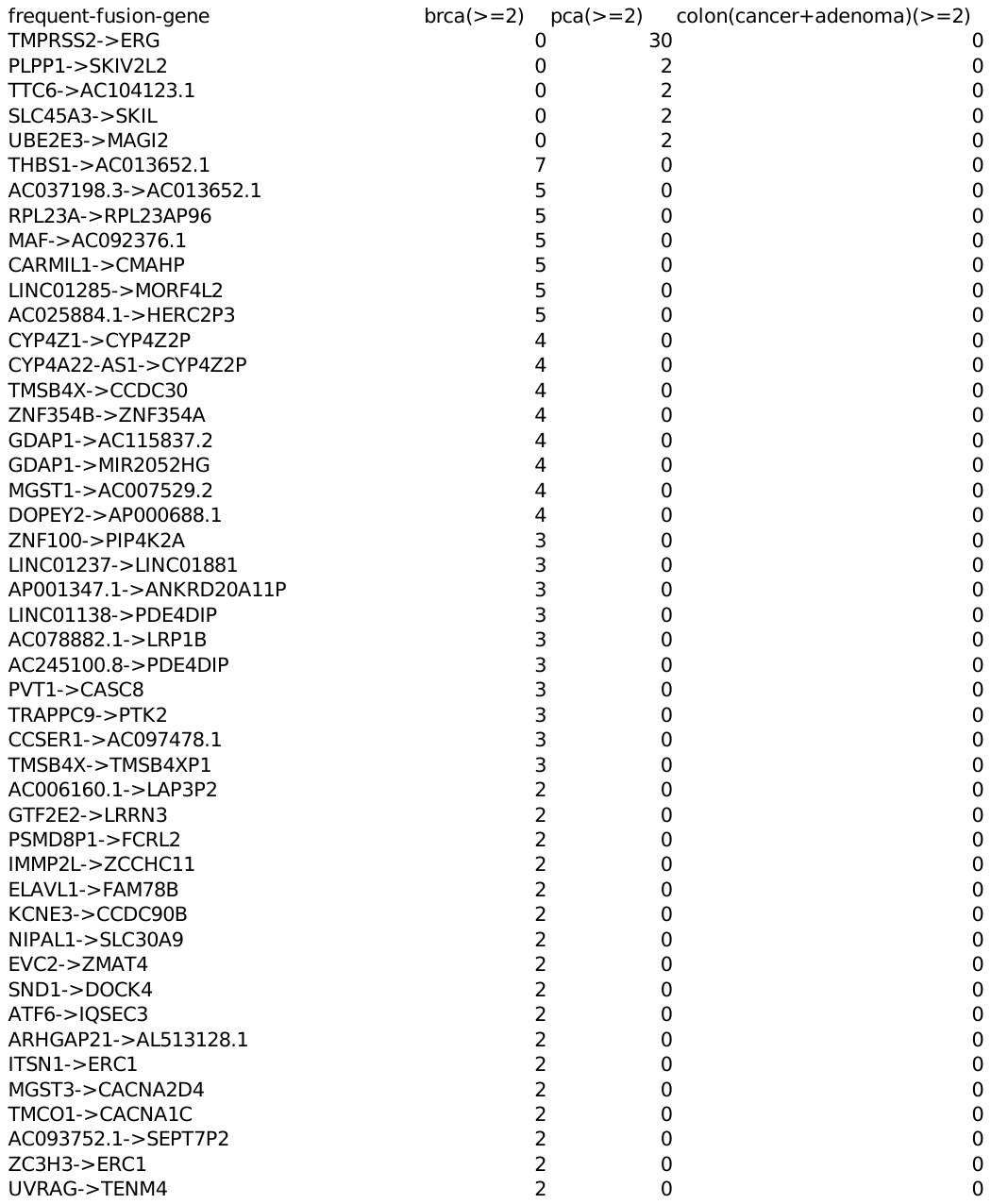

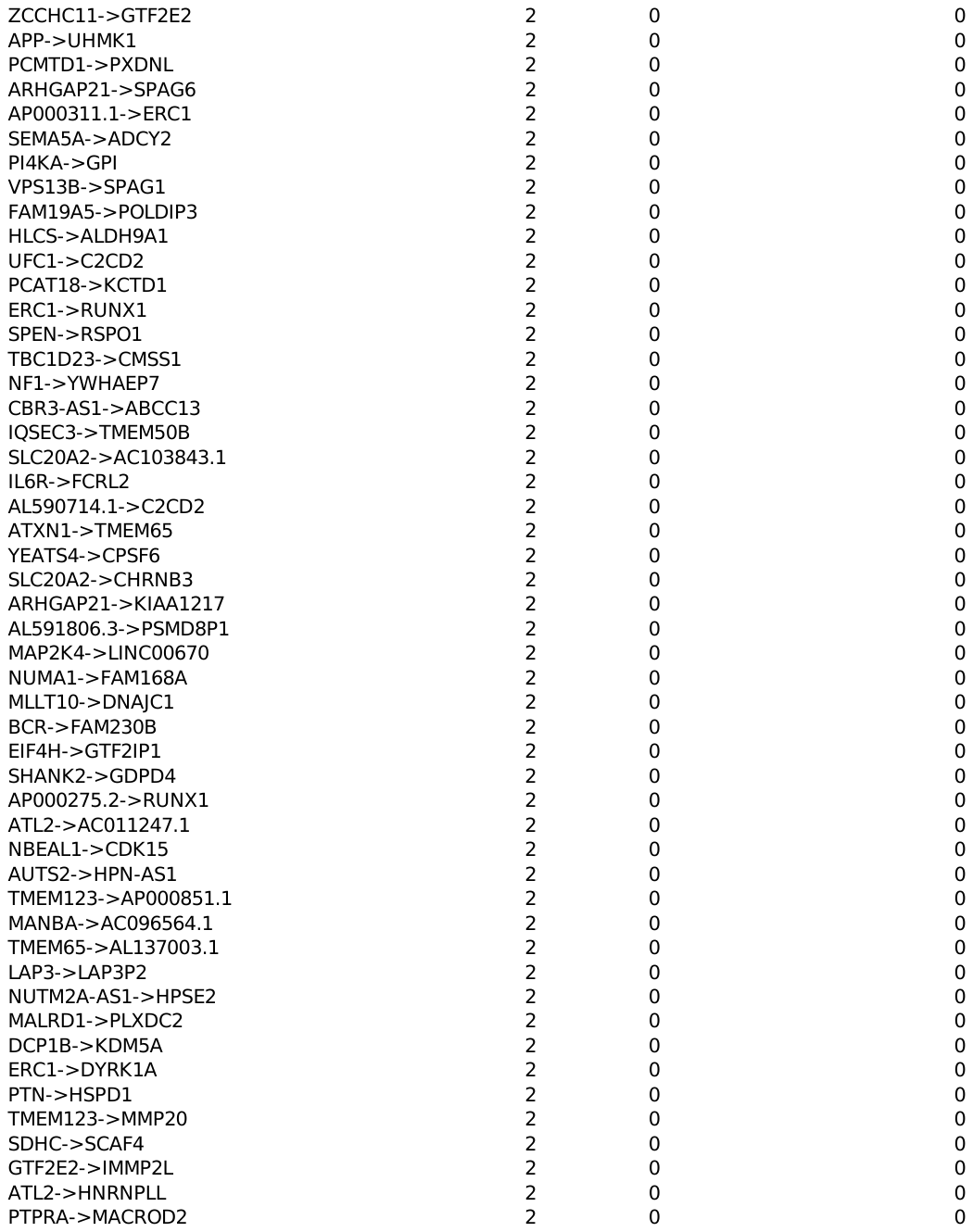

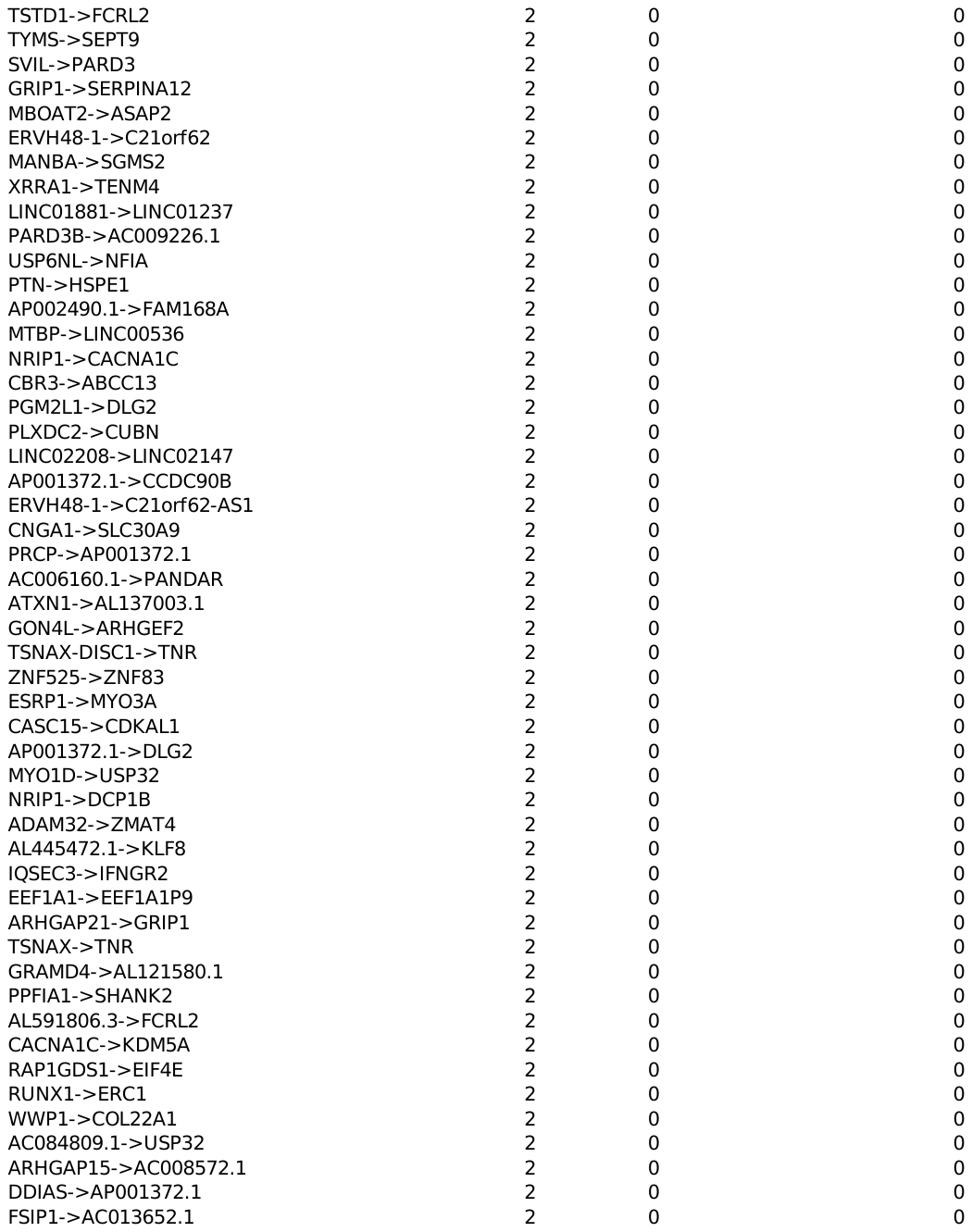

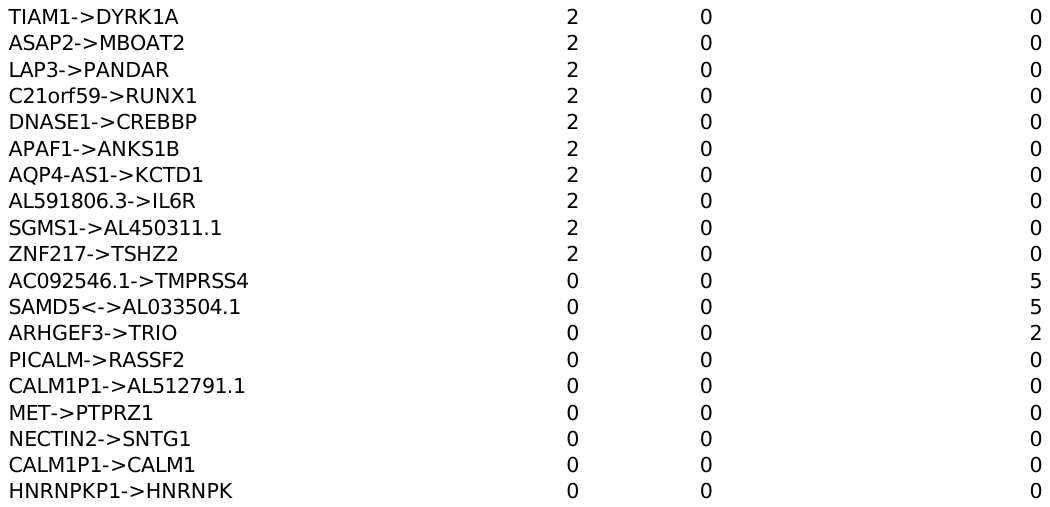
