## Supplementary Methods for "Detection of fusion transcripts and their genomic breakpoints from RNA sequencing data"

Data analysis

### Library prep

### Main Computational pipeline

The following datasets were analysed with the exact same pipeline described below:

- NGS ProToCol
- polyA+ cell lines
- glioma / SRR934749

The raw sequencing reads were aligned with STAR 2.4.2 (static build) to hg38 provided by UCSC. The following settings were used:

--outSAMtype BAM SortedByCoordinate

--alignEndsType Local

--alignIntronMax 200000

--alignMatesGapMax 200000

--alignSJDBoverhangMin 10

--chimJunctionOverhangMin 12

--chimSegmentMin 12

--outFilterIntronMotifs None

--outSAMstrandField intronMotif

--twopass1readsN -1

--twopassMode Basic

The concordant and discordant aligned reads are separated (Aligned.sortedByCoord.out.bam and Chimeric.out.sam, respectively). These chimeric (discordant) sam files were used as input for dr-disco version 0.17.8. The pipeline was performed as follows:

dr-disco fix

dr-disco detect -m 8

dr-disco classify

--blacklist-regions 'bin/drdisco/share/blacklist-regions.hg38.bed'

--blacklist-junctions 'bin/drdisco/share/blacklist-junctions.hg38.txt' --only-valid

dr-disco integrate

--fasta 'gtffiles_hg38/hg38.fa'

--gtf 'gtffiles_hg38/Ensemble.Homo_sapiens.GRCh38.89.gtf'

Where the blacklist files are shipped with dr-disco, the genome reference is from UCSC and the gene reference from Ensemble.

#### Ribo-minus prostate cell lines (HMG5MBBXX)

This dataset is rich in duplicate reads. To reduce this, we collapsed duplicate reads. Currently available tools cannot do this appropriately on discordant alignments as they do not take multiple aligned segments properly into account. We therefore collapsed duplicate reads at fastq level, using:

hts_SuperDeduper -f -e 1000000

These FASTQ files were then used for the main pipeline.

The tool is part of the following software package: <https://github.com/ibest/HTStream>

#### Ribo-minus FFPE / low-q

Because FFPE material is often degraded, the insert sizes may be smaller than the read length and additional adapter sequences may be present. Therefore, this dataset was first adapter cleaned with ‘sickle’ v1.33:

sickle pe -q 10 -q 1 -t sanger -l 40 -n

These FASTQ files were then used for the main pipeline.

The dr-disco classify was performed without `--only-valid` and with `--ffpe` to allow a higher number of alignment mismatches.

#### BrCa BASIS

The BrCa BASIS alignments were part of the manuscript “*10.1038/ncomms12910*” in which alignment was performed on hg38 with *chrEBV* included, using the following alignment settings:

--outSAMstrandField intronMotif

--outFilterIntronMotifs RemoveNoncanonicalUnannotated

--chimSegmentMin 12

--chimJunctionOverhangMin 12

--alignSJDBoverhangMin 10

--alignMatesGapMax 200000

--alignIntronMax 200000

--outSAMtype BAM Unsorted

--chimOutType WithinBAM

--outSAMunmapped Within

--alignEndsType Local

--twopassMode Basic

--twopass1readsN -1

Starting from these chimeric alignments, the main computational Dr Disco pipeline was performed.
