## Supplementary material for "Detection of fusion transcripts and their genomic breakpoints from RNA sequencing data": Dr. Disco technical specification

Manual and Technical Implementation

### Technical implementation Dr. Disco

#### First step: ‘fix’

##### SA:Z:-tags

The SAM output files for chimeric reads generated by RNA STAR have some small limitations. They lack ‘SA:Z:’-tags and ‘FI:i:’-tags. These tags indicate that the alignment of a read was split in two pieces and in which order the split reads were placed. During development of the Dr. Disco software package, STAR (2.7/2.6 and above) has included SA:Z: tags within the `--chimOutType WithinBAM’ output files. However, in these files spanning reads are not marked as such and thus only split reads can be extracted. Currently, Dr. Disco is only compatible with the Chimeric SAM output.

Further in the pipeline, the added tags will be used to pinpoint directly to the other aligned pieces, without the requirement of using name-sorted bam files. The SA:Z: tags places remaining aligned pieces of the pair in the following order: for split reads, first the other aligned piece of the same mate followed by the other mate. For the spanning read(s), only the mate is described in the SA:Z:-tag.

##### RNEXT & PNEXT field

The RNEXT and PNEXT fields refer to the genomic location where the mate / next read is mapped to. IGV has the function to show a split screen, allowing visualizing the reads mate in the other window. The RNEXT and PNEXT fields were used for defining the split view regions. For spanning reads, both mates are displayed in a separate screen. However, with split reads typically one of the split pieces is linked to the properly aligned mate. This makes it difficult to open both split pieces within a split view. Let’s assume that we have a read A and split read B (split into parts B1 and B2). If you start with read A to open the split view, you will be able to display only 1 of the 2 split reads; either B1 or B2. Only 1 of the parts of B (either B1 or B2) allows opening the split view with A, while the other cannot be used to open a split view. From the user's perspective, it is more convenient that starting with read A, you can split the view to B1 and from B1 split the view further to B2 and from B2 back to A. The fix step thus changes the RNEXT and PNEXT field in such a way that this becomes possible. It also allows the user to navigate more easily with "go to next mate" as it allows viewing all aligned pieces that belong to the reads linked in IGV through a complete circle of references.

##### FI:i:-tags

The FI tag stands for “the index of segment in the template” and is added in the FIX step.

##### Read type as read group

The type of discordant read (split / spanning and direction) is added as read group. It can be used to color a discordant read by its type, by setting the ‘Color alignments by’ to ‘read group’.

#### Second step: “detect”

The fixed bam file provides breakpoints more or less ‘as is’ by the SA-tags. The major difficulty is to distinguish between discordant reads that correspond to a fusion gene or translocation from those that are derived from something else such as mapping artifacts. STAR marks a read discordant when the read cannot originate from a transcript of a classical gene because of inconsistent orientation, distance to its mate or an introduced split. The main goal of the “dr-disco detect” step is to gather multiple reads that belong to the same fusion event. This will increase confidence in determining whether a set of discordant reads represent a junctions derived from a structural variant, circRNAs or not. The data structure in which this information is processed is a breakpoint graph. “Detect” is the computationally intensive step for which the ‘fixed’ chimeric alignment are the input, with putative junctions already annotated within the SA:Z:-tags.

We first introduce the input data before we will dive deeper into the detect-step and the breakpoint graph. Reads are either singleton or paired-end. Singletons are reads of which the mate was excluded from alignment or was not aligned and paired-end are those of which both mates were aligned. Discordant reads are classified in two major types: split reads and spanning reads. Singletons are always split reads while paired-end reads can be both split and spanning. If paired-end reads have a mate that was split and the remaining mate was not split, the remaining mate is called a ‘silent’ mate, which is neglected in fusion analysis. Splitting an alignment is an obvious case for marking a read discordant as it might be split over a genomic or exon-to-exon junction. Spanning reads have no split point but have a too large or too small inner distance or are inverted with respect to each other. Note that any discordant read (singleton or paired-end, split or spanning) can also be a spliced read.

The main data structure of the computational analysis is a graph, representing the junctions derived from discordant reads. From the graph’s perspective, each junction is inserted as an edge between two genomic locations and these locations are represented by nodes. The determination of the breakpoints is different for split and spanning reads. The breakpoints of a split read are determined by the genomic locations of both split points. For spanning reads, the R1-read’s last aligned base and the R2-read’s first aligned base represent the junction. If during insertion of an edge into the graph no identical edge was found, it will be inserted in the graph and labelled with the corresponding subtype. Otherwise, the (split/spanning subtype specific) weight is increased by one. Splice junctions will be inserted as edges into a separate graph. The graph data structure is inefficient for accessing targeted genomic regions, for instance to find closely adjacent edges. For quick genomic coordinate based access, a reference to all edges is added to a genomic interval data structure, implemented with the HTSeq library.

##### Merging edges that originate from the same event

Of all Chimeric reads inserted in the graph, only a fraction belongs to actual structural variants or circRNAs. Of all the distinct breakpoints, many are supported perfectly by multiple reads as well as by the slightly shifted junctions as result of the inner distance (spanning reads). By combining the edges that originate from the same event, they can be filtered or classified with higher confidence later. Typically, split reads are located exactly on a breakpoint while spanning reads are slightly shifted. A spanning read is shifted no more than its inner distance. Based on insert size statistics, we have set the maximum inner distance, a parameter for the algorithm, to 450 bp. Split and spanning reads that originate from the same event correspond to edges that are at genomic scale on both sides in close proximity. In this step, edges corresponding to the same event will be merged to bundle evidence that allows better filtering later on.

Merging split and spanning reads derived from the same junction starts by finding the edge with the largest weight (number of reads merged into the exact same junction). Because split reads are more precise in determining an exact breakpoint, they give edges a +50% increase in weight over spanning reads, only during estimation of the heaviest edge. The next step is to search for other edges that have both their nodes no more than the maximum inner distance away and merge them with the heaviest edge. As result of merging, the total weight of the graph stays identical, while the number of unique edges decreases. This is illustrated in **Figure 1** (middle and bottom); the number of edges decreases from 2 to 1, while the weight stays 3 (2 split, 1 discordant).

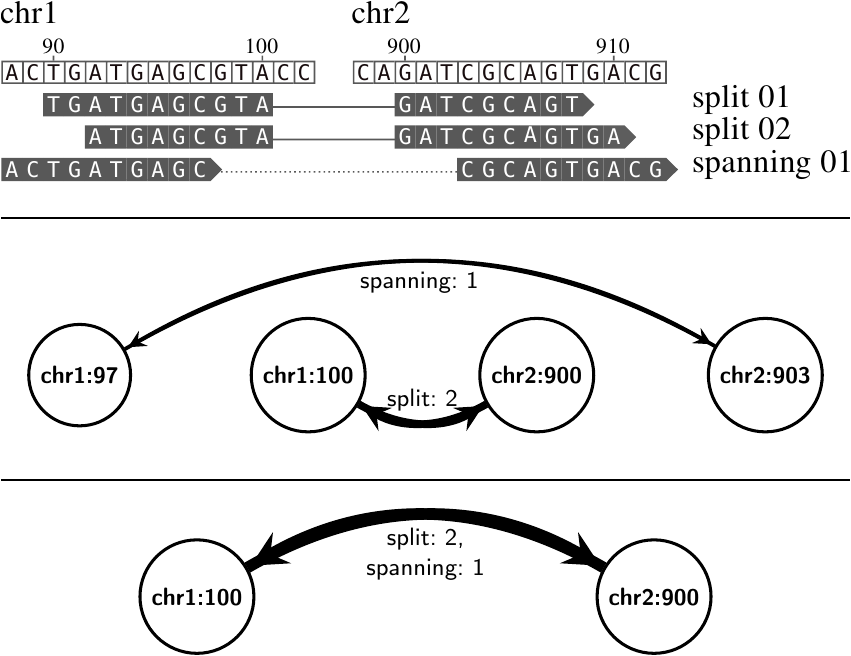

*Figure 1: Merging closely adjacent edges. (****top****) An alignment consisting of two split reads and a spanning read pair that contain two distinct gaps (chr1:100-chr2:900 & chr1:97-chr2:903). (****middle****) After transforming this alignment into a graph, one edge represents two split reads and the other edge represents one discordant mate pair. The weight of the edge that represents two split reads is two and therefore indicated with a thicker line. Both nodes at chr1 and chr2 are only 3 bp away from each other. (****bottom****) As both edges are in close proximity (3 bp + 3 bp), they are presumed to be derived from the same event. After both merging, only one substituted edge will remain of which the weight is the sum of the weight of the edges before merging.*

##### Extract edges from different splice isoforms

In fusion genes with multiple fusion-splice variants, exon-to-exon junctions corresponding to splice variants typically share exons, of which the corresponding edges are overlapping. Based on these overlapping edges, different splice isoforms of the same fusion gene are brought together. In the previous step, edges that belong to the same junction (either exon-to-exon, or DNA breakpoint) were merged, but exon-to-exon junctions from different splice isoforms of the same fusion gene are still separated. This is because introns are typically much larger than the maximum inner distance. Edges that correspond to different splice isoforms of the same fusion gene are extracted in a two-step approach.

First, the shortest possible distance at mRNA level between any two nodes is calculated. These mRNA level distances will be referred to as s-links.

Second, edges that have a distance at the mRNA level of less than the maximum inner distance are extracted from the graph.

Estimation of the distance between two nodes at mRNA level is demonstrated using an example in (**Figure 2**). The gene on chr1 contains two nodes, chr1:1000 and chr1:2500, separated from each other with a genomic distance of 1500 bp. Between these nodes there is a splice junction (chr1:1003-chr1:2490). Splice junctions are determined by the NGS data (CIGAR strings; the N-operations) and not by gene annotations. For each node, there will be searched for the closest splice junction within a genomic distance of 450 bp. Starting with node chr1:1000, there exists one such splice junction at chr1:1003, with a distance of 3 bp. Similarly, for node chr1:2500, there exists one such splice junction, with a distance of 10 bp (to node chr1:2490). The distance at mRNA level between nodes chr1:1000 to chr1:2500 is 3 + 10 = 13 bp and consequently the nodes shall be connected by inserting an edge that represents the distance at mRNA level (s-link).

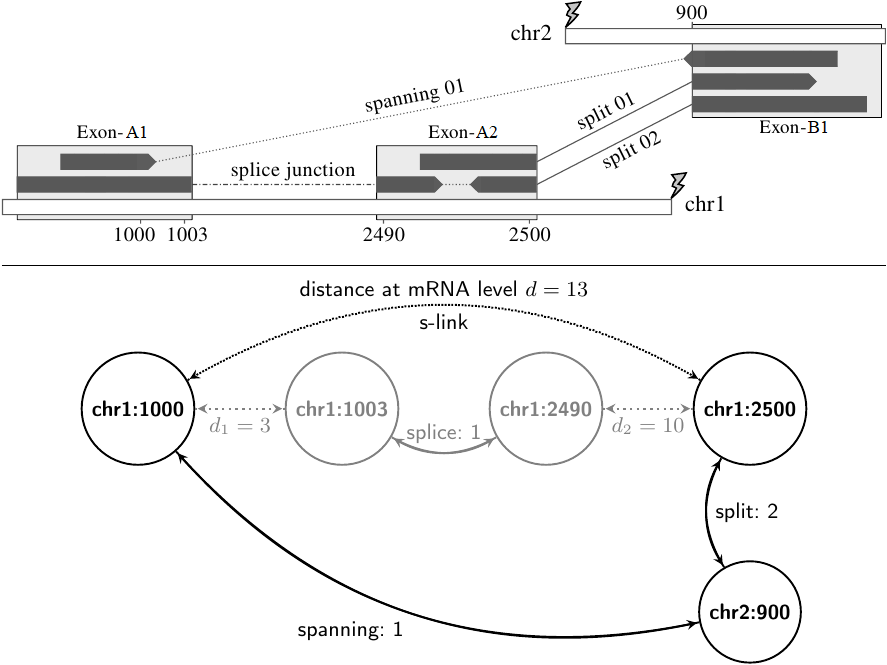

*Figure 2: Extracting edges from different splice isoforms. (top) This schematic view of a fusion gene illustrates how edges corresponding to different splice isoforms are extracted. Exons are indicated as light gray blocks and the junctions within the discordant reads are indicated as lines between the exons. Between the two exons at chr1 and one at chr2 are two split reads (exon-A2 − exon-B1) and a spanning read (exon-A1 − exon-B1). The presence of a splice junction at chr1:1003-2490 indicates these reads belong to the same fusion. Note that no intronic breaks are indicated. (bottom) The total genomic distance between edges chr1:1000-chr2:900 and chr1:2500-chr2:900 is (2500−1000)+(900−900) = 1500 bp. Distances at mRNA level are calculated for each pair of nodes on the same chromosome. These are calculated as sum of the two shortest genomic distances to the nodes that correspond to the splice junction. The distances to the closest splice junctions are d1 = 1003−1000 = 3 and d2 = 2500 − 2490 = 10 bp. The distance at the mRNA level is then calculated as the sum of both distances to the closest splice junction (3 + 10 = 13 bp). If there are two nodes of which the genomic distance minus the splice junction distance is smaller than the maximum inner distance, an s-link is inserted. Although the genomic distance in the example is 1500 bp, the distance at the mRNA level is only 13 bp. Since 13 bp is smaller than the maximum inner distance (450 bp), this allows edges chr1:1000-chr2:900 and chr1:2500-chr2:900, from different isoforms, to be merged.*

Extracting edges that correspond to different splice isoforms starts again with the edge with the highest weight. Considering the example in **Figure 2** (bottom) that contains two edges, the heaviest edge is chr1:2500-chr2:900 with 2 split reads. The distance at mRNA level (s-link) will be traversed separately and recursively, until a maximum cumulative mRNA distance of the inner distance, 450 bp, is reached to store all nodes found during traversal. This selection procedure will look for all nodes within an acceptable mRNA distance, starting with node chr1:2500. This node has only one s-link, with an mRNA distance of 13 bp to node chr1:1000. Because this distance is smaller than 450 bp, the set of node(s) {chr1:2500} gets extended with node chr1:1000 into {chr1:1000,chr1:2500}. Although there are still 437 bases left for further iterations, no other s-links are within the acceptable range surrounding chr1:1000 and the recursion ends. Because there are no s-links connected to node chr2:900, it will return a set only containing itself: {chr2:900}. The merged subgraph will be composed of all existing junctions between both sets of nodes. All edges that exist between the sets of nodes {chr1:1000,chr1:2500} and {chr2:900} are chr1:1000-chr2:900 and chr1:2500-chr2:900. These edges will be extracted as they are likely to be different splice isoforms of the same (fusion) gene structure. Because splice junctions are relatively far apart due to typically large intron sizes, it was decided that such edges will not be merged but will be extracted as a subgraph. These subgraphs preserve multiple exon-to-exon junctions and thus the fusion transcript structure. All edges present in the subgraph are removed from the main graph, so they can only participate in one subgraph, corresponding to one event only. This process is repeated until all edges have been extracted and the main graph is empty.

Sometimes splice sites are not covered by reads, for example when the sequencing depth or coverage near a certain transcript is low. As a result, when spliced reads are absent, extraction of subgraphs based on (splice junction derived) s-links cannot take place. Edges that are located at exons but lack aligned splice junctions will stay separated in the graph, even though they belong to the same (fusion) gene. An example in which this behavior was repeatedly found is alternative exon-0 of TMPRSS2. This is an exon upstream of TMPRSS2, not present in most gene annotations such as RefSeq and UCSC. It was repeatedly observed that spliced reads in this exon were absent and consequently fusion transcripts involving this exon did not get extracted with the rest of the fusion, despite the fact that they are derived from splice isoforms of a TMPRSS2-ERG fusion. In the original setting, this would result in 2 distinct fusions; the TMPRSS2-ERG fusion and a TMPRSS2-exon-0-ERG fusion. To also merge these subgraphs, we added a step that merges based on genomic distance. It compares all extracted subgraphs with all other subgraphs. Given that each edge contains two nodes, we denote for every subgraph two vectors. Each node in an edge is considered left or right, where a left node has a genomic location lexicographically smaller than the right node. Then for every two subgraphs, the smallest genomic distances to a node in the other vector is calculated for both the left and right set of nodes, which is illustrated in **Figure 3**. If the number of nodes in both sets is not identical, the distance to the nodes in the shortest vector are used to determine the distance. Note that genomic distances must be used since splice junctions are missing. For both vectors, a root mean square distance (RMS) is calculated. Using these two vectors (left and right) of minimal genomic distances and the RMS values, it is determined whether two subgraphs will be merged by evaluating:

- If in both vectors there are distances smaller than the maximum inner distance (450 bp).
- If in one vector:
  - 100% of the distances are smaller than the maximum inner distance and of the other vector the RMS is less than 15,000 bp.
  - 70% of the distances are smaller than the maximum inner distance and of the other vector RMS is less than 10,000 bp.
  - 30% of the distances are smaller than the maximum inner distance and of the other vector RMS is less than 5,000 bp.

This fix allowed inclusion of TMPRSS2 exon-0 into the rest of the TMPRSS2-ERG results. After this procedure, each returned sub-graph is a set of edges that can be derived from genomic breaks and from splice junctions of fusion genes. Based on the genomic distance to the closest splice junctions, the SV is classified as either *intronic* or *exonic*. The resulting subgraphs are exported to a list including candidate fusion genes.

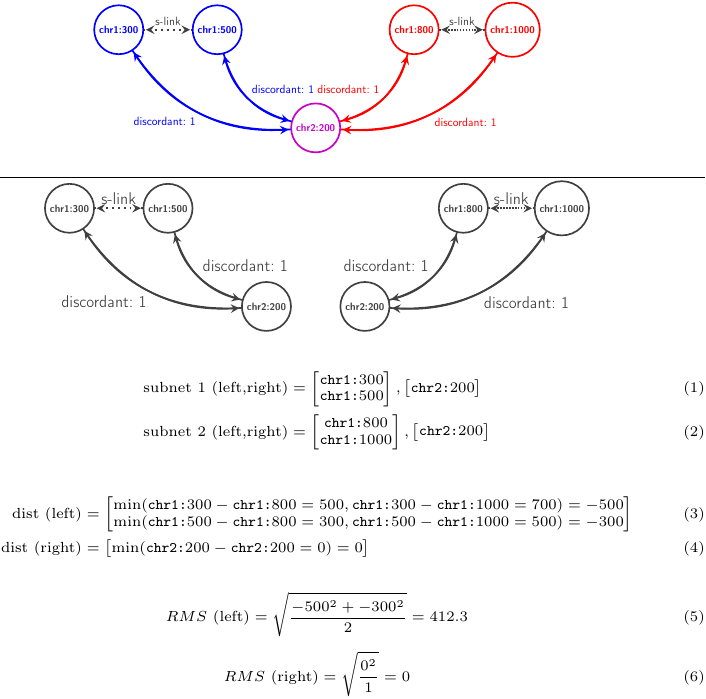
*Figure 3: Extracting edges from different splice isoforms. (top) Two subgraphs (blue and red) that share a common exon are not merged because no splice junction exists between the red and blue nodes. (upper) The two subgraphs completely separated. (mid) To estimate whether both subgraphs can be merged based on genomic distance, vectors containing the corresponding left and right genomic locations are estimated. For subnet 1, the left locations are {chr1:300, chr1:500} and the right locations are {chr2:200}. For subnet 2, the left locations are {chr1:800, chr1:1000} and the right locations are {chr2:200}. (lower) Then the minimum distances between both left and right vectors are estimated. In case the number of nodes in two vectors is not identical, the shortest distances relative to the smallest vector are used and because of this the function becomes symmetrical (d(a, b) == d(b, a)). The estimated minimal distances are: (left): {-500, -300} and (right): {0}. (bottom) Root mean square values of these distances are calculated and are used in combination with the distance vectors to make a decision. Because the calculated RMS values are 412.3 (left) and 0 (right) and both vectors contain distances smaller than 450, these subgraphs can be put together as they are likely to come from the same fusion gene despite no splice junctions were found.*

#### Third: filtering

In the NGS ProToCol PCa dataset, the Dr. Disco Detect step results in approximately 400,000-600,000 subgraph entries, thus candidate events, per sample. Of these, we expect that typically fewer than 100 originate from actual structural variants. Discordant reads do not necessarily originate from fusion genes or structural variants but can also originate from alignment artefacts, sequencing artefacts, non-human contamination or related to population wide variation, read-throughs, and circRNAs. Different filters are applied to erase junctions that do not originate from fusion genes. When there are a high number of identical copies of sequencing reads, the alignment has a rectangular shape. In contrast, when the alignment consists mostly of unique reads, the alignment has on both sides of the junction a more triangular shape.

Recurrent junctions were further analyzed on sequence homology, surrounding sequence complexity and sequencing mismatches. Entries that were considered invalid typically by visual inspection (IGV) contained regional low sequence entropy (poly-A/C/T/G), small repeats, made use of large (>200.000bp) splice junctions to cryptic exons, were circRNAs with back-splice junctions (larger than 200.000 bp), present within annotated rRNA genes, present within UCSC annotated small genomic Simple Repeats. These junctions or surrounding regions that deemed unlikely were added to a blacklist file. A combination of the following features is used for filtering:

(1) the entropy of the start and end positions of the reads that span the breakpoints,

(2) the relation between the number of discordant reads and the number of nodes in the subgraph,

(3) the number of total reads in relation to the subgraph size,

(4) the number of spit and spanning reads,

(5) the slope of the alignment,

(6) symmetry / overhang of alignment between both sides of the breakpoint,

(7) the number of soft-clips near the breakpoints and

(8) the presence of the breakpoint(s) within blacklisted regions.

Precise parameters can be found online at: [https://github.com/yhoogstrate/dr-disco/blob/master/drdisco/DetectOutput.py](https://github.com/yhoogstrate/dr-disco/blob/master/drdisco/DetectOutput.pyf) [function: DetectOutput::classify()]

#### Fourth: integrate with genome and gene reference

In the final, step the returned subgraphs are annotated using a gene annotation and reference sequence. The gene names of remaining candidates are added, and, using a genome reference, also the sequence motifs are determined and compared with canonical splice junction sites. Using the gene annotation, it is determined if there is a potential to produce protein coding transcripts and whether such proteins would be out of frame. If the DNA break (intronic) and fusion transcript (exonic) of the same fusion are found, they are grouped together.
